# Highly diverse mesophotic reef fish communities in Raja Ampat, West Papua

**DOI:** 10.1101/640490

**Authors:** Dominic A. Andradi-Brown, Angela J. E. Beer, Luigi Colin, Hastuti, Catherine E. I. Head, Nur Ismu Hidayat, Steven J Lindfield, Catherine R. Mitchell, Defy N. Pada, Nikola M. Piesinger, Purwanto, Gabby N. Ahmadia

## Abstract

Mesophotic coral ecosystems (MCEs; reefs 30-150 m depth) are poorly studied, with existing research heavily geographically biased away from the most species-rich reef regions. Yet, MCEs are of high interest because of their unique species and potential to act as refuges from the impacts of fishing. Using baited remote underwater video systems, we surveyed reef fish communities from 2 to 85 m depths throughout the Raja Ampat archipelago in West Papua, Indonesia – an area considered the heart of the Coral Triangle where coral reef biodiversity is greatest. We sought to provide the first assessment of fish communities across this depth gradient in the region and identify whether human population density and market access differently affected fish abundance based on depth. Here we show that—similar to shallow reefs—Raja Ampat MCEs are exceptionally diverse, with 152 fish species recorded at depths greater than 40 m. We found that fish community structures were highly depth driven, with declines in fish abundance at increased depth. In contrast to previous studies elsewhere in the world, we found that the proportion of planktivores declined across the shallow reef to MCE depth gradient. While greater human population density correlated with lower Epinephelidae and Lutjanidae abundance (two key fisheries families), we did not find evidence that MCEs provide a depth refuge from fishing. Surprisingly, we also found that fish abundance declined at greater distances from the major regional market – likely caused by historical fisheries pressure in more remote areas. These results both expand upon and contrast some previously established MCE-depth patterns and human impact patterns on fish communities, suggesting that regional context and historical pressures matters. Our findings highlight the need for future MCE studies within the Coral Triangle region.

## Introduction

Mesophotic coral ecosystems (MCEs)—reefs from approximately 30-150 m depth—are of increased conservation and research interest because of their unique species assemblages (Rocha et al. 2018; Pinheiro et al. 2019) and potential to act as refuges from shallow reef impacts (Hinderstein et al. 2010; Loya et al. 2016). MCEs are found in tropical and sub-tropical waters globally, and contain a rich array of biodiversity including corals, fishes, sponges, and algae (Kahng et al. 2010; Baker et al. 2016). MCEs have been poorly documented compared to shallower coral reefs because of sampling limitations at greater depths. With advances in survey technology there has been a greater understanding of MCEs, especially in regions such as the Caribbean, Red Sea, Australia, and Hawaii (Turner et al. 2017; Laverick et al. 2018). The Coral Triangle (comprising: Indonesia, Malaysia, Papua New Guinea, the Philippines, Timor Leste, and the Solomon Islands) hosts the greatest global shallow reef biodiversity (Veron et al. 2009). Yet—despite this diversity—there is comparatively little research effort in the Coral Triangle (Fisher et al. 2011) especially for MCEs (Turner et al. 2017; Laverick et al. 2018). Coral Triangle MCE surveys have been mostly limited to several sites in the Philippines, where highly diverse MCEs have been identified (Cabaitan et al. 2019; Pinheiro et al. 2019).

Based on ecological community composition, MCEs can be divided into upper-MCEs (approximately 30-60 m depth) and lower-MCEs (approximately 60-150 m depth) (Loya et al. 2016). Coral reef fish species richness, abundance, and biomass has broadly been reported to decline across the depth gradient from shallow reefs to MCEs (Kahng et al. 2010; 2014). While reef fish communities are variable between geographical locations, some patterns have begun to emerge. For example, habitat complexity is an important predictor of fish abundance and biomass on MCEs (Boland and Parrish 2005; Brokovich et al. 2008; Bryan et al. 2013). Herbivorous fish—though present in upper-MCEs—often decline in abundance with increased depth and can be near-absent from lower-MCEs (Thresher and Colin 1986; Brokovich et al. 2008; Rocha et al. 2018). While planktivores, in contrast, often increase as a proportion of the fish community on deeper reefs (Thresher and Colin 1986; Brokovich et al. 2008; Rocha et al. 2018).

MCEs may act as fish refuges because shallow reefs maybe more exposed to habitat damage and overfishing, or MCEs may be located at greater distances from shore and so be subject to fewer human impacts (Bridge et al. 2013). Greater biomass of fisheries target species has been observed on MCEs compared to adjacent shallow reefs in many locations (Bejarano et al. 2014; Lindfield et al. 2016; Andradi-Brown et al. 2017). This supports the idea that some MCEs may be exposed to lower fisheries pressures than adjacent shallow reefs. Many small-scale fisheries gears are highly depth specific, with differential impacts across small depth gradients (Ashworth and Ormond 2005; Tyler et al. 2009; Goetze et al. 2011). Small-scale fisheries gears that reach MCEs require additional cost and effort compared to those used in the shallows (Wood et al. 2006; Lindfield et al. 2014). Yet, in some regions MCEs may support important fisheries – for example, in Papua New Guinea mesophotic species comprise large a proportion of some subsistence fish catches (Longenecker et al. 2017; 2019). Testing potential MCE refuge effects is challenging, however, as fish abundance, biomass, and community composition transitions with depth (Thresher and Colin 1986; Brokovich et al. 2008; Rocha et al. 2018; Pinheiro et al. 2019) and with benthic composition and structural complexity (Boland and Parrish 2005; Gratwicke and Speight 2005; Bridge et al. 2012). Despite some potential protection by virtue of their depth, MCEs are still vulnerable to many human impacts including some fisheries, sedimentation, and global stressors such as coral bleaching and increased tropical storm frequency and intensity (Andradi-Brown et al. 2016a; Rocha et al. 2018). With few—though increasing—MCE studies in the Coral Triangle, human impacts on Coral Triangle MCEs remain poorly understood.

Raja Ampat, located in West Papua, Indonesia, and at the center of the Coral Triangle contains extensive reefs, as well as mangrove forests and seagrass beds (Mangubhai et al. 2012). Raja Ampat shallow reefs contain the greatest richness of scleractinian corals (Veron et al. 2009) and fishes (Allen and Erdmann 2009) recorded in a single region – with over 550 scleractinian coral species and over 1,400 fish species (Mangubhai et al. 2012). Given Raja Ampat’s unique diversity, there have been several focused efforts to build regional species lists. For example, Allen and Erdmann (2009) combined detailed field surveys with other records to produce a fish species list for the region. Scleractinian corals have been recorded to 160 m depth within Raja Ampat, though reefs beyond the depths of conventional scuba diving remain poorly characterized (Mangubhai et al. 2012). Unlike many reefs globally, the shallow reefs of Raja Ampat have not faced widespread mortality from bleaching events and shallow reef scleractinian coral cover remained stable between 2010-2016 at approximately 30% (Ahmadia et al. 2017).

Local people living within Raja Ampat have high marine resource dependence, with approximately two-thirds reliant on marine fish for over half their dietary protein needs, and approximately a quarter reliant on marine capture fisheries as their primary occupation (Ahmadia et al. 2017). Gleaning, hand-lines, and spear guns are the dominant fishing gears used by local communities, though commercial trawlers fish in the region (Mangubhai et al. 2012; Ahmadia et al. 2017). Destructive fishing gears such as dynamite and cyanide, while present, are generally less prevalent in Raja Ampat than elsewhere in Indonesia (Mangubhai et al. 2012). Since the 1990s, several non-governmental organizations (NGOs) have worked in Raja Ampat in partnership with local communities and the district and provincial government to establish a marine protected area (MPA) network. These MPAs are zoned based on periodic harvest closure areas, no-take areas, and large sustainable fisheries areas (Grantham et al. 2013), with the intention of allowing local communities to control their marine resources and reduce the number of commercial vessels from elsewhere in Indonesia fishing within Raja Ampat (Mangubhai et al. 2012; Ahmadia et al. 2017). In addition, the entirety of Raja Ampat regency was declared a shark and ray sanctuary in 2012, banning all shark and ray fisheries within the region (Local regulation No. 9, 2012). Despite fisheries impacts, Raja Ampat reefs retain high fish biomass including top predators compared to many reefs elsewhere in the world (Ahmadia et al. 2017). These conservation initiatives—while not specifically targeting MCEs—provide benefits to many MCEs. For example, the Raja Ampat MPA network extends protection beyond shallow coral reefs over many MCE and deep-sea areas. While the shark and ray sanctuary is declared based on the boundary of Raja Ampat Regency—representing a political boundary rather than an ecological boundary—and so encompasses all reefs regardless of depth within Raja Ampat.

In this study, we investigate fish communities on MCEs within Raja Ampat and compare them to adjacent shallow reefs. We specifically identify: (i) whether the high fish species richness that is known to occur on the shallow reefs of Raja Ampat extends into the mesophotic zone; (ii) which fish species are indicative of Raja Ampat MCEs; (iii) patterns in fish species richness, abundance, and trophic groups across the depth gradient; and (iv) we consider how human impacts have differentially affected fish abundance on MCEs compared to shallow reefs. Our study represents one of the largest MCE fish community sampling efforts undertaken to date within the Coral Triangle region, and the first within Raja Ampat.

## Material and methods

### Study sites

Surveys were conducted in January–February 2014 by the Unit Pelaksana Teknis Daerah (UPTD; MPA management authority) in Raja Ampat and Conservation International Indonesia staff from the KM Imbekwan research vessel. Two regions within the Raja Ampat MPA network were sampled in this study: Suaka Alam Perairan (SAP; Marine Sanctuary) Kepulauan Waigeo Sebelah Barat and Kawasan Konservasi Perairan Daerah (KKPD; Regional Marine Protected Area) Selat Dampier (Fig. 1). SAP Kepulauan Waigeo Sebelah Barat is 271,630 ha in area and was declared in 2009 (Ministerial Decree Kepmen-KP No. 64, 2009), while KKPD Selat Dampier is 336,200 ha and was declared in 2008 (Local regulation No. 27, 2008). At the time of the surveys, provisional zonation plans were being developed and implemented for both MPAs and there was limited MPA enforcement. Sites for SAP Kepulauan Waigeo Sebelah Barat were within the provisionally designated no-take MPA zones and in areas to the south around Kawe Island (henceforth grouped with the MPA sites as Waigeo). Sites in KKPD Selat Dampier were within the provisionally designated no-take and sustainable fisheries zones (henceforth Dampier) and covering both Batanta (on the south side of the Dampier Strait) and Kri (on the north side of the Dampier Strait). As these MPAs were new, and zonation implementation and MPA patrolling and enforcement were still under development, it is likely that any differences between sites inside the MPA and outside, or between different MPA zones reflect pre-MPA differences in reef condition. In this study we therefore do not compare sites inside and outside the MPAs (Waigeo), or between no-take and sustainable fishing zones (Dampier).

**Figure 1.**
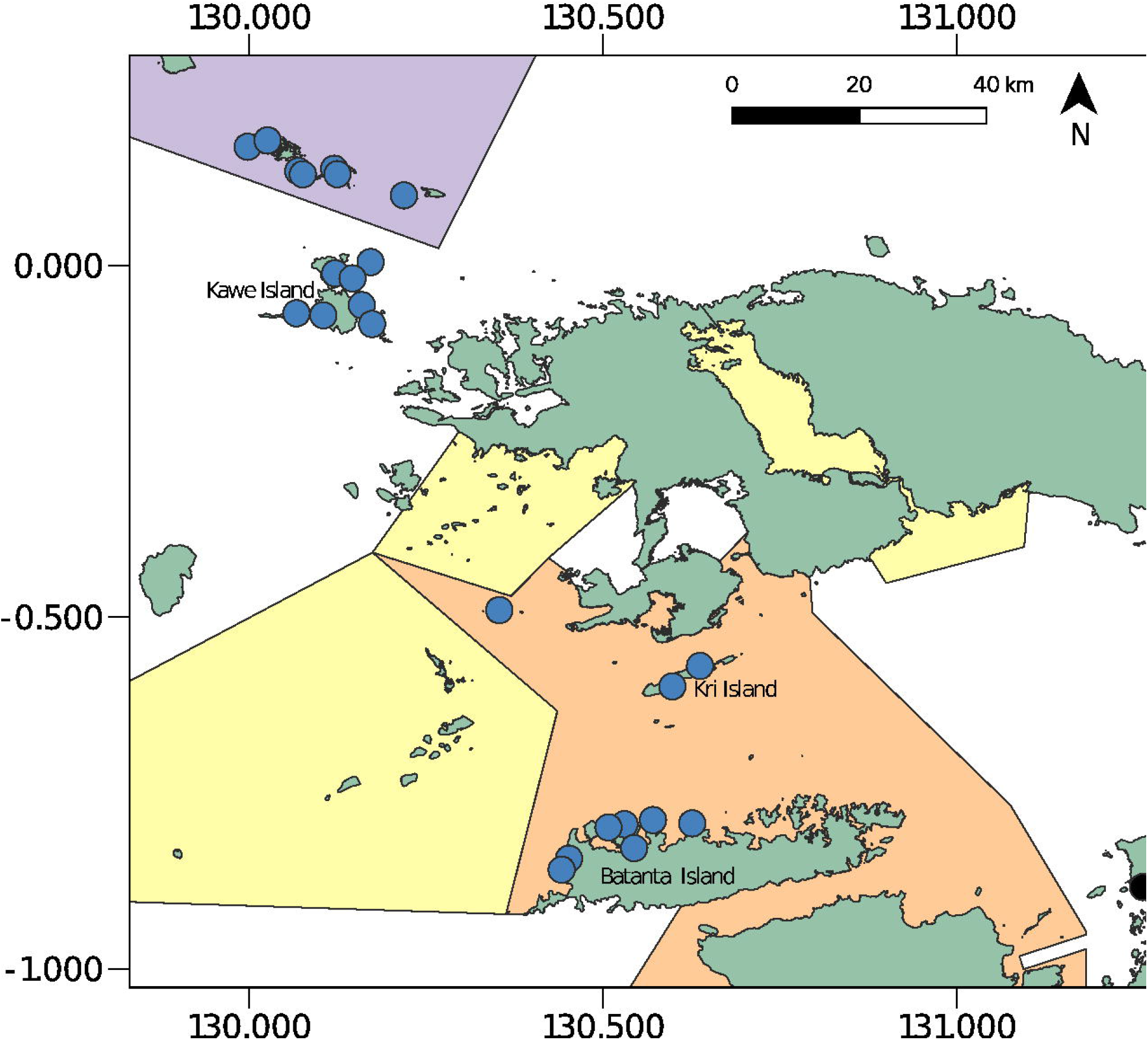
Study sites and marine protected areas within Raja Ampat. Sites were grouped into two regions: Waigeo and Dampier. Waigeo sites included those within SAP Kepulauan Waigeo Sebelah Barat and around Kawe Island. Dampier sites were within KKPD Selat Dampier including both Batanta Island (on the south side of the Dampier Strait) and Kri (on the north side of the Dampier Strait).

Surveys were conducted using baited remote underwater video (BRUV) during daylight hours (07:00-16:00). At each site, surveys were conducted within three depth zones: (i) shallow (2-12 m), (ii) intermediate/mid (15-30 m), and (iii) deep/MCE (40-85 m). A minimum of two BRUV drops were conducted per depth zone at each site – see Electronic Supplementary Material (ESM) Table 1. BRUV replicates within a single site and depth combination were deployed a minimum of 500 m apart. Sites were located a minimum of 1.5 km apart, with 14 sites surveyed within Waigeo, and 10 sites surveyed in Dampier. In total across all sites and depths 160 BRUV drops were completed (see ESM Table 2 for GPS locations). The BRUV consisted of a single GoPro Hero 3 camera, mounted in a frame 0.6 m above the benthos, and recording forward at the BRUV bait arm and seascape. The BRUV bait arm was 1.5 m long and was baited with 0.8-1.0 kg of crushed skipjack tuna. More details on the survey design and methods are available in Beer (2015). BRUVs were left for 60 min on the benthos before being recovered.

**Table 1.** PERMANOVA testing effects of Region, site, benthic principal component 1 (PC1), and depth on fish community structure. Fish community structure is calculated as fourth-root transformed abundance (MaxN) per BRUV drop using Bray-Curtis dissimilarities. Degrees of freedom (DF), sum of squares (SS), and mean square (MS) are shown. Significant p-values at the p<0.05 level are shown in bold.

|  | DF | SS | MS | Pseudo-F | p(perm) |
| --- | --- | --- | --- | --- | --- |
| Region | 1 | 1.09 | 1.09 | 3.91 | **<0.001** |
| Site | 14 | 8.10 | 0.58 | 2.07 | **<0.001** |
| Benthic PC1 | 1 | 3.77 | 3.77 | 13.51 | **<0.001** |
| Depth | 2 | 3.48 | 1.74 | 6.23 | **<0.001** |
| Region:Benthic PC1 | 1 | 0.42 | 0.42 | 1.52 | **0.027** |
| Site:Benthic PC1 | 14 | 6.42 | 0.46 | 1.65 | **<0.001** |
| Region:Depth | 2 | 0.83 | 0.42 | 1.49 | **0.007** |
| Site:Depth | 28 | 11.33 | 0.40 | 1.45 | **<0.001** |
| Benthic PC1:Depth | 2 | 0.53 | 0.26 | 0.95 | 0.591 |
| Region:Benthic PC1:Depth | 2 | 0.35 | 0.17 | 0.63 | 0.999 |
| Site:Benthic PC1:Depth | 28 | 8.17 | 0.29 | 1.05 | 0.169 |
| Residuals | 64 | 17.85 | 0.28 |  |  |
| Total | 159 | 62.34 |  |  |  |

**Table 2.** Dufrëne and Legendre indicator species analysis showing fish species indicative of intermediate and MCE depth zones. Analysis is based on fish abundance with results showing the Dufrëne and Legendre indicator value (measuring the association between the fish species and the depth group) and the associated permutational p-value (based on 99999 permutations). Only fish species with a p<0.05 association for the depth band are shown. See ESM Table 6 for an expanded version of this table including fish species indicative of the shallow depth zone.

| Depth/Family | Species | Indicator Value | p(perm) | Trophic Group |
| --- | --- | --- | --- | --- |
| **Intermediate** |  |  |  |  |
| Acanthuridae | *Acanthurus thompsoni* | 0.22 | 0.008 | Planktivore |
|  | *Naso thynnoides* | 0.32 | <0.001 | Omnivore |
| Balistidae | *Odonus niger* | 0.33 | 0.032 | Planktivore |
|  | *Sufflamen bursa* | 0.39 | 0.001 | Mobile Invert. |
| Caesionidae | *Pterocaesio digramma* | 0.17 | 0.028 | Planktivore |
| Carcharhinidae | *Triaenodon obesus* | 0.18 | 0.042 | Piscivore & Mobile Invert. |
| Chaetodontidae | *Coradion chrysozonus* | 0.22 | 0.033 | Omnivore |
|  | *Forcipiger flavissimus* | 0.17 | 0.030 | Mobile Invert. |
|  | *Forcipiger longirostris* | 0.19 | 0.015 | Mobile Invert. |
| Labridae | *Bodianus diana* | 0.19 | 0.009 | Mobile Invert. |
|  | *Bodianus dictynna* | 0.22 | 0.010 | Mobile Invert. |
|  | *Choerodon zosterophorus* | 0.22 | 0.012 | Mobile Invert. |
|  | *Halichoeres chrysus* | 0.39 | <0.001 | Mobile Invert. |
|  | *Halichoeres prosopeion* | 0.33 | 0.002 | Mobile Invert. |
| Lethrinidae | *Monotaxis grandoculis* | 0.22 | 0.020 | Mobile Invert. |
| Lutjanidae | *Macolor macularis* | 0.29 | 0.004 | Piscivore & Mobile Invert. |
| Monacanthidae | *Paraluteres prionurus* | 0.17 | 0.031 | Omnivore |
| Pomacanthidae | *Centropyge bicolor* | 0.3 | 0.002 | Mobile Invert. |
|  | *Genicanthus lamarck* | 0.29 | 0.001 | Planktivore |
| Pomacentridae | *Pomacentrus amboinensis* | 0.36 | <0.001 | Herbivore & Planktivore |
|  | *Pomacentrus nigromanus* | 0.27 | 0.007 | Planktivore |
| Serranidae | *Plectropomus oligacanthus* | 0.19 | 0.016 | Piscivore & Mobile Invert. |
|  | *Variola albimarginata* | 0.19 | 0.034 | Piscivore |
| Tetraodontidae | *Canthigaster valentini* | 0.17 | 0.031 | Omnivore |
| **MCE** |  |  |  |  |
| Balistidae | *Abalistes stellatus* | 0.41 | <0.001 | Piscivore & Mobile Invert. |
|  | *Sufflamen fraenatum* | 0.23 | 0.023 | Mobile Invert. |
| Carangidae | *Carangoides ferdau* | 0.35 | 0.011 | Piscivore & Mobile Invert. |
| Labridae | *Choerodon jordani* | 0.25 | 0.003 | Mobile Invert. |
| Lethrinidae | *Lethrinus amboinensis* | 0.25 | 0.017 | Piscivore & Mobile Invert. |
|  | *Lethrinus lentjan* | 0.17 | 0.031 | Piscivore & Mobile Invert. |
| Malacanthidae | *Hoplolatilus purpureus* | 0.25 | 0.002 | Planktivore |
| Mullidae | *Parupeneus heptacanthus* | 0.20 | 0.027 | Piscivore & Mobile Invert. |
| Nemipteridae | *Pentapodus nagasakiensis* | 0.36 | 0.004 | Mobile Invert. |
|  | *Pentapodus paradiseus* | 0.39 | <0.001 | Piscivore & Mobile Invert. |
|  | *Scolopsis vosmeri* | 0.25 | 0.003 | Mobile Invert. |
| Pomacentridae | *Pomacentrus nagasakiensis* | 0.17 | 0.031 | Herbivore & Planktivore |

### Video analysis and data processing

Video footage from each BRUV drop was analyzed for all fish species using EventMeasure (SeaGIS, Melbourne, Australia). The first 50 min of footage was analyzed from when the BRUV reached the seabed and stabilized, recording the maximum number (MaxN) for each species. MaxN is defined as the largest number of individuals of a species visible in a single video frame and therefore is a relative measure of fish abundance. MaxN is a widely used metric for static video fish analysis, as it avoids problems of counting individual fish multiple times as they leave and then re-enter the camera field of view (Watson et al. 2010). MaxN has been shown to provide reliable fish community data when compared to other conventional reef survey techniques (e.g. Watson et al. 2010; Andradi-Brown et al. 2016c). Fish identification followed Allen and Erdmann (2012), and all fish were identified to species level or the greatest taxonomic resolution possible (e.g. genus or family) if full species identification was not possible.

We categorized each fish species based on trophic group. All fish data were filtered to remove records of manta and devil rays (Mobulidae), and eagle rays (Myliobatidae), as these are large-bodied transient species that are unlikely to be accurately recorded in our BRUV sampling. Whitetip, blacktip, and grey reef sharks (Carcharhinidae), nurse sharks (Ginglymostomatidae), bamboo sharks (Hemiscylliidae), wedgefishes (Rhinidae), and whiptail rays (Dasyatidae) were retained in our analysis alongside all other reef fishes. Trophic groups were allocated following the Fishbase ‘Food Items’ table, using the Food l-lll hierarchical classification of food items eaten by a species, based on diet composition of >20% of recorded items accessed through rfishbase (Boettiger et al. 2012; Froese and Pauly 2015). Trophic groups were: (i) Herbivore, (ii) Herbivore and Planktivore, (iii) Planktivore, (iv) Planktivore and Mobile Invertebrate Feeder, (v) Omnivore, (vi) Corallivore, (vii) Mobile Invertebrate Feeder, (viii) Piscivore and Mobile Invertebrate Feeder, and (ix) Piscivore. Many studies identifying changes in fish trophic group across depth gradients have occurred in locations with moderate or high fishing pressure – likely influencing recorded patterns. In contrast, Raja Ampat reefs experience lower fishing pressure than many Indo-Pacific reefs (Ahmadia et al. 2017). We therefore also ran our fish trophic comparisons across the depth gradient removing fish species identified as commercially important at the global level to identify how fisheries may alter fish community structure. High commercial value fish species were identified based on the Fishbase commercial value listing, with species listed at ‘high’ or ‘very high’ commercial value included, and sharks of the families Carcharhinidae, Ginglymostomatidae, Hemiscylliidae, and Rhinidae also included.

We explored how broad human and environmental impacts affect reef fish abundance across the depth gradient and region. We used three regionally-relevant reef fish abundance groupings: (i) regionally important fisheries species (Families/Subfamilies: Acanthuridae, Carangidae, Dasyatidae, Epinephelidae, Ginglymostomatidae, Haemulidae, Hemiscylliidae, Lethrinidae, Lutjanidae, Rhinidae, Scarinae, Scombridae, Siganidae, and the species *Cheilinus undulatus)* identified by Ahmadia et al. (2013), (ii) herbivorous fish (Scarinae and Acanthuridae), and (iii) slow-growing high value species (Epinephelidae and Lutjanidae). We sourced three additional human/environmental variables from the Marine Socio-Environmental Covariates database (Yeager et al. 2017) for each BRUV drop. These variables were: (i) mean annual wave exposure (kW/m), (ii) 2015 human population within 10 km, and (iii) linear distance to major regional market – in this case Sorong (Fig. 1) on the West Papuan mainland (km). A 10 km human population buffer was used as studies in the Coral Triangle have suggested that fishers using reef gears typically travel distances <10 km for fishing and generally fish closer to their villages when possible (Soede et al. 2001; Teh et al. 2012).

Fish communities are highly correlated with benthic habitat cover and complexity on both shallow reefs (Gratwicke and Speight 2005) and MCEs (Brokovich et al. 2008; Bryan et al. 2013). To account for varying habitat complexity at each BRUV drop, we visually assessed from the video the Habitat Assessment Score (HAS) following Gratwicke and Speight (2005). HAS is based on six benthic categories: (i) rugosity of substratum, (ii) variety of benthic organism growth forms, (iii) benthic habitat architecture height, (iv) refuge size categories present in the reef, (v) live cover of benthic organisms, and (vi) % hard substratum present (Gratwicke and Speight 2005). Each benthic category is given a value between 1-5, with 5 representing the greatest habitat complexity/structure/cover. All raw data is available from DOI: 10.6084/m9.figshare.8097788.

### Data analysis

To identify differences between individual depth groups within regions for benthic habitat and fish species richness and abundance when plotting, pairwise permutational analysis of variance (permutational-ANOVA) tests (99999 permutations) were used and p-values adjusted with a Bonferroni correction. To visualize changes in benthic HAS and generate a single variable representing benthic habitat we used a Principal Component Analysis (PCA) based on the six benthic HAS categories (each scored between 1-5). PCAs based on the correlation matrix were fitted using the ‘prcomp’ function in R (R Core Team 2013) and visualized with 95% confidence ellipses using the ggbiplot package (Vu 2011).

To visualize differences in fish communities we used non-metric multidimensional scaling (nMDS), based on fourth root transformed fish abundance and fitted using the vegan package (Oksanen et al. 2013). Fourth root transformation was used to down weight the influence of highly abundant species (Anderson et al. 2008). To identify differences in the fish community based on region (Waigeo or Dampier), site, benthic habitat, and depth we used permutational multivariate analysis of variance (PERMANOVA) based on a fourth root transformed Bray-Curtis dissimilarity matrix of fish abundance where each BRUV drop represented a row in the matrix. PERMANOVA was run for 99999 permutations using the ‘adonis’ function from vegan (Oksanen et al. 2013). As we had unbalanced data (i.e. differing number of replicates per site and depth combination), and were testing multiple factors, the interaction sums of squares model terms are not independent (Anderson et al. 2008). We therefore used Type 1—sequential—sums of squares, where each term is fitted after taking account of previous fitted terms. While this means that the order in which terms are fitted can affect our results, this is considered appropriate for nested experimental designs where model terms exhibit a natural hierarchical ordering (Anderson et al. 2008). We fitted PERMANOVAs in the form of Region * Site * Benthic PC1 * Depth. Where ‘*’ means the separate fixed effect and interaction between the variables. Region represents whether the sites were in the Waigeo or Dampier. Benthic PC1 represents the first principal component axis of a PCA of the benthic data. This was used to combine the six HAS benthic categories into a single explanatory variable, as the six categories exhibited high collinearity.

To identify fish species indicative of different depths, we used Dufrêne and Legendre indicator species analysis (Dufrêne and Legendre 1997) on the fish abundance data across the three depth zones. Dufrêne and Legendre indicator species analysis identifies the association between fish species and a particular depth zone. Dufrêne and Legendre indicator species analysis was conducted and permutational p-values were generated using the ‘indval’ function from the labdsv package (Roberts 2013).

To identify how fish community trophic structure changed across the depth gradient we summed fish abundance per trophic group per BRUV (sum of MaxN for each individual species within the trophic group) for each depth zone at each site, and then conducted permutational-ANOVA to test for differences with depth. Depth was treated as a factor: shallow, mid, or deep (MCE). For trophic groups with significant effects of depth on abundance, we identified potential fish species that could be driving these by using SIMPER analysis (Clarke and Warwick 2001). Given that we were interested in identifying species responsible for already identified changes in fish trophic group abundance, we used untransformed species abundance data for the SIMPER analysis, so as not to downweigh the influence of more abundant species.

To identify how fish abundance across the depth gradient was affected by human and environmental conditions we used linear mixed-effect models. These models used fourth root transformed total fish abundance per BRUV as the response variable and included Site as a random factor. As we were specifically interested in whether the human and environmental variables differently affected fish abundance based on depth, and to avoid model overparameterization (Harrison et al. 2018), we limited the global model to only examine each variable and their interactions with depth. Models initially had the form: fish abundance^(1/4)^ ~ Region * Depth + Benthic PC1 * Depth + Wave exposure * Depth + Human population * Depth + Distance to market * Depth + (11 Site). All continuous explanatory variables were scaled and centered prior to model fitting. We first fit this full model, but then simplified using Akaike Information Criterion with a correction for small samples (AICc) to remove non-significant interactions. Models were fit using the ‘lmer’ function from the lmerTest package (Kuznetsova et al. 2017), which builds on the lme4 package (Bates et al. 2015) in R. We used Satterthwaite approximations to generate degrees of freedom and p-values for the final reported model – which is a preferred method for evaluating significance from mixed-effects models because of low Type I error rates (Luke 2017). We used partial effects plots to visualize the effects of human population density and distance to market on fish abundance based on the remef package (Hohenstein and Kliegl 2019).

## Results

### Benthic habitat complexity

Habitat complexity declined as depth increased for both Dampier and Waigeo (Fig. 2A). Mean total HAS per BRUV declined from 19 ± 1 (mean ± SE) on shallow reefs to 9 ± 1 on MCEs in Dampier (shallow vs. MCE pairwise comparison: F_1,18_ = 43, adjusted-p = <0.001) and from 19 ± 1 on shallow reefs to 10 ± 1 on MCEs in Waigeo (shallow vs. MCE pairwise comparison: F_1,26_ = 70, adjusted-p = < 0.001). All six HAS categories (rugosity, variety of growth forms, height, refuge size category, live cover, and hard substratum) strongly correlate with the first principal component axis (explaining 80% of benthic variation observed), which shows clear depth groupings (Fig. 3A).

**Figure 2.**
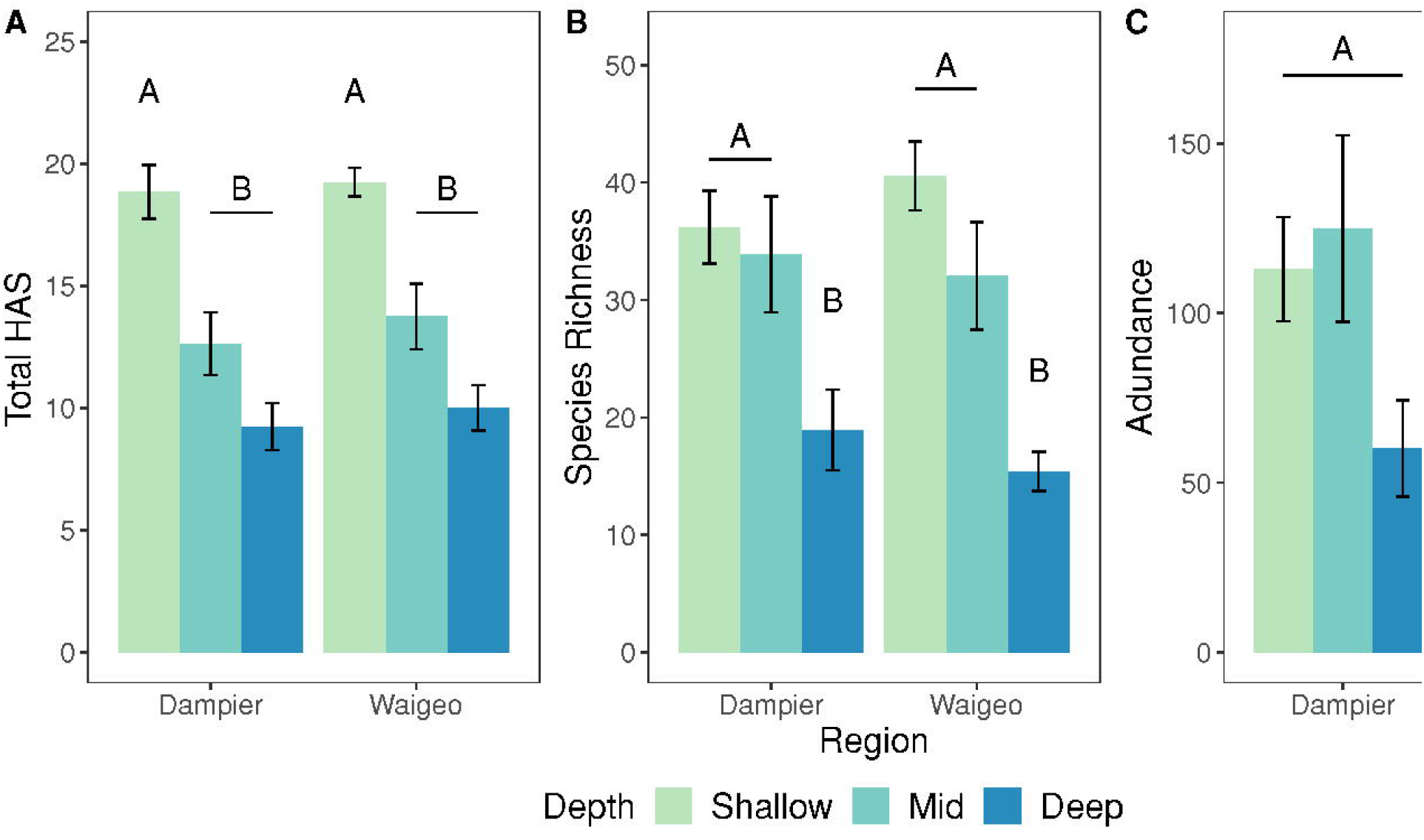
Overall benthic and fish community differences with depth. (A) Habitat assessment score (HAS), (B) fish species richness, and (C) fish abundance. (A) shows the mean total HAS per BRUV drop, with values per BRUV potentially ranging between 6-30 based on scoring a value of 1-5 for each of: rugosity, variety of growth forms, height, refuge size category, live cover, and hard substratum. Abundance represents the mean number of individual fish per BRUV (ind./BRUV), based on summing the MaxN for each species per BRUV. Significant pairwise differences (p<0.05) between depth groups are indicated by letter groupings. Error bars show 1 standard error above and below the mean.

**Figure 3.**
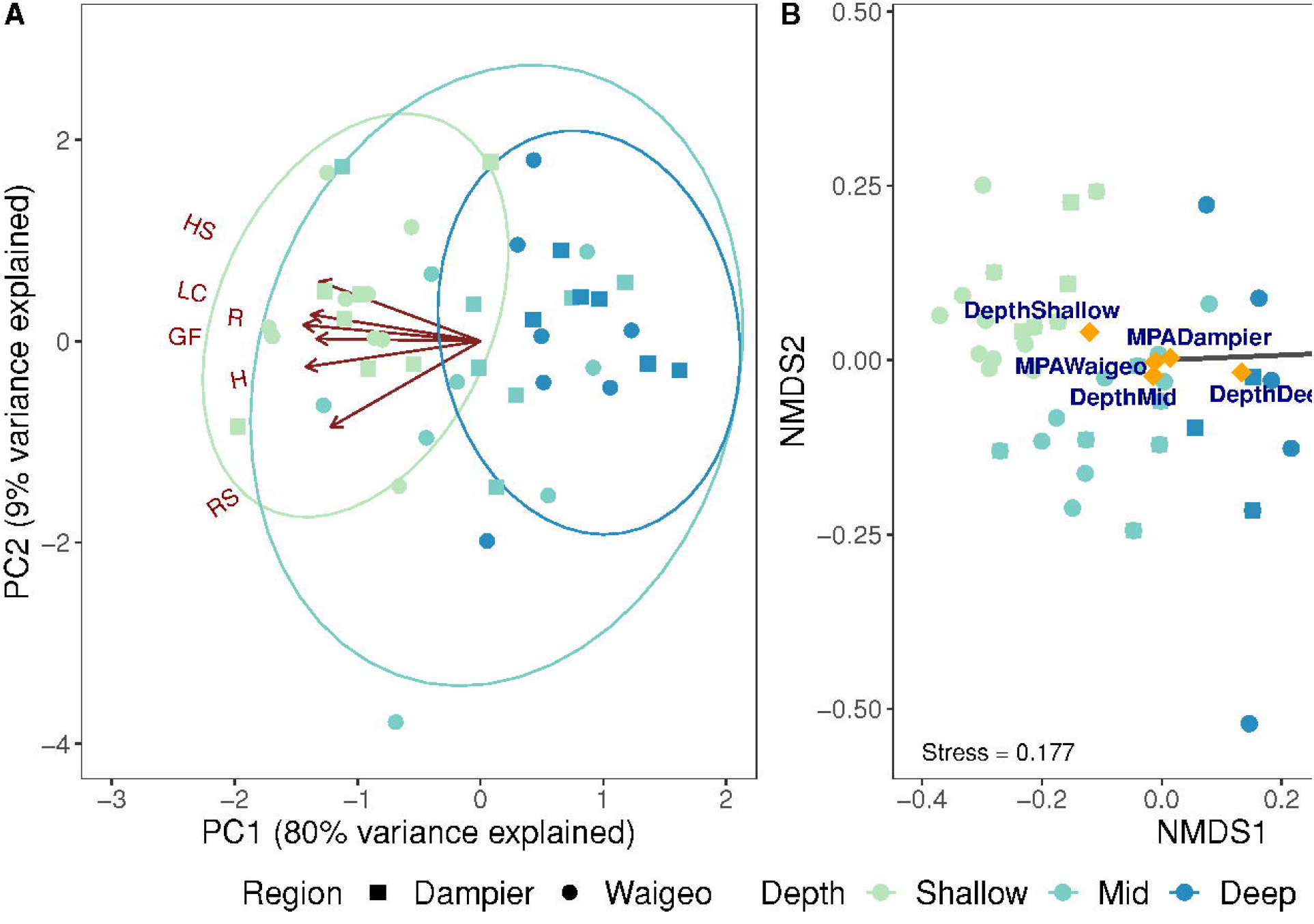
Patterns in benthic and fish community structure based on depth and region. (A) PCA of benthic communities based on mean HAS per depth at each site. 95% confidence ellipses are shown. (B) Fish community nMDS based on fourth root transformed abundance. Correlations with continuous variables (benthic PC1) and averages for factor levels (Region and Depth) are shown.

### Fish species richness and abundance

Overall, we recorded 521 fish species across the two regions and all depth zones, with 328, 307, and 152 fish species recorded on shallow, intermediate, and mesophotic reefs, respectively. Our surveys identified 31 species not previously recorded by the Allen and Erdmann (2009) fish species list for Raja Ampat (ESM Table 3). Of these, 13 were potential new records for the region, and 18 had previously been recorded from the adjacent West Papuan reefs of Cenderawasih Bay, Fakfak, or Kaimana (see Allen and Erdmann 2009). We found 99 fish species at greater depths than their maximum depth listed on Fishbase, of which 76 were extensions onto MCEs in the 40-85 m depth range, and 23 were extensions on shallow or intermediate depth reefs (<40 m maximum depth; ESM Table 4). Species with depth extensions included commercially important families such as emperor (Lethrinidae – nine species with depth extensions) and snapper (Lutjanidae – one species). Some important fisheries species were also exclusively recorded on MCEs (ESM Table 5) – for example four emperor species *(Gymnocranius elongatus, Lethrinus laticaudis, Lethrinus lentjan,* and *Lethrinus rubrioperculatus)* and one snapper *(Lutjanus argentimaculatus). We* recorded International Union for Conservation of Nature (IUCN) red listed species on MCEs in Raja Ampat, such as the endangered humphead wrasse *(Cheilinus undulatus)* and the vulnerable green bumphead parrotfish *(Bolbometopon muricatum).* Humphead wrasse were found across all depths at a similar density (ESM Table 5), but bumphead parrotfish were only recorded on intermediate depth reefs and MCEs – where they were observed to 41 m maximum depth (ESM Table 4; ESM Table 5).

**Table 3.** Fish trophic group species richness and comparison of abundance across the depth gradient. Richness represents the number of unique species recorded for a trophic grouping within a depth zone. Abundance represents the mean total abundance per BRUV drop within a depth zone based on summing species level MaxN for all species within a tropic group per BRUV drop. Changes in abundance across the depth gradient were tested with a permutational-ANOVA of the form Trophic Group Abundance ~ Region + Site + Benthic PC1 + Depth. As permutational-ANOVA uses sequential sums of squares, the abundance depth effect represent the effect of depth on trophic group abundance after controlling for the effect of region, site, and benthic habitat variation. Significant p-values at the p<0.05 level are shown in bold. Abundance values represent MaxN ± Standard Error.

| Trophic Group | Shallow |  | Intermediate |  | MCE |  | Abundance depth effect |  |
| --- | --- | --- | --- | --- | --- | --- | --- | --- |
|  | Richness | Abundance (MaxN/BRUV) | Richness | Abundance (MaxN/BRUV) | Richness | Abundance (MaxN/BRUV) | Pseudo-F | p(perm) |
| Herbivore | 50 | 12.92 ± 1.91 | 38 | 7.26 ± 0.84 | 9 | 2.35 ± 0.52 | 3.75 | **0.026** |
| Herbivore & Planktivore | 36 | 17.61 ± 3.65 | 35 | 14.56 ± 2.89 | 10 | 9.39 ± 4.11 | 1.02 | 0.367 |
| Planktivore | 56 | 46.06 ± 7.19 | 59 | 46.35 ± 9.38 | 26 | 8.62 ± 1.9 | 4.27 | **0.016** |
| Planktivore & Mobile Invertebrate | 11 | 2.88 ± 0.62 | 14 | 4.77 ± 1.56 | 7 | 2.5 ± 0.97 | 0.79 | 0.513 |
| Omnivore | 43 | 10.67 ± 3.14 | 29 | 7.05 ± 1.24 | 18 | 3.41 ± 0.63 | 2.14 | 0.110 |
| Corallivore | 22 | 6.14 ± 0.53 | 20 | 2.98 ± 0.39 | 7 | 2.42 ± 0.38 | 5.90 | **0.004** |
| Mobile Invertebrate | 94 | 22.92 ± 1.49 | 97 | 27.64 ± 9.24 | 67 | 17.97 ± 3.27 | 0.52 | 0.662 |
| Piscivore & Mobile Invertebrate | 95 | 11.83 ± 0.83 | 95 | 16.21 ± 1.65 | 66 | 17.72 ± 3.04 | 4.66 | **0.010** |
| Piscivore | 16 | 4.52 ± 1.45 | 18 | 3.95 ± 1.05 | 10 | 1.75 ± 0.3 | 2.82 | 0.057 |

**Table 4.**
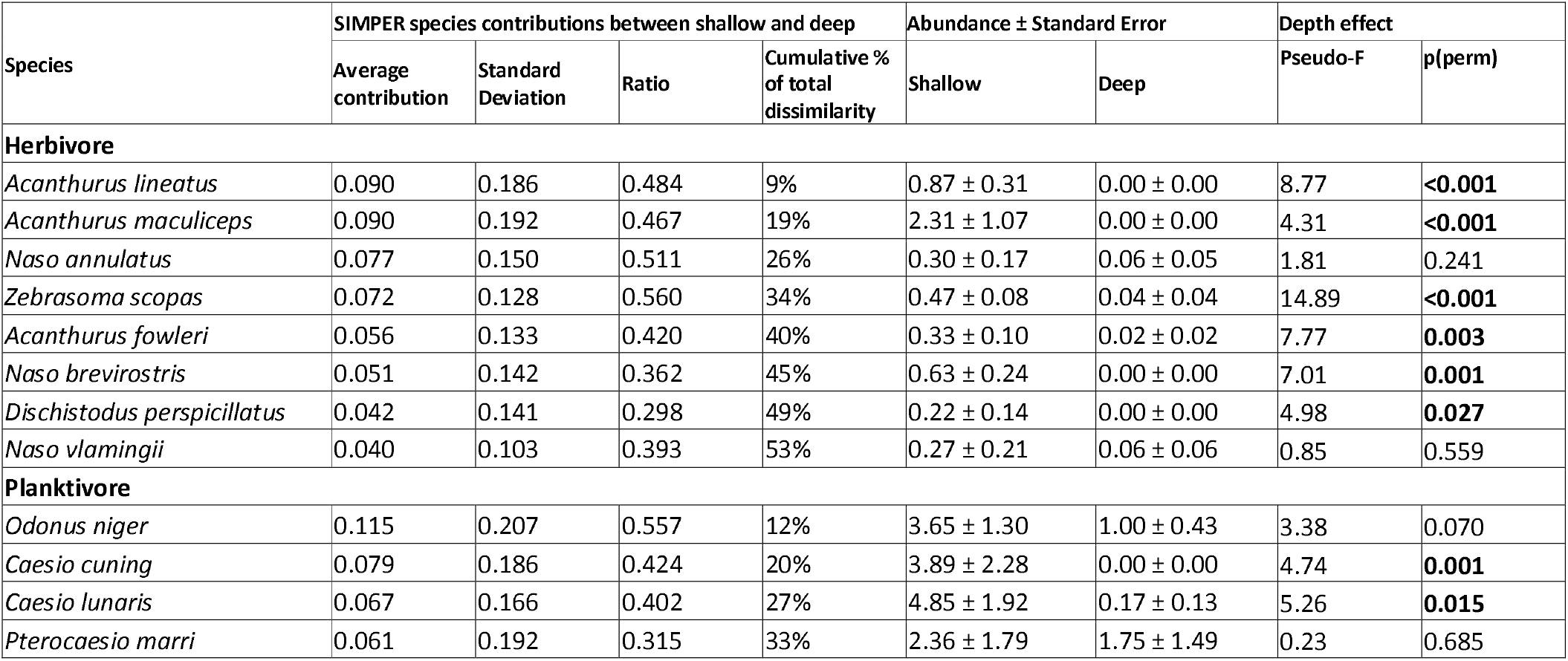
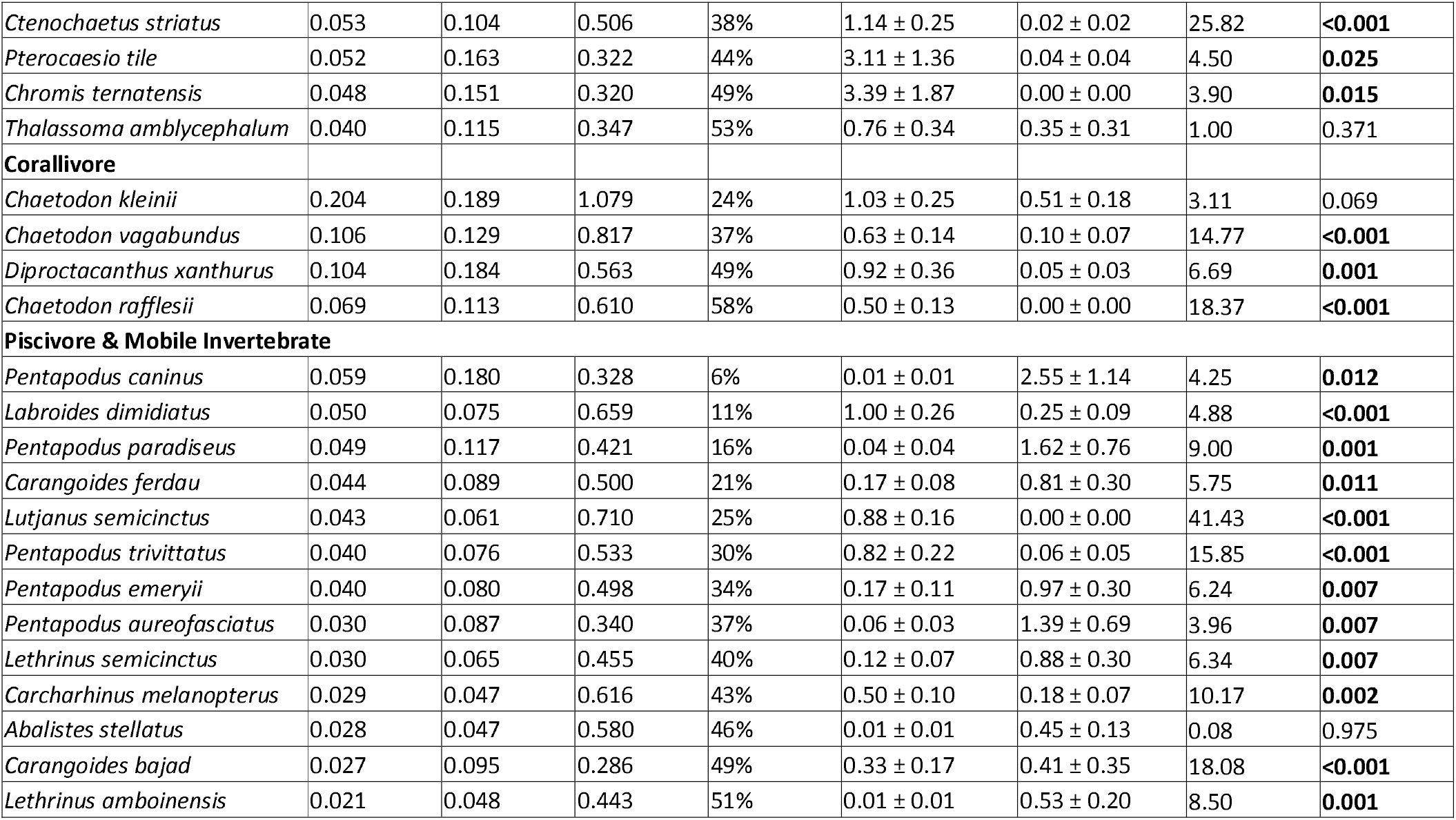
Fish species potentially driving changes in herbivore, planktivore, corallivore, and piscivore and mobile invertebrate feeder abundance between shallow and deep reefs (MCEs). SIMPER analysis based on Bray-Curtis dissimilarities from fish abundance data analyzed by trophic group. Contributions from each species to average dissimilarity when comparing shallow reef versus deep reef fish communities. Standard deviation for contributions, and the ration between species average contribution and standard deviation is also shown. Species are ordered by average contribution to dissimilarity between shallow and deep reefs, represent the most influential and cumulatively responsible for >50% of the total dissimilarity. Total number of species recorded on shallow and deep reefs for each group were: herbivores – 56, planktivores – 82, corallivores – 24, piscivores and mobile invertebrate feeders – 127. Abundance represents the mean MaxN per BRUV drop. Changes in abundance across the depth gradient were tested with a permutational-ANOVA of the form MaxN ~ Depth. Only shallow and deep reef data was used in this analysis. Significant p-values at the p<0.05 level are shown in bold.

| Species | SIMPER species contributions between shallow and deep |  |  |  | Abundance ± Standard Error |  | Depth effect |  |
| --- | --- | --- | --- | --- | --- | --- | --- | --- |
|  | Average contribution | Standard Deviation | Ratio | Cumulative % of total dissimilarity | Shallow | Deep | Pseudo-F | p(perm) |
| **Herbivore** |  |  |  |  |  |  |  |  |
| *Acanthurus lineatus* | 0.090 | 0.186 | 0.484 | 9% | 0.87 ± 0.31 | 0.00 ± 0.00 | 8.77 | **<0.001** |
| *Acanthurus maculiceps* | 0.090 | 0.192 | 0.467 | 19% | 2.31 ± 1.07 | 0.00 ± 0.00 | 4.31 | **<0.001** |
| *Naso annulatus* | 0.077 | 0.150 | 0.511 | 26% | 0.30 ± 0.17 | 0.06 ± 0.05 | 1.81 | 0.241 |
| *Zebrasoma scopas* | 0.072 | 0.128 | 0.560 | 34% | 0.47 ± 0.08 | 0.04 ± 0.04 | 14.89 | **<0.001** |
| *Acanthurus fowleri* | 0.056 | 0.133 | 0.420 | 40% | 0.33 ± 0.10 | 0.02 ± 0.02 | 7.77 | **0.003** |
| *Naso brevirostris* | 0.051 | 0.142 | 0.362 | 45% | 0.63 ± 0.24 | 0.00 ± 0.00 | 7.01 | **0.001** |
| *Dischistodus perspicillatus* | 0.042 | 0.141 | 0.298 | 49% | 0.22 ± 0.14 | 0.00 ± 0.00 | 4.98 | **0.027** |
| *Naso vlamingii* | 0.040 | 0.103 | 0.393 | 53% | 0.27 ± 0.21 | 0.06 ± 0.06 | 0.85 | 0.559 |
| **Planktivore** |  |  |  |  |  |  |  |  |
| *Odonus niger* | 0.115 | 0.207 | 0.557 | 12% | 3.65 ± 1.30 | 1.00 ± 0.43 | 3.38 | 0.070 |
| *Caesio cuning* | 0.079 | 0.186 | 0.424 | 20% | 3.89 ± 2.28 | 0.00 ± 0.00 | 4.74 | **0.001** |
| *Caesio lunaris* | 0.067 | 0.166 | 0.402 | 27% | 4.85 ± 1.92 | 0.17 ± 0.13 | 5.26 | **0.015** |
| *Pterocaesio marri* | 0.061 | 0.192 | 0.315 | 33% | 2.36 ± 1.79 | 1.75 ± 1.49 | 0.23 | 0.685 |
| *Ctenochaetus striatus* | 0.053 | 0.104 | 0.506 | 38% | 1.14 ± 0.25 | 0.02 ± 0.02 | 25.82 | **<0.001** |
| *Pterocaesio tile* | 0.052 | 0.163 | 0.322 | 44% | 3.11 ± 1.36 | 0.04 ± 0.04 | 4.50 | **0.025** |
| *Chromis ternatensis* | 0.048 | 0.151 | 0.320 | 49% | 3.39 ± 1.87 | 0.00 ± 0.00 | 3.90 | **0.015** |
| *Thalassoma amblycephalum* | 0.040 | 0.115 | 0.347 | 53% | 0.76 ± 0.34 | 0.35 ± 0.31 | 1.00 | 0.371 |
| **Corallivore** |  |  |  |  |  |  |  |  |
| *Chaetodon kleinii* | 0.204 | 0.189 | 1.079 | 24% | 1.03 ± 0.25 | 0.51 ± 0.18 | 3.11 | 0.069 |
| *Chaetodon vagabundus* | 0.106 | 0.129 | 0.817 | 37% | 0.63 ± 0.14 | 0.10 ± 0.07 | 14.77 | **<0.001** |
| *Diproctacanthus xanthurus* | 0.104 | 0.184 | 0.563 | 49% | 0.92 ± 0.36 | 0.05 ± 0.03 | 6.69 | **0.001** |
| *Chaetodon rafflesii* | 0.069 | 0.113 | 0.610 | 58% | 0.50 ± 0.13 | 0.00 ± 0.00 | 18.37 | **<0.001** |
| **Piscivore & Mobile Invertebrate** |  |  |  |  |  |  |  |  |
| *Pentapodus caninus* | 0.059 | 0.180 | 0.328 | 6% | 0.01 ± 0.01 | 2.55 ± 1.14 | 4.25 | **0.012** |
| *Labroides dimidiatus* | 0.050 | 0.075 | 0.659 | 11% | 1.00 ± 0.26 | 0.25 ± 0.09 | 4.88 | **<0.001** |
| *Pentapodus paradiseus* | 0.049 | 0.117 | 0.421 | 16% | 0.04 ± 0.04 | 1.62 ± 0.76 | 9.00 | **0.001** |
| *Carangoides ferdau* | 0.044 | 0.089 | 0.500 | 21% | 0.17 ± 0.08 | 0.81 ± 0.30 | 5.75 | **0.011** |
| *Lutjanus semicinctus* | 0.043 | 0.061 | 0.710 | 25% | 0.88 ± 0.16 | 0.00 ± 0.00 | 41.43 | **<0.001** |
| *Pentapodus trivittatus* | 0.040 | 0.076 | 0.533 | 30% | 0.82 ± 0.22 | 0.06 ± 0.05 | 15.85 | **<0.001** |
| *Pentapodus emeryii* | 0.040 | 0.080 | 0.498 | 34% | 0.17 ± 0.11 | 0.97 ± 0.30 | 6.24 | **0.007** |
| *Pentapodus aureofasciatus* | 0.030 | 0.087 | 0.340 | 37% | 0.06 ± 0.03 | 1.39 ± 0.69 | 3.96 | **0.007** |
| *Lethrinus semicinctus* | 0.030 | 0.065 | 0.455 | 40% | 0.12 ± 0.07 | 0.88 ± 0.30 | 6.34 | **0.007** |
| *Carcharhinus melanopterus* | 0.029 | 0.047 | 0.616 | 43% | 0.50 ± 0.10 | 0.18 ± 0.07 | 10.17 | **0.002** |
| *Abalistes stellatus* | 0.028 | 0.047 | 0.580 | 46% | 0.01 ± 0.01 | 0.45 ± 0.13 | 0.08 | 0.975 |
| *Carangoides bajad* | 0.027 | 0.095 | 0.286 | 49% | 0.33 ± 0.17 | 0.41 ± 0.35 | 18.08 | **<0.001** |
| *Lethrinus amboinensis* | 0.021 | 0.048 | 0.443 | 51% | 0.01 ± 0.01 | 0.53 ± 0.20 | 8.50 | **0.001** |

**Table 5.** Linear mixed-effect model of the effects of depth, benthic cover, wave exposure, human population within 10 km, and the distance to major market on fourth root transformed fish abundance. Fish abundance was measured as the summed MaxN values for each species within the fish group, and was fourth root transformed prior to analysis. Benthic PC1 indicates the first principal components axis from a PCA of each component of the benthic HAS per BRUV drop.

| Model Parameter | Estimate | SE | df | t-value | p |
| --- | --- | --- | --- | --- | --- |
| **Regional fisheries species** |  |  |  |  |  |
| Intercept | 1.73 | 0.27 | 30.85 | 6.49 | **<0.001** |
| Population density | -0.15 | 0.08 | 26.46 | -1.89 | 0.070 |
| Depth Mid | -0.14 | 0.13 | 146.94 | -1.15 | 0.253 |
| Depth Deep | -0.34 | 0.15 | 151.69 | -2.31 | **0.022** |
| Benthic PC1 | 0.07 | 0.10 | 155.56 | 0.67 | 0.505 |
| Distance | -0.57 | 0.24 | 25.02 | -2.33 | **0.028** |
| MPA Wayag | 0.82 | 0.44 | 24.83 | 1.86 | 0.075 |
| Depth Mid: Benthic PC1 | 0.27 | 0.13 | 155.66 | 2.19 | **0.030** |
| Depth Deep: Benthic PC1 | 0.16 | 0.14 | 153.86 | 1.14 | 0.257 |
| **Scarinae and Acanthuridae** |  |  |  |  |  |
| Intercept | 1.49 | 0.16 | 85.96 | 9.20 | **<0.001** |
| Depth Mid | -0.08 | 0.16 | 148.14 | -0.52 | 0.603 |
| Depth Deep | -1.01 | 0.19 | 152.70 | -5.40 | **<0.001** |
| Benthic PC1 | 0.30 | 0.13 | 156.00 | 2.23 | **0.027** |
| MPA Wayag | 0.03 | 0.14 | 21.28 | 0.23 | 0.824 |
| Depth Mid: Benthic PC1 | 0.33 | 0.16 | 154.63 | 2.07 | **0.040** |
| Depth Deep: Benthic PC1 | -0.14 | 0.18 | 152.92 | -0.76 | 0.449 |
| **Epinephelidae and Lutjanidae** |  |  |  |  |  |
| Intercept | 0.68 | 0.22 | 27.50 | 3.04 | **0.005** |
| Depth Mid | 0.09 | 0.11 | 147.77 | 0.81 | 0.420 |
| Depth Deep | -0.06 | 0.13 | 154.97 | -0.42 | 0.674 |
| Population density | 0.05 | 0.10 | 102.09 | 0.43 | 0.666 |
| Benthic PC1 | 0.42 | 0.06 | 158.18 | 7.37 | **<0.001** |
| Distance | -0.28 | 0.23 | 34.89 | -1.22 | 0.230 |
| MPA Wayag | 0.78 | 0.39 | 26.07 | 1.99 | 0.057 |
| Depth Mid: Population density | -0.35 | 0.15 | 137.12 | -2.42 | **0.017** |
| Depth Deep: Population density | -0.17 | 0.14 | 136.93 | -1.19 | 0.237 |
| Depth Mid: Distance | -0.34 | 0.15 | 136.55 | -2.30 | **0.023** |
| Depth Deep: Distance | -0.05 | 0.14 | 136.29 | -0.33 | 0.741 |

Fish species richness declined as depth increased similarly for both regions (Fig. 2B). Richness dropped from a mean number of fish species per BRUV drop of 36 ± 3 on shallow reefs to 19 ± 3 on MCEs in Dampier (shallow verses MCE pairwise comparison: F_1,18_ = 55, adjusted-p = 0.006) and 41 ± 3 on shallow reefs to 15 ± 2 on MCEs in Waigeo (shallow verses MCE pairwise comparison: F_1,26_ = 55, adjusted-p = <0.001). Fish abundance also declined with increasing depth in Waigeo from 136 ± 13 ind./BRUV on shallow reefs to 40 ±67 ind./BRUV on MCEs (shallow vs. MCE pairwise comparison: F_1,26_ = 48, adjusted-p = <0.001; Fig. 2C). While mean fish abundance was lower at deeper depths for Dampier (113 ± 15 ind./BRUV on shallow reefs verses 60 ± 14 ind./BRUV on MCEs this did not represent a significant decline (shallow vs. MCE pairwise comparison: F_1,18_ = 6, adjusted-p = 0.062; Fig. 2C) – likely caused by high spatial variation in fish abundance.

Differences in fish communities were identified between the two regions, individual sites, benthic habitat structure, and depth (Table 1). Depth was an important driver of fish communities (Fig. 3B), and indicator species analysis identified 74 species associated with the shallows (ESM Table 6), 24 fish species associated with intermediate depths, and 13 species associated with MCEs (Table 2). Of particular note on MCEs were the presence of three Nemipteridae species *(Pentapodus nagasakiensis, Pentapodus paradiseus,* and *Scolopsis vosmeri),* which were commonly recorded. Purple sand tilefish *(Hoplolatilus purpureus)* were also indicative of MCEs, which given the lower MCE HAS score and that this species associates with sand/rubble areas is unsurprising. Overall, 90 fish species significantly changed in abundance based on depth, with 62 of these fish species at lower abundance on MCEs than shallow reefs, while 18 fish species were at greater abundance on MCEs than shallow reefs (ESM Table 5).

### Fish trophic structure

Mobile invertebrate feeders and joint piscivore and mobile invertebrate feeders comprised the largest component of MCE fish communities, together accounting for 65% (Dampier) and 55% (Waigeo) of the community by abundance (Fig. 4A). The majority (83%) of indicator species associated with MCEs were also mobile invertebrate feeders (both exclusively or combined with piscivory) (Table 2). This MCE community contrasts with the shallows, where herbivorous and planktivorous fishes made up 53% (Dampier) and 57% (Waigeo) of the community abundance (Fig 4A). Shallow reef indicator species were comprised by 14% herbivores, 11% corallivores, and only 19% mobile invertebrate feeders and/or piscivores (ESM Table 6). Removing commercially valuable fish species altered the reef fish community trophic structure considerably (Fig. 4B) by disproportionately removing mobile invertebrate feeders and piscivores. This led to a shallow reef community dominated by herbivores and planktivores, but a more even trophic group distribution on MCEs (Fig 4B).

**Figure 4.**
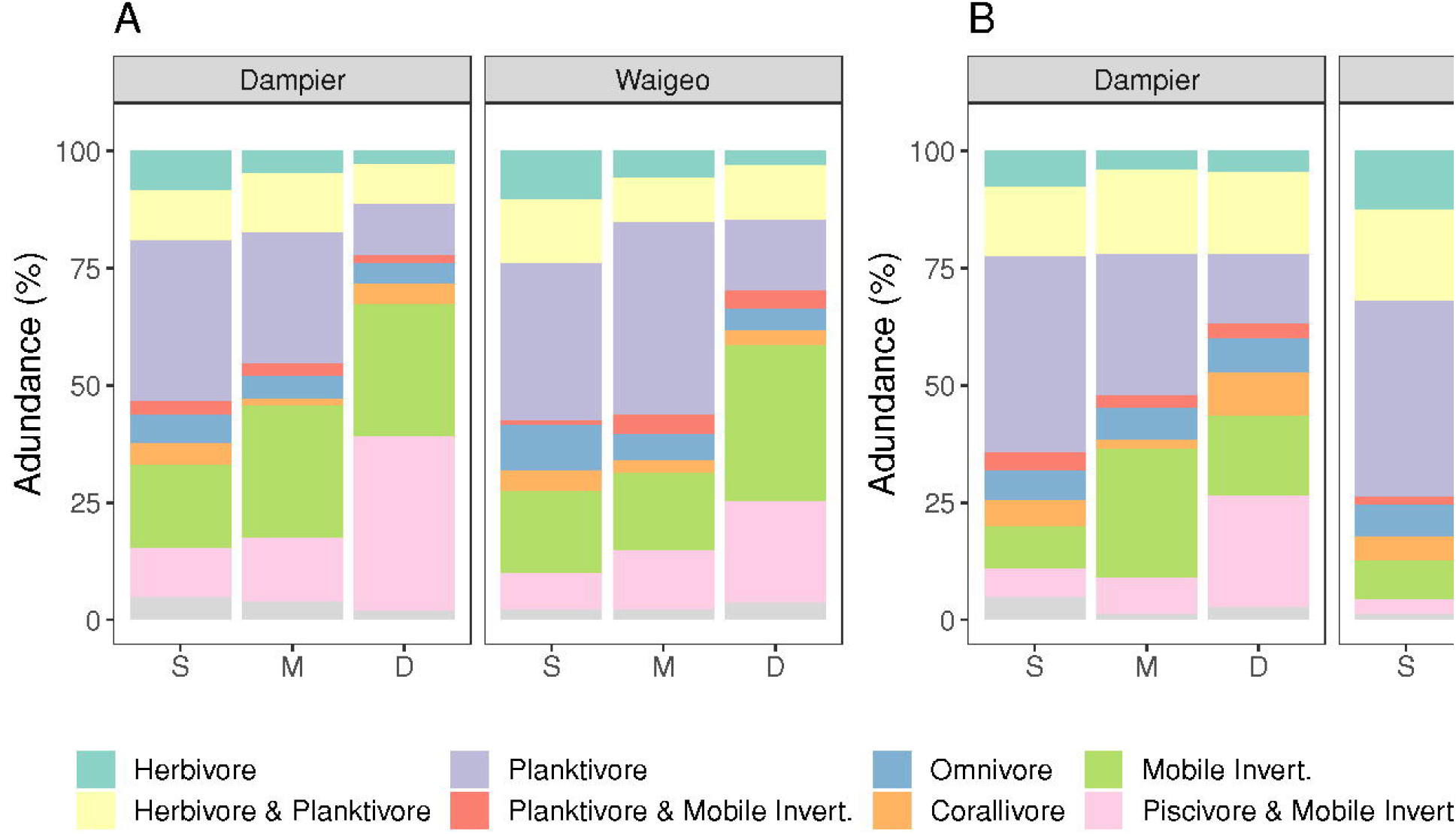
Community composition of each trophic group by depth. (A) abundance of all fish, (B) abundance of all fish excluding high/very high value fish and sharks. S=Shallow, M=Mid/lntermediate, D=Deep (MCE).

We tested for differences in overall abundance of each trophic group across the depth gradient, finding that abundance of herbivores, planktivores, corallivores, and piscivores and mobile invertebrate feeders declined at increased depth (Table 3). Herbivores had the greatest abundance decline—with a percentage change of-82% between shallow reefs and MCEs—followed by planktivores at −79% (Table 3). Piscivores and mobile invertebrate feeders showed the opposite pattern – with a 50% increase from shallow reefs to MCEs (Table 3). SIMPER analysis identified multiple species responsible for trophic group abundance changes across the depth gradient (Table 4). Eight out of the 56 recorded herbivore species explained over 50% of the total dissimilarity between shallow reef and MCE herbivore communities, with approximately 9% explained each by of the surgeonfishes *Acanthurus lineatus* and *Acanthurus maculiceps* – both absent from the MCEs (Table 4). Five herbivore species with significant depth effects were more abundant on shallow reefs than MCEs, while two species showed the reverse pattern (ESM Table 5). For planktivores, eight out of 82 species explained 50% of the dissimilarity between shallow and mesophotic reefs (Table 4). The triggerfish—*Odonus niger—*explained 12% of the dissimilarity, while two fusiliers *(Caesio cuning* and *Casio lunaris)* explained a further 15% – with all three species recorded at lower abundance on MCEs. Overall, depth effects were detected for six planktivores with greater abundance on shallow reefs than MCEs, and four planktivores that were more abundant on MCEs than shallow reefs (ESM Table 5). Four out of 24 corallivores explained 58% of the dissimilarity between shallow reefs and MCEs – with a single species—*Chaetodon kleinii—*explaining 24% of the dissimilarity (Table 4). Six corallivores with significant depth effects were more abundant on shallow reefs than MCEs, while two species showed the reverse (ESM Table 5). For piscivores and mobile invertebrate feeders, 13 out of 127 species explained >50% of the dissimilarity, with 34% of the contribution from four *Pentapodus* species and *Labroides dimidiatus, Carangoides ferdau,* and *Lutjanus semicinctus* (Table 4). Three of these *Pentapodus* species and *Carangoides ferdau* were at greater abundance on MCEs than shallow reefs, while the other three species were at lower abundance on MCEs. Overall, for piscivores and mobile invertebrate feeders we detected significant depth effects for 16 species that were more abundant on shallow reefs than MCEs, and two species that had the reverse pattern (ESM Table 5).

### Human impacts on reef fish abundance

Our final mixed-effects model identified significant effects of depth, benthic habitat, and distance to market for regional fisheries species abundance (Table 5). Candidate models and AICc scores are included in ESM Table 7. Regional fisheries species abundance declined as the distance to market (Sorong – the main market center) increased (Table 5; Fig. 5A) – i.e. more remote reefs had lower fish abundance. Though there was high variation, our partial effects plot indicated that increasing distance from Sorong from 80 km to 180 km, led to a 97% decline in regional fisheries species abundance from 33.3 adjusted ind./BRUV to 1.2 adjusted ind./BRUV after controlling for other factors (Fig. 5A). We found effects of depth and benthic habitat on Scarinae and Acanthuridae abundance, but no effects of the human or environmental variables so these were removed from the final model (Table 5; ESM Table 7). We also identified significant interactions between depth and benthic habitat for regional fisheries species and Scarinae and Acanthuridae. These suggested that under some benthic habitat conditions fish abundance was higher on intermediate depth reefs than shallow reefs (Table 5).

**Figure 5.**
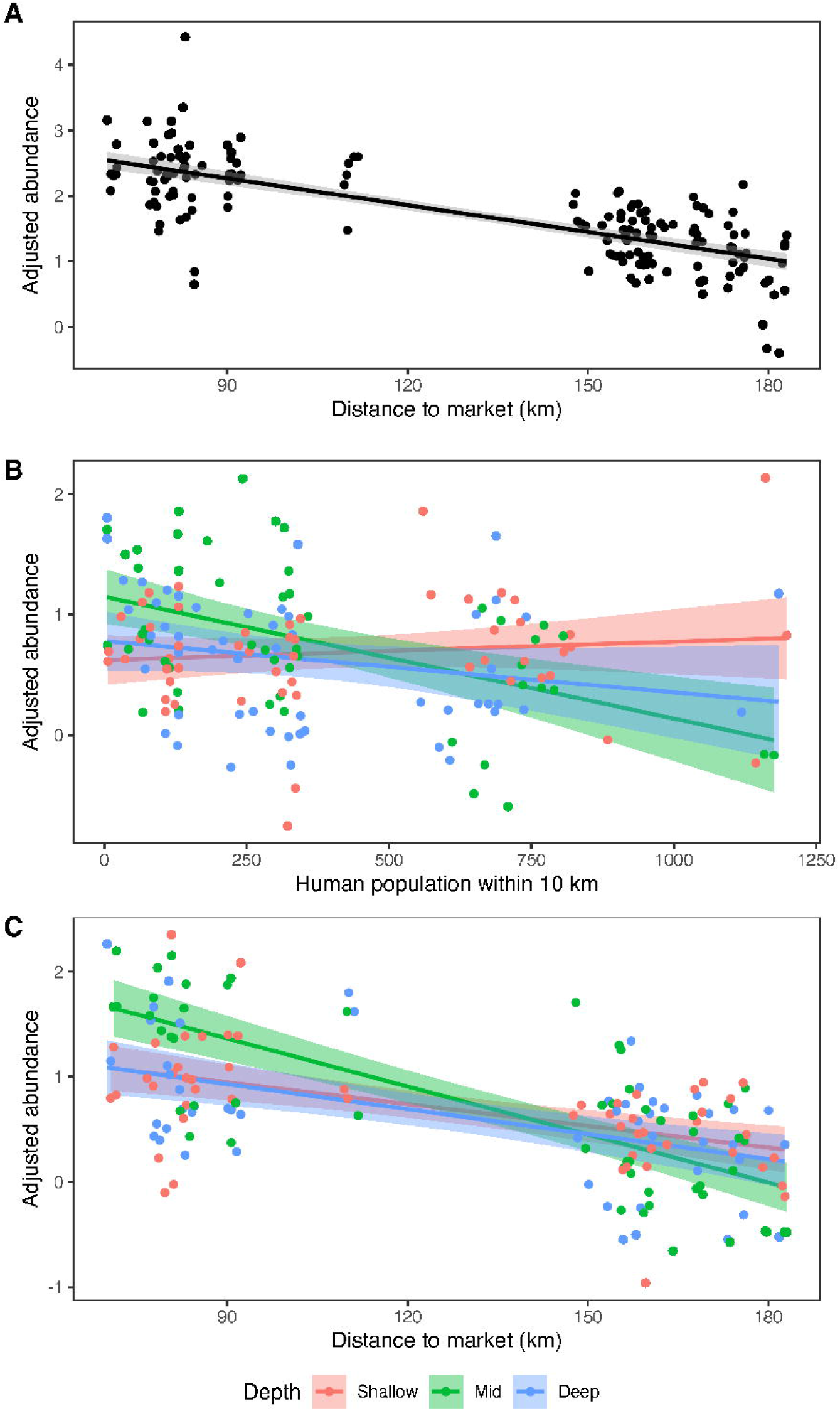
Partial effects plots for (A) distance to market based on regional fisheries abundance, (B) interaction between human population within 10 km and depth based on Epinephelidae and Lutjanidae abundance, and (C) interaction between distance to market and depth based on Epinephelidae and Lutjanidae abundance. Partial effect plots adjust the response variable (fourth root transformed fish abundance) for all model fixed (including interactions) and random effects except the plotted variables of interest. Adjusted abundance, therefore, allows the relationship between the fixed effect of interest and fish abundance to be examined after controlling for all other fixed and random parts of the model. See Table 5 for full model. Shaded areas show 95% confidence intervals around model estimates.

For Epinephelidae and Lutjanidae, in addition to benthic habitat, we identified depth:population density and depth:distance interactions. We found for intermediate depth reefs, the greater the human population within 10 km, the lower the abundance of Epinephelidae and Lutjanidae was recorded. For example, an increase in human population density from 0 to 500 people within 10 km, led to a 90% decline in Epinephelidae and Lutjanidae abundance on intermediate depth reefs, from 1.8 adjusted ind./BRUV to 0.2 adjusted ind./BRUV after controlling for other factors (Fig. 5B). This suggested intermediate depth reefs are particularly sensitive to fishing effects. We also found for intermediate depth reefs that the greater the distance to market (Sorong) the lower the Epinephelidae and Lutjanidae abundance (Table 5; Fig. 5C). Reefs at 80 km distance from Sorong were found to have a Epinephelidae and Lutjanidae adjusted abundance of 5.3 ind./BRUV after controlling for other factors, which declined to close to 0 adjusted ind./BRUV at 180 km from Sorong.

## Discussion

This study represents one of the largest *in situ* assessments of coral reef fish community structure across the shallow-mesophotic depth gradient within the Coral Triangle region, based on 160 BRUV drops conducted across the 2-85 m depth gradient. Our results show that fish community structure in Raja Ampat is strongly driven by both depth and benthic habitat complexity, and that highly diverse mesophotic fish communities exist within Raja Ampat. We identified commercially valuable and IUCN red listed fish species on MCEs and found 99 fish species at depths greater than their previously recorded fishbase depth range. We found some evidence of depleted reef fish abundance in areas with higher human population, with this effect mediated by depth. We also found that sites further from Sorong (the regional market center) have lower fish abundance. Overall, we recorded 521 fish species across all depth zones, with 152 fish species recorded on MCEs, placing Raja Ampat MCEs amongst the most fish species rich MCEs identified globally.

### Changes in benthic habitat complexity with depth

Benthic habitat structure declined with depth, in a similar pattern at both Dampier and Waigeo. Unsurprisingly, all six benthic categories recorded in HAS (rugosity, variety of growth forms, height, refuge size category, live cover, and hard substratum) were highly correlated, with many of the deepest BRUV drops associated with low relief reefs or even recording flat sandy benthos with soft corals. Our results contrast with the only previous shallow-MCE benthic habitat study within Indonesia, which was conducted within the Wakatobi Marine National Park (South East Sulawesi), where reef hard substrate cover was lowest in the shallows and increased down to 80 m depth (Bell et al. 2019). Of the previous MCE studies in the Coral Triangle, changes in benthic communities and complexity in Raja Ampat appear most similar to observations from Apo Island in the Philippines, where the proportion of sand/rubble increased exponentially with depth from 10-80 m depth (Abesamis et al. 2018). Our HAS results suggested a consistent decline in benthic habitat structure with increased depth, with intermediate depth reefs having intermediate habitat structure when compared to shallow reefs and MCEs. Studies in the Philippines have found more variable benthic community composition across the depth gradient, including that shallow benthic communities may not be a good predictor of adjacent MCE benthic communities (Dumalagan et al. 2019). Within the Coral Triangle—and for MCEs more broadly—MCEs are highly variable and influenced by local conditions, for example river outflows can increase MCE sedimentation (Cabaitan et al. 2019) or storms can damage reefs (Abesamis et al. 2018). We used BRUVs to survey benthic communities, unlike most previous MCE benthic studies from Indonesia (Bell et al. 2019) and the Philippines (Cabaitan et al. 2019; Dumalagan et al. 2019; Quimpo et al. 2019), which used diver transects or remote-operated vehicles. BRUVs have to rest on the seabed, preventing our surveys from incorporating walls or steep slopes. Therefore, the areas we surveyed are likely to have differing benthic complexity to previous studies in the region, as reef slope is an important predictor of MCE benthic communities (Lesser et al. 2018).

### Fish species richness and indicator species

The number of fish species recorded on Raja Ampat MCEs—152 species—were amongst the highest recorded to date on MCEs anywhere in the world. For example, in Hawaii 162 fish species were recorded on MCEs by diver surveys spanning 161 sites over 2010-2016 (Fukunaga et al. 2017). In the Philippines a combination of BRUV and technical diving recorded 277 fish species on MCEs (Pinheiro et al. 2019) – though only 92 MCE fish species were recorded by BRUV (Abesamis et al. 2018). These results from the Philippines, however, were based on 29 BRUV drops and so are likely under-sampling fish species richness in comparison to our 160 BRUV drops in Raja Ampat. While focused fish collection surveys detect greater fish species richness than BRUVs on MCEs (Pinheiro et al. 2019), work on Caribbean MCEs suggests that BRUVs detect greater fish species richness than standardized video transects (Andradi-Brown et al. 2016c). Given the wide variation in methods used, it is challenging to directly compare fish species richness between geographical locations. With increased survey effort—including other sampling methods such as targeted fish collection by technical diving—more fish species are likely to be recorded on Raja Ampat MCEs. Therefore, our results should be considered an underestimate of the true fish species richness on Raja Ampat MCEs. Our results, however, confirm that the exceptional fish species richness present on shallow coral reefs within Raja Ampat (Allen and Erdmann 2009) also extends into the mesophotic zone.

Our surveys indicate Raja Ampat MCEs have approximately 40% of the species richness of adjacent shallow reefs. Declines in mean fish species richness at increased depths are recognized across shallow reef and MCE depth gradients at many sites globally (Kahng et al. 2010; 2014). For example, declines have been recorded in the Caribbean (Bejarano et al. 2014; Pinheiro et al. 2016; Andradi-Brown et al. 2016b), Red Sea and Indian Ocean (Brokovich et al. 2008; Andradi-Brown et al. 2019), and Pacific (Thresher and Colin 1986; Lindfield 2015; Pyle et al. 2016; Asher et al. 2017). In the Coral Triangle, previous comparisons of whole community fish species richness have mostly been limited to smaller depth gradients. In Kimbe Bay, Papua New Guinea, fish species richness declined 15% from 5 to 30 m depth (MacDonald et al. 2016). In the Philippines, fish species richness was lower on MCEs (30-35 m) than shallow reefs (8-20 m), though declines with depth are highly variable – ranging from relatively modest declines to species richness more than halving (Quimpo et al. 2018; 2019).

We identified 12 fish indicator species of MCEs in Raja Ampat, of which ten were piscivores and/or mobile invertebrate feeding species and three were from regionally important fisheries families *(Carangoides ferdau, Lethrinus amboinensis,* and *Lethrinus lentjan).* Reef fish species composition across depth gradients are known to be driven by factors such as habitat availability (MacDonald et al. 2018), water temperature (Simon et al. 2016; Gress et al. 2017), fish morphological traits (Bridge et al. 2016), and broader trophic groups (Bejarano et al. 2014; Andradi-Brown et al. 2016b). Many of the MCE indicator species we identified have also been identified as deep species by studies in the Philippines, such as *Hoplolatilus purpureus* and *Parupeneus heptacanthus* (Abesamis et al. 2018).

Species identified as characteristic of Raja Ampat MCEs included two species of *Pentapodus – Pentapodus nagasakiensis* and *Pentapodus paradiseus. We* also found five *Pentapodus* species were identified by our piscivore and mobile invertebrate feeder SIMPER analysis, and so contributed to the trend of greater piscivore and mobile invertebrate feeder abundance on MCEs. Several *Pentapodus species* have elongate trailing caudal fin filaments (Allen and Erdmann 2012), which has been associated with reducing hydrodynamic disturbance when swimming (Bridge et al. 2016). This is believed to aid predator avoidance on deeper reefs where water turbulence is less (Bridge et al. 2016). *Pentapodus* species are also known to favor sandy areas immediately adjacent to reefs (Gomelyuk 2009), which are similar to MCE habitats we recorded in Raja Ampat. As several of our analyses have identified MCE *Pentapodus,* this genus could be used as an easy to identify indicator group for MCE fish communities in Raja Ampat.

Whilst there was broad agreement between our indicator species and other studies from the Coral Triangle, several Raja Ampat indicator species were identified at different depths to previous studies. For example, *Sufflamen fraenatum* was found to be a depth generalist between 10-80 m in the Philippines (Abesamis et al. 2018) and *Pentapodus paradiseus* was also found to be common at 30-35 m depth in the Philippines (Quimpo et al. 2018), but our indicator species analysis clustered both as MCE species in Raja Ampat. In even greater contrast, work in the Philippines identified two of our MCE indicator species, *Lethrinus lentjan* and *Pomacentrus nagasakiensis,* as shallow water species (Abesamis et al. 2018). This could reflect real differences between the Philippines survey sites and Raja Ampat (e.g. caused by habitat variation between depths at different locations) or they could be the result of methodological differences between surveys that affect detection of species. Surveys using standardized methods across the region are required to directly identify differences in the fish community, but our results imply that depth ranges should not be generalized for species across the Coral Triangle.

Our intermediate depth fish indicator species more broadly aligns with previous studies. For example, *Forcipiger flavissimus* and *Pomacentrus nigromanus,* which we identified as intermediate depth species, are both common at intermediate depths in Papua New Guinea (MacDonald et al. 2016). Studies from Papua New Guinea have highlighted the absence of Acanthuridae from MCEs (Pyle 2000; Longenecker et al. 2019), but their presence at intermediate depths and the transition zone with upper-MCEs (MacDonald et al. 2016; Longenecker et al. 2019). Comparing with our results, we find that *Acanthurus thompsoni* was identified as an intermediate depth indicator species in Raja Ampat. This species is common in coral reef communities at intermediate depths in both Papua New Guinea (MacDonald et al. 2016) and the Philippines (Quimpo et al. 2018).

### Fish abundance and community structure across the depth gradient

Fish abundance significantly declined with increased depth in Waigeo. In Dampier, however, fish abundance trended downwards at increased depth, but was not statistically significant – implying that in some rare cases MCEs may support similar fish abundances to shallow reefs. Globally, differences in fish abundance across shallow reef to MCE depth gradients are highly variable. In many locations abundance declines at increased depths, for example in Honduras (Andradi-Brown et al. 2016b), Curaçao (Pinheiro et al. 2016), Red Sea (Brokovich et al. 2008; 2010), the Chagos Archipelago (Andradi-Brown et al. 2019), and Hawaii (Asher et al. 2017). Whereas, abundance and biomass increases with depth in some locations, for example Bermuda (Pinheiro et al. 2016) and the Mariana Islands (Lindfield et al. 2016), before reducing on lower MCEs (>70 m depth). In the Coral Triangle, fish abundance reflects this variation, with declines at increased depth in Papua New Guinea and the Philippines (MacDonald et al. 2016; Quimpo et al. 2018; 2019) – but some locations where fish abundance increases from the surface to a peak at 30 m depth before declining (Abesamis et al. 2018; Quimpo et al. 2018). As we only recorded three broad depth groups, it is not possible to identify fine scale changes in fish abundance in Raja Ampat. However, our results suggest that differences in abundance with depth between Waigeo and Dampier are in part driven by changes in benthic complexity. Previous BRUV studies in Hawaii have found that benthic structure complexity alters fish patterns across the depth gradient, with unconsolidated substrates having similar overall densities of fish across the depth gradient, while hard bottom habitats show declines (Asher et al. 2017).

We found that MCE fish communities shifted towards increased mobile invertebrate feeders and piscivores at deeper depths, and away from omnivores, corallivores, and planktivores which were more common in the shallows. Increases in piscivore abundance on MCEs compared to shallow reefs is consistent with other studies, as are declines in omnivores and corallivores (Abesamis et al. 2018; Quimpo et al. 2019). However, reductions planktivores as a proportion of the fish community at increased depths is unusual. Previously, most studies—including some based on BRUV surveys—from the western Atlantic (Bejarano et al. 2014; Pinheiro et al. 2016; Andradi-Brown et al. 2016b), the Marshall Islands (Thresher and Colin 1986), the Red Sea (Brokovich et al. 2008; 2010), and the Philippines (Abesamis et al. 2018; Quimpo et al. 2018; 2019) have characterized MCEs fish communities as planktivore dominated. The decline in planktivores as a proportion of the fish community at increased depths in Raja Ampat could in part be explained by commercially valuable species. Many locations where planktivores are dominant on MCEs have experienced historically high fishing pressure – removing commercially valuable fish species. Raja Ampat has been exposed to limited fishing pressure in the past and so retains a high abundance of commercially valuable fish species. Filtering our data to remove commercially valuable fish species shifted the MCE fish community towards a greater proportion of planktivores. However, planktivores also often aggregate around areas of increased reef structure and adjacent to deep-water drop-offs (Hobson and Chess 1978; Friedlander and Parrish 1998) which may have been under sampled in our surveys. Future work should examine general trends in planktivorous MCE fish communities based on both reef structure and fisheries pressures.

### Human impacts on MCE reef fishes

Remote reefs often have greater fish abundance and biomass (Graham and McClanahan 2013; Pinheiro et al. 2016), largely attributed to increased accessibility cost and effort required for fishers (Maire et al. 2016). We found reefs further away from Sorong— the major urban center and fish market within the Raja Ampat region (Ahmadia et al. 2017)—had lower regional fisheries species abundance and lower abundance of Epinephelidae and Lutjanidae on intermediate depth reefs. At a global scale, fish biomass is often unchanged within 0-14 km of markets, but then increases exponentially away from the market (Cinner et al. 2013). Though some studies have found no effect of distance to market (e.g. Campbell et al. 2018), within Indonesia, remote shallow reefs generally have greater fish biomass (Campbell et al. 2020). While linear distance strongly correlates with market access at a global scale, it can be locally highly variable (Maire et al. 2016). For example, travel times for reefs 105-115 km from the nearest market can vary from 2-13 hours (Maire et al. 2016). Reefs in this study varied from 70-183 km from Sorong, with Dampier sites 70-112 km and Waigeo sites 148-183 km. However, while this static view of market centers may be useful at the global scale, it misses local nuance. In the period up to 2014, Waigeo reefs were visited by large freezer-equipped vessels that purchased fish from villages, effectively creating mobile market centers. These vessels were disproportionally focused on purchasing higher value fish species, such as Epinephelidae and Lutjanidae. In addition, prior to the 2012 shark fishing ban many shark fishers from Indonesian provinces such as Maluku travelled eastwards to fish within Raja Ampat regency (Jaiteh et al. 2016). Therefore, reefs further from Sorong—such as Waigeo—may actually have been more accessible to this historic fishing effort.

Tourism could also play a role in lower fish abundance closer to Sorong. Dampier MPA is subject to high and increasing reef tourism (Spalding et al. 2017; Papilaya et al. 2019). During the period 2012-2016 many residents in Dampier shifted primary occupation from wild capture fisheries to wage labor (believed to be tourism related), which was accompanied by a decline in the number of households fishing regularly (Claborn et al. 2017). Despite this decline in households fishing, Dampier residents continue to fish at the highest rate within Raja Ampat (Ahmadia et al. 2017), and demand for fish for tourist consumption is likely to be growing. We identified several important fisheries species exclusively on MCEs, which could be threatened by this fisheries demand. Tourism brings a range of sustainability and environmental management challenges for the region – including managing coastal development and pollution. Elsewhere in the world, MCEs are known to be damaged by fisheries overexploitation, sedimentation, and pollution (Andradi-Brown et al. 2016a; Rocha et al. 2018). It is therefore important for tourism development in the region to be sustainable managed to ensure that benefits are received by local communities and it does not lead to environmental degradation (Atmodjo et al. 2017; Papilaya et al. 2019).

Increased human population density in areas where local communities have high marine resource dependence is well-known to be associated with lower reef fish abundance and biomass (e.g. Bellwood et al. 2012; Williams et al. 2015). Our results show that intermediate depth reefs closer to higher population densities have lower Epinephelidae and Lutjanidae abundance. MCEs are particularly likely to act as refuges from fisheries when spear-fishing is a dominant fishing method (Lindfield et al. 2014; Andradi-Brown et al. 2017). Community surveys in Dampier region show that people typically use simple hand gears for subsistence fishing, including spear-fishing and single hand lines, but it was not reported which was the most dominant form (Claborn et al. 2017). If MCEs were acting as fisheries refuges, we would have expected to identify a significant interaction between human population density and deeper reefs, as spear-fishers would be unable to reach the full depth range of MCEs. The lack of interaction in our results suggests that hand lines are likely to be more dominant in the fishery allowing access for fishers to all depth ranges. However, the interaction between intermediate depth reefs and population density suggests that reefs in the 15-30 m depth range are particularly vulnerable to fisheries exploitation.

Caution is required, however, in interpreting fisheries impacts across the depth gradient from our fish abundance data as we were unable to record individual fish lengths and calculate fish biomass. Studies have shown fish length distributions change across shallow reef to MCE depth gradients, with larger individuals often present on deeper reefs (Andradi-Brown et al. 2016b; Gress et al. 2018). Therefore, fish biomass can show different patterns across the depth gradient to fish abundance. Further research is needed to identify whether changes in fish body size with depth occurs for species in Raja Ampat, and the effect this may have on fish biomass and fisheries impacts with depth.

Our results provide the first overview of fish communities on MCEs in Raja Ampat and represent one of the most extensive MCE fish surveys conducted in the Coral Triangle to date. These results suggest that MCEs in Raja Ampat are likely amongst the most rich in fish species globally and retain substantial abundance of higher fish trophic levels. Our findings help increase global understanding of MCE fish ecology and provide the first insight of how human interactions can shape MCEs in the center of coral reef biodiversity.

### Conflict of interest

On behalf of all authors, the corresponding author states that there is no conflict of interest.

## Supporting information

Electronic Supplementary Material

## Acknowledgments

We wish to thank: Balai Kawasan Konservasi Perairan Nasional (BKKPN) Kupang of the Ministry of Marine Affairs and Fisheries (MMAF) through the Satker KKPN Raja Ampat; the Walton Family Foundation who provide funding for the field surveys; Captain Wempy Boari and Captain Andreas of MV Imbekwan and their crew; Conservation International Indonesia staff who assisted with surveys (including: Abraham Sianipar, Ronald Mambrasar, Abdy Hasan); UPTD Raja Ampat staff who assisted with surveys (including: Aser Burdam, Risyart Mirino, Elvis Mambraku, Yance Mayor, Naftali Manggara, Imanuel Mofu); students from the International Master of Science in Marine Biodiversity and Conservation (EMBC+) and the International Master in Marine Biological Resources (IMBRSea) and the University of Oxford who assisted with BRUV video analysis. We also thank the following who assisted with this study: Edy Setyawan, Florencia Cerutti, Matthew Fox, Meity Mongdong, Timore Kristiani, Alberth Nebore, Permenas Mambrasar, Alex David Rogers, and Mark Erdmann. DAAB acknowledges a Fisheries Society of the British Isles (FSBI) PhD studentship. AJEB acknowledges a NSERC IPS grant. CEIH is supported by the Bertarelli Foundation as part of the Bertarelli Programme in Marine Science. We thank the reviewers and the editor for comments that substantially improved this manuscript.

## Notes

### Competing Interest Statement

The authors have declared no competing interest.

### Summary of Updates

This manuscript has been revised. We have removed the fish biomass analysis, and now provide an analysis of fish abundance in relation to contextual variables (e.g. distance to Sorong, human population). We also now use SIMPER analysis in place for the previous PCA axis correlations.

