## Supplementary material for "Highly diverse mesophotic reef fish communities in Raja Ampat, West Papua": Electronic Supplementary Material

ESM Table 1. Number of BRUV drops per site and depth for both regions.

A minimum of two BRUV drops were conducted per depth zone at each site with the exception of MCEs at WNTZ site 7 where recorded footage was only analyzable from one BRUV.

| Region/Site | Number of replicate BRUV drops per depth | | |
| --- | --- | --- | --- |
|  | Shallow | Intermediate | MCE |
| **Dampier (DNTZ)** |  |  |  |
| 1 | 3 | 2 | 3 |
| 2 | 3 | 2 | 2 |
| 3 | 3 | 2 | 3 |
| 4 | 2 | 2 | 2 |
| 5 | 2 | 2 | 2 |
| **Dampier (DUZ)** |  |  |  |
| 1 | 3 | 3 | 2 |
| 2 | 3 | 2 | 2 |
| 3 | 3 | 2 | 2 |
| 4 | 2 | 2 | 3 |
| 5 | 2 | 2 | 3 |
| **Waigeo (WNTZ)** |  |  |  |
| 1 | 2 | 2 | 3 |
| 2 | 2 | 3 | 2 |
| 3 | 2 | 2 | 2 |
| 4 | 2 | 2 | 2 |
| 5 | 2 | 4 | 2 |
| 6 | 2 | 2 | 2 |
| 7 | 2 | 2 | 1 |
| **Waigeo (WUZ)** |  |  |  |
| 1 | 2 | 2 | 2 |
| 2 | 2 | 2 | 2 |
| 3 | 2 | 2 | 2 |
| 4 | 2 | 2 | 2 |
| 5 | 2 | 2 | 2 |
| 6 | 2 | 4 | 2 |
| 7 | 2 | 2 | 2 |

ESM Table 2. GPS coordinates for each BRUV drop.

‘Region’ indicates both the region and fisheries type as follows: DNTZ – Dampier no-take zone, DUZ – Dampier use zone (sustainable fisheries zone within the MPA), WNTZ – Waigeo no-take zone, and WUZ – Waigeo use area (area open to fisheries outside the MPA). Depth represents depth zone as follows: S – Shallow, M – Mid (intermediate), and D – Deep (MCE). ‘Site number’ refers to the site number as used in the manuscript. The ‘Raw data site number’ and ‘Replicate numbers’ represent the original Conservation International survey site and replicate numbers, which are used in the raw data. Not all sites/replicates surveyed were analyzed in this study, hence the numbers are not continuous. All GPS points are given in WGS84 format decimal degrees.

| **Region** | **Site number** | **Raw data site number** | **Depth** | **Replicate number** | **Longitude** | **Latitude** |
| --- | --- | --- | --- | --- | --- | --- |
| DNTZ | 1 | 1 | D | 1 | 130.530697 | -0.794603 |
| DNTZ | 1 | 1 | D | 2 | 130.526488 | -0.7905658 |
| DNTZ | 1 | 1 | D | 3 | 130.523987 | -0.783556 |
| DNTZ | 1 | 1 | M | 1 | 130.524713 | -0.795062 |
| DNTZ | 1 | 1 | M | 3 | 130.52132 | -0.7972476 |
| DNTZ | 1 | 1 | S | 1 | 130.523 | -0.794309 |
| DNTZ | 1 | 1 | S | 2 | 130.520595 | -0.798318 |
| DNTZ | 1 | 1 | S | 3 | 130.532855 | -0.7987367 |
| DNTZ | 2 | 2 | D | 1 | 130.508027 | -0.7989828 |
| DNTZ | 2 | 2 | D | 2 | 130.513062 | -0.7867608 |
| DNTZ | 2 | 2 | M | 3 | 130.509315 | -0.7927859 |
| DNTZ | 2 | 2 | M | 4 | 130.515608 | -0.7952866 |
| DNTZ | 2 | 2 | S | 1 | 130.507272 | -0.793176 |
| DNTZ | 2 | 2 | S | 2 | 130.497068 | -0.794196 |
| DNTZ | 2 | 2 | S | 3 | 130.511291 | -0.8056208 |
| DNTZ | 3 | 3 | D | 1 | 130.452206 | -0.843469 |
| DNTZ | 3 | 3 | D | 2 | 130.454281 | -0.851093 |
| DNTZ | 3 | 3 | D | 3 | 130.448862 | -0.855004 |
| DNTZ | 3 | 3 | M | 1 | 130.455182 | -0.847037 |
| DNTZ | 3 | 3 | M | 2 | 130.449541 | -0.839482 |
| DNTZ | 3 | 3 | S | 1 | 130.449029 | -0.83451 |
| DNTZ | 3 | 3 | S | 2 | 130.453843 | -0.855849 |
| DNTZ | 3 | 3 | S | 3 | 130.452008 | -0.859498 |
| DNTZ | 4 | 4 | D | 2 | 130.434489 | -0.863857 |
| DNTZ | 4 | 4 | D | 3 | 130.440475 | -0.8744669 |
| DNTZ | 4 | 4 | M | 1 | 130.441935 | -0.8590817 |
| DNTZ | 4 | 4 | M | 2 | 130.449238 | -0.8638667 |
| DNTZ | 4 | 4 | S | 1 | 130.439157 | -0.857953 |
| DNTZ | 4 | 4 | S | 2 | 130.434941 | -0.860401 |
| DNTZ | 5 | 8 | D | 1 | 130.353733 | -0.49002 |
| DNTZ | 5 | 8 | D | 2 | 130.34567 | -0.488285 |
| DNTZ | 5 | 8 | M | 1 | 130.357852 | -0.487746 |
| DNTZ | 5 | 8 | M | 3 | 130.338402 | -0.4898128 |
| DNTZ | 5 | 8 | S | 1 | 130.362512 | -0.486074 |
| DNTZ | 5 | 8 | S | 3 | 130.358208 | -0.483863 |
| DUZ | 1 | 1 | D | 2 | 130.635423 | -0.788386 |
| DUZ | 1 | 1 | D | 3 | 130.641301 | -0.788727 |
| DUZ | 1 | 1 | M | 1 | 130.626402 | -0.793261 |
| DUZ | 1 | 1 | M | 2 | 130.625086 | -0.796811 |
| DUZ | 1 | 1 | M | 4 | 130.631527 | -0.791067 |
| DUZ | 1 | 1 | S | 1 | 130.626423 | -0.794241 |
| DUZ | 1 | 1 | S | 2 | 130.630811 | -0.791413 |
| DUZ | 1 | 1 | S | 3 | 130.635497 | -0.790103 |
| DUZ | 2 | 2 | D | 1 | 130.544367 | -0.82834 |
| DUZ | 2 | 2 | D | 3 | 130.527688 | -0.827546 |
| DUZ | 2 | 2 | M | 1 | 130.541011 | -0.824919 |
| DUZ | 2 | 2 | M | 2 | 130.536438 | -0.826965 |
| DUZ | 2 | 2 | S | 1 | 130.540425 | -0.824592 |
| DUZ | 2 | 2 | S | 2 | 130.536746 | -0.826036 |
| DUZ | 2 | 2 | S | 3 | 130.537214 | -0.830708 |
| DUZ | 3 | 4 | D | 1 | 130.570988 | -0.788506 |
| DUZ | 3 | 4 | D | 2 | 130.575959 | -0.787782 |
| DUZ | 3 | 4 | M | 1 | 130.575876 | -0.795152 |
| DUZ | 3 | 4 | M | 3 | 130.563995 | -0.797067 |
| DUZ | 3 | 4 | S | 1 | 130.570746 | -0.792993 |
| DUZ | 3 | 4 | S | 2 | 130.579564 | -0.797969 |
| DUZ | 3 | 4 | S | 3 | 130.561255 | -0.802375 |
| DUZ | 4 | 7 | D | 1 | 130.637406 | -0.569417 |
| DUZ | 4 | 7 | D | 2 | 130.632069 | -0.570607 |
| DUZ | 4 | 7 | D | 3 | 130.626858 | -0.571275 |
| DUZ | 4 | 7 | M | 2 | 130.623917 | -0.572667 |
| DUZ | 4 | 7 | M | 3 | 130.638624 | -0.569266 |
| DUZ | 4 | 7 | S | 2 | 130.634321 | -0.5711356 |
| DUZ | 4 | 7 | S | 3 | 130.61781 | -0.573148 |
| DUZ | 5 | 9 | D | 1 | 130.598434 | -0.598411 |
| DUZ | 5 | 9 | D | 2 | 130.603696 | -0.594229 |
| DUZ | 5 | 9 | D | 3 | 130.607279 | -0.589442 |
| DUZ | 5 | 9 | M | 1 | 130.576687 | -0.609784 |
| DUZ | 5 | 9 | M | 3 | 130.594597 | -0.599805 |
| DUZ | 5 | 9 | S | 1 | 130.566139 | -0.61247 |
| DUZ | 5 | 9 | S | 2 | 130.571207 | -0.610765 |
| WNTZ | 1 | 1 | D | 1 | 130.069786 | 0.134305 |
| WNTZ | 1 | 1 | D | 2 | 130.06853 | 0.15673002 |
| WNTZ | 1 | 1 | D | 3 | 130.070163 | 0.14955629 |
| WNTZ | 1 | 1 | M | 2 | 130.061787 | 0.15101625 |
| WNTZ | 1 | 1 | M | 3 | 130.065415 | 0.15681 |
| WNTZ | 1 | 1 | S | 2 | 130.060455 | 0.153615 |
| WNTZ | 1 | 1 | S | 3 | 130.065935 | 0.15262331 |
| WNTZ | 2 | 2 | D | 2 | 130.120479 | 0.137607 |
| WNTZ | 2 | 2 | D | 3 | 130.13899 | 0.133177 |
| WNTZ | 2 | 2 | M | 2 | 130.130658 | 0.134857 |
| WNTZ | 2 | 2 | M | 3 | 130.136195 | 0.13559832 |
| WNTZ | 2 | 2 | M | 4 | 130.147438 | 0.133255 |
| WNTZ | 2 | 2 | S | 2 | 130.128722 | 0.134322 |
| WNTZ | 2 | 2 | S | 3 | 130.142698 | 0.13507001 |
| WNTZ | 3 | 3 | D | 1 | 130.223638 | 0.09694 |
| WNTZ | 3 | 3 | D | 3 | 130.210911 | 0.105589 |
| WNTZ | 3 | 3 | M | 2 | 130.216657 | 0.1027 |
| WNTZ | 3 | 3 | M | 3 | 130.212705 | 0.10462 |
| WNTZ | 3 | 3 | S | 1 | 130.219081 | 0.099818 |
| WNTZ | 3 | 3 | S | 2 | 130.228548 | 0.095535 |
| WNTZ | 4 | 5 | D | 1 | 130.125093 | 0.12996401 |
| WNTZ | 4 | 5 | D | 3 | 130.129542 | 0.120686 |
| WNTZ | 4 | 5 | M | 2 | 130.132819 | 0.12261601 |
| WNTZ | 4 | 5 | M | 3 | 130.138764 | 0.12245701 |
| WNTZ | 4 | 5 | S | 1 | 130.123098 | 0.125012 |
| WNTZ | 4 | 5 | S | 2 | 130.133877 | 0.120297 |
| WNTZ | 5 | 6 | D | 1 | 130.076229 | 0.129769 |
| WNTZ | 5 | 6 | D | 2 | 130.067475 | 0.13330202 |
| WNTZ | 5 | 6 | M | 1 | 130.062805 | 0.13976103 |
| WNTZ | 5 | 6 | M | 2 | 130.080999 | 0.147264 |
| WNTZ | 5 | 6 | M | 3 | 130.079717 | 0.13855998 |
| WNTZ | 5 | 6 | M | 4 | 130.078602 | 0.13337703 |
| WNTZ | 5 | 6 | S | 1 | 130.072124 | 0.13152597 |
| WNTZ | 5 | 6 | S | 3 | 130.07027 | 0.13389001 |
| WNTZ | 6 | 7 | D | 1 | 129.998199 | 0.16885721 |
| WNTZ | 6 | 7 | D | 2 | 130.004007 | 0.163066 |
| WNTZ | 6 | 7 | M | 1 | 130.002036 | 0.17793842 |
| WNTZ | 6 | 7 | M | 2 | 130.002848 | 0.17372911 |
| WNTZ | 6 | 7 | S | 1 | 130.006936 | 0.17944406 |
| WNTZ | 6 | 7 | S | 2 | 130.010678 | 0.178129 |
| WNTZ | 7 | 9 | D | 3 | 130.030903 | 0.17054901 |
| WNTZ | 7 | 9 | M | 2 | 130.040895 | 0.17433504 |
| WNTZ | 7 | 9 | M | 3 | 130.030178 | 0.165611 |
| WNTZ | 7 | 9 | S | 1 | 130.02632 | 0.178 |
| WNTZ | 7 | 9 | S | 2 | 130.039838 | 0.167497 |
| WUZ | 1 | 2 | D | 1 | 130.171925 | 0.004977 |
| WUZ | 1 | 2 | D | 2 | 130.177582 | 0.006271 |
| WUZ | 1 | 2 | M | 1 | 130.168195 | -0.003426 |
| WUZ | 1 | 2 | M | 2 | 130.166462 | 0.001249 |
| WUZ | 1 | 2 | S | 2 | 130.176871 | 0.00416229 |
| WUZ | 1 | 2 | S | 3 | 130.1771 | -0.000226 |
| WUZ | 2 | 3 | D | 2 | 130.143002 | 0.004364 |
| WUZ | 2 | 3 | D | 3 | 130.118273 | 0.001968 |
| WUZ | 2 | 3 | M | 2 | 130.12406 | 0.000755 |
| WUZ | 2 | 3 | M | 3 | 130.137578 | 0.00481004 |
| WUZ | 2 | 3 | S | 1 | 130.121792 | -0.0110666 |
| WUZ | 2 | 3 | S | 2 | 130.124705 | -0.0037417 |
| WUZ | 3 | 4 | D | 2 | 130.123823 | -0.026527 |
| WUZ | 3 | 4 | D | 3 | 130.128838 | -0.025605 |
| WUZ | 3 | 4 | M | 1 | 130.146495 | -0.018221 |
| WUZ | 3 | 4 | M | 3 | 130.141314 | -0.021141 |
| WUZ | 3 | 4 | S | 1 | 130.134777 | -0.025737 |
| WUZ | 3 | 4 | S | 3 | 130.126522 | -0.036198 |
| WUZ | 4 | 5 | D | 1 | 130.159497 | -0.056352 |
| WUZ | 4 | 5 | D | 2 | 130.160028 | -0.05167 |
| WUZ | 4 | 5 | M | 1 | 130.165126 | -0.061347 |
| WUZ | 4 | 5 | M | 3 | 130.157356 | -0.045629 |
| WUZ | 4 | 5 | S | 2 | 130.154767 | -0.0556245 |
| WUZ | 4 | 5 | S | 3 | 130.15199 | -0.0405267 |
| WUZ | 5 | 6 | D | 1 | 130.174079 | -0.0831957 |
| WUZ | 5 | 6 | D | 2 | 130.188744 | -0.091517 |
| WUZ | 5 | 6 | M | 1 | 130.178132 | -0.085585 |
| WUZ | 5 | 6 | M | 3 | 130.184552 | -0.102447 |
| WUZ | 5 | 6 | S | 1 | 130.191393 | -0.098528 |
| WUZ | 5 | 6 | S | 3 | 130.179898 | -0.094061 |
| WUZ | 6 | 8 | D | 1 | 130.10519 | -0.0707138 |
| WUZ | 6 | 8 | D | 3 | 130.11609 | -0.09311 |
| WUZ | 6 | 8 | M | 1 | 130.109248 | -0.068807 |
| WUZ | 6 | 8 | M | 2 | 130.113974 | -0.089298 |
| WUZ | 6 | 8 | M | 3 | 130.114045 | -0.084448 |
| WUZ | 6 | 8 | M | 4 | 130.114045 | -0.084448 |
| WUZ | 6 | 8 | S | 1 | 130.111878 | -0.0725609 |
| WUZ | 6 | 8 | S | 2 | 130.097381 | -0.06316 |
| WUZ | 7 | 9 | D | 1 | 130.067343 | -0.067851 |
| WUZ | 7 | 9 | D | 2 | 130.043359 | -0.073457 |
| WUZ | 7 | 9 | M | 3 | 130.048681 | -0.074707 |
| WUZ | 7 | 9 | M | 4 | 130.027712 | -0.073842 |
| WUZ | 7 | 9 | S | 2 | 130.067135 | -0.072522 |
| WUZ | 7 | 9 | S | 3 | 130.038239 | -0.074227 |

ESM Table 3. Potential new species records for Raja Ampat.

Table contains species we recorded in Raja Ampat absent from Allen and Erdmann (2009) Raja Ampat records. Allen and Erdmann combined field surveys within the Bird’s Head Seascape (BHS) including Raja Ampat with literature searching to compile the most comprehensive fish species list for the region. Where Allen and Erdmann (2009) identified a species record from the wider BHS region—but not Raja Ampat—this is noted with the location. Other BHS locations include Cenderawasih Bay (CB) and Fakfak/Kaimana (FK). Species in this table represent absences from Allen and Erdmann (2009), and we believe may represent new records for Raja Ampat.

| Species | Species Authority | Notes |
| --- | --- | --- |
| *Bodianus dictynna* | Gomon, 2006 | Previous records of *Bodianus diana* (Lacepède, 1801) from the region are likely to be *Bodianus dictynna*. |
| *Carangoides oblongus* | Cuvier, 1833 |  |
| *Carcharhinus obscurus* | Lesueur, 1818 |  |
| *Cheilodipterus parazonatus* | Gon, 1993 | Previously recorded in CB |
| *Chelmon muelleri* | Klunzinger, 1879 | Previously recorded in FK |
| *Choerodon monostigma* | Ogilby, 1910 |  |
| *Chromis fumea* | Tanaka, 1917 |  |
| *Chrysiptera flavipinnis* | Allen & Robertson, 1974 |  |
| *Chrysiptera glauca* | Cuvier, 1830 | Previously recorded in CB |
| *Chrysiptera parasema* | Fowler, 1918 |  |
| *Cirrhilabrus beauperryi* | Allen, Drew & Barber, 2008 |  |
| *Cirrhilabrus cyanopleura* | Bleeker, 1851 |  |
| *Coradion melanopus* | Cuvier, 1831 | Previously recorded in CB, FK |
| *Gymnocranius elongatus* | Senta, 1973 | Previously recorded in FK |
| *Hoplolatilus chlupatyi* | Klausewitz, McCosker, Randall & Zetzsche, 1978 | Previously recorded in FK |
| *Lactoria cornuta* | Linnaeus, 1758 | Previously recorded in CB |
| *Lagocephalus sceleratus* | Gmelin, 1789 |  |
| *Lethrinus microdon* | Valenciennes, 1830 | Previously recorded in FK |
| *Lethrinus rubrioperculatus* | Sato, 1978 |  |
| *Lutjanus maxweberi* | Popta, 1921 | Previously recorded in CB |
| *Myripristis botche* | Cuvier, 1829 | Previously recorded in CB, FK |
| *Naso caesius* | Randall & Bell, 1992 |  |
| *Parupeneus chrysopleuron* | Temminck & Schlegel, 1843 | Previously recorded in FK |
| *Pentapodus caninus* | Cuvier, 1830 | Previously recorded in CB |
| *Pentapodus nagasakiensis* | Tanaka, 1915 |  |
| *Pentapodus porosus* | Valenciennes, 1830 | Previously recorded in FK |
| *Pseudojuloides splendens* | Victor, 2017 | Previously recorded in FK as *Pseudojuloides cerasinus* |
| *Rhabdamia spilota* | Allen & Kuiter, 1994 | Previously recorded in CB, FK |
| *Scomberoides tol* | Cuvier, 1832 | Previously recorded in CB, FK |
| *Siganus doliatus* | Guérin-Méneville, 1829-38 | Previously recorded in FK |
| *Valenciennea wardii* | Playfair, 1867 | Previously recorded in FK |

ESM Table 4. Depth extensions for species.

Maxiumum fishbase depth represents the maximum depth recorded for the species as stated in fishbase (Froese & Pauly, 2019). Raja Ampat depth represents the maximum, minimum, or mean BRUV drop depth that the species was recorded on. Mean depth was calculated based on weighting by fish abundance (MaxN) by depth.

| FAMILY | Species | Maximum fishbase depth | Raja Ampat depth | | |
| --- | --- | --- | --- | --- | --- |
|  |  |  | Maximum | Minimum | Mean |
| ACANTHURIDAE | *Ctenochaetus striatus* | 35 | 42 | 2 | 10 |
|  | *Naso caeruleacauda* | 45 | 50 | 2 | 16 |
|  | *Naso thynnoides* | 40 | 45 | 2 | 19 |
| APOGONIDAE | *Ostorhinchus hoevenii* | 30 | 40 | 40 | 40 |
| BALISTIDAE | *Odonus niger* | 40 | 66 | 2 | 16 |
|  | *Rhinecanthus verrucosus* | 20 | 47 | 2 | 12 |
|  | *Sufflamen chrysopterum* | 30 | 52 | 2 | 18 |
| BLENNIIDAE | *Aspidontus taeniatus* | 25 | 61 | 17 | 34 |
|  | *Meiacanthus crinitus* | 20 | 51 | 30 | 47 |
|  | *Meiacanthus grammistes* | 20 | 59 | 3 | 26 |
| CAESIONIDAE | *Pterocaesio marri* | 30 | 50 | 2 | 17 |
|  | *Pterocaesio randalli* | 30 | 40 | 40 | 40 |
| CHAETODONTIDAE | *Chaetodon oxycephalus* | 40 | 43 | 2 | 16 |
|  | *Chaetodon ulietensis* | 30 | 42 | 2 | 14 |
|  | *Chaetodon vagabundus* | 30 | 44 | 2 | 12 |
|  | *Coradion altivelis* | 30 | 60 | 2 | 27 |
| CIRRHITIDAE | *Cirrhitichthys aprinus* | 40 | 47 | 27 | 40 |
| EPHIPPIDAE | *Platax batavianus* | 40 | 64 | 27 | 46 |
|  | *Platax orbicularis* | 30 | 59 | 2 | 31 |
| GERREIDAE | *Gerres erythrourus* | 40 | 49 | 49 | 49 |
| HAEMULIDAE | *Plectorhinchus lineatus* | 35 | 59 | 4 | 32 |
|  | *Plectorhinchus polytaenia* | 40 | 41 | 8 | 28 |
| LABRIDAE | *Anampses neoguinaicus* | 30 | 52 | 3 | 29 |
|  | *Choerodon jordani* | 30 | 66 | 2 | 31 |
|  | *Choerodon zosterophorus* | 40 | 49 | 2 | 22 |
|  | *Coris dorsomacula* | 40 | 50 | 50 | 50 |
|  | *Diproctacanthus xanthurus* | 25 | 38 | 2 | 7 |
|  | *Halichoeres bicolor* | 20 | 42 | 2 | 8 |
|  | *Halichoeres chloropterus* | 10 | 27 | 2 | 12 |
|  | *Halichoeres melanochir* | 25 | 55 | 6 | 29 |
|  | *Halichoeres nigrescens* | 10 | 20 | 2 | 11 |
|  | *Halichoeres prosopeion* | 40 | 64 | 2 | 23 |
|  | *Halichoeres scapularis* | 20 | 50 | 2 | 18 |
|  | *Iniistius twistii* | 20 | 65 | 65 | 65 |
|  | *Labroides bicolor* | 40 | 60 | 3 | 21 |
|  | *Labroides dimidiatus* | 40 | 66 | 2 | 12 |
|  | *Oxycheilinus celebicus* | 40 | 47 | 2 | 9 |
|  | *Stethojulis bandanensis* | 30 | 35 | 21 | 28 |
|  | *Thalassoma amblycephalum* | 15 | 59 | 2 | 18 |
| LABRIDAE (SCARINAE) | *Bolbometopon muricatum* | 40 | 41 | 2 | 17 |
|  | *Scarus oviceps* | 20 | 40 | 2 | 15 |
|  | *Scarus tricolor* | 25 | 43 | 2 | 17 |
| LETHRINIDAE | *Gnathodentex aureolineatus* | 30 | 52 | 2 | 8 |
|  | *Lethrinus atkinsoni* | 30 | 60 | 3 | 44 |
|  | *Lethrinus amboinensis* | 30 | 61 | 2 | 36 |
|  | *Lethrinus erythropterus* | 25 | 55 | 2 | 29 |
|  | *Lethrinus harak* | 20 | 65 | 2 | 25 |
|  | *Lethrinus laticaudis* | 35 | 60 | 2 | 46 |
|  | *Lethrinus obsoletus* | 30 | 52 | 2 | 28 |
|  | *Lethrinus ornatus* | 30 | 47 | 23 | 42 |
|  | *Lethrinus semicinctus* | 35 | 66 | 2 | 27 |
| LUTJANIDAE | *Lutjanus biguttatus* | 36 | 49 | 6 | 9 |
| MICRODESMIDAE | *Gunnellichthys pleurotaenia* | 15 | 44 | 4 | 17 |
| MONACANTHIDAE | *Amanses scopas* | 18 | 55 | 4 | 25 |
|  | *Paramonacanthus japonicus* | 46 | 61 | 2 | 41 |
|  | *Pseudomonacanthus macrurus* | 10 | 18 | 18 | 18 |
| MONODACTYLIDAE | *Monodactylus argenteus* | 12 | 27 | 27 | 27 |
|  | *Upeneus tragula* | 42 | 60 | 2 | 44 |
| NEMIPTERIDAE | *Pentapodus aureofasciatus* | 35 | 55 | 2 | 21 |
|  | *Pentapodus emeryii* | 35 | 61 | 2 | 21 |
|  | *Pentapodus trivittatus* | 30 | 43 | 2 | 6 |
|  | *Scolopsis bilineata* | 25 | 47 | 2 | 15 |
|  | *Scolopsis ciliata* | 25 | 66 | 2 | 26 |
|  | *Scolopsis margaritifera* | 25 | 50 | 2 | 14 |
|  | *Scolopsis temporalis* | 35 | 51 | 2 | 13 |
|  | *Scolopsis trilineata* | 20 | 61 | 2 | 13 |
|  | *Scolopsis vosmeri* | 25 | 64 | 2 | 41 |
| PHOLIDICHTHYIDAE | *Pholidichthys leucotaenia* | 30 | 50 | 50 | 50 |
| PINGUIPEDIDAE | *Parapercis hexophtalma* | 25 | 28 | 2 | 7 |
|  | *Parapercis lineopunctata* | 35 | 51 | 3 | 26 |
|  | *Parapercis snyderi* | 40 | 51 | 41 | 42 |
|  | *Parapercis tetracantha* | 25 | 47 | 2 | 13 |
| POMACANTHIDAE | *Centropyge bicolor* | 25 | 43 | 2 | 20 |
|  | *Chaetodontoplus dimidiatus* | 35 | 60 | 2 | 31 |
|  | *Chaetodontoplus mesoleucus* | 20 | 40 | 2 | 18 |
| POMACENTRIDAE | *Amblyglyphidodon batunai* | 12 | 30 | 18 | 20 |
|  | *Amblypomacentrus clarus* | 25 | 46 | 3 | 4 |
|  | *Cheiloprion labiatus* | 3 | 35 | 35 | 35 |
|  | *Chromis margaritifer* | 20 | 52 | 2 | 11 |
|  | *Chrysiptera bleekeri* | 35 | 52 | 2 | 38 |
|  | *Chrysiptera cyanea* | 10 | 28 | 2 | 5 |
|  | *Dischistodus fasciatus* | 8 | 27 | 4 | 20 |
|  | *Dischistodus melanotus* | 12 | 35 | 3 | 5 |
|  | *Dischistodus pseudochrysopoecilus* | 5 | 27 | 2 | 7 |
|  | *Neoglyphidodon crossi* | 12 | 18 | 2 | 7 |
|  | *Neoglyphidodon oxyodon* | 4 | 21 | 5 | 16 |
|  | *Neopomacentrus filamentosus* | 12 | 27 | 2 | 23 |
|  | *Pomacentrus adelus* | 8 | 15 | 15 | 15 |
|  | *Pomacentrus amboinensis* | 40 | 59 | 2 | 25 |
|  | *Pomacentrus chrysurus* | 5 | 27 | 2 | 8 |
|  | *Pomacentrus lepidogenys* | 12 | 28 | 2 | 20 |
|  | *Pomacentrus littoralis* | 5 | 28 | 5 | 18 |
|  | *Pomacentrus nagasakiensis* | 35 | 47 | 2 | 25 |
|  | *Pomacentrus simsiang* | 10 | 35 | 3 | 10 |
|  | *Pomacentrus tripunctatus* | 3 | 35 | 35 | 35 |
|  | *Stegastes albifasciatus* | 4 | 15 | 15 | 15 |
| PSEUDOCHROMIDAE | *Pseudochromis perspicillatus* | 27 | 51 | 48 | 50 |
| TETRAODONTIDAE | *Arothron immaculatus* | 30 | 49 | 49 | 49 |
|  | *Arothron mappa* | 30 | 45 | 2 | 24 |

ESM Table 5. Change in fish species abundance with depth across all sites. Abundance represents the mean MaxN per BRUV drop within a depth zone. ‘Mid’ represents intermediate depths. Changes in abundance across the depth gradient were tested with a permutational-ANOVA of the form Abundance ~ Region + Site + Benthic PC1 + Depth. As permutational-ANOVA uses sequential sums of squares, the biomass depth effects represent the effect of depth on abundance after controlling for the effect of region, site, and benthic habitat variation. Significant p-values at the p<0.05 level are shown in bold. Abundance values represent MaxN ± Standard Error.

| Family | Species | Trophic Group | Shallow | Mid | Deep | F | P |
| --- | --- | --- | --- | --- | --- | --- | --- |
| Acanthuridae | *Acanthurus bariene* | Herbivore | 0.04 ± 0.04 | 0.02 ± 0.02 | 0.00 ± 0.00 | 0.16 | 0.913 |
| Acanthuridae | *Acanthurus blochii* | Herbivore | 1.04 ± 0.78 | 0.60 ± 0.22 | 0.00 ± 0.00 | 0.38 | 0.795 |
| Acanthuridae | *Acanthurus fowleri* | Herbivore | 0.33 ± 0.11 | 0.54 ± 0.31 | 0.02 ± 0.02 | 1.55 | 0.199 |
| Acanthuridae | *Acanthurus leucocheilus* | Herbivore | 0.15 ± 0.06 | 0.14 ± 0.06 | 0.00 ± 0.00 | 0.67 | 0.525 |
| Acanthuridae | *Acanthurus lineatus* | Herbivore | 0.87 ± 0.29 | 0.00 ± 0.00 | 0.00 ± 0.00 | 5.48 | **0.003** |
| Acanthuridae | *Acanthurus maculiceps* | Herbivore | 2.09 ± 0.99 | 0.28 ± 0.15 | 0.00 ± 0.00 | 3.22 | **0.033** |
| Acanthuridae | *Acanthurus mata* | Planktivore | 0.26 ± 0.24 | 0.42 ± 0.36 | 0.07 ± 0.07 | 0.41 | 0.724 |
| Acanthuridae | *Acanthurus nigricans* | Herbivore | 0.11 ± 0.08 | 0.04 ± 0.04 | 0.00 ± 0.00 | 2.18 | 0.104 |
| Acanthuridae | *Acanthurus nigricauda* | Omnivore | 0.67 ± 0.35 | 0.06 ± 0.04 | 0.00 ± 0.00 | 1.72 | 0.179 |
| Acanthuridae | *Acanthurus nigrofuscus* | Herbivore | 0.15 ± 0.08 | 0.16 ± 0.09 | 0.00 ± 0.00 | 0.88 | 0.430 |
| Acanthuridae | *Acanthurus nubilus* | Planktivore | 0.00 ± 0.00 | 0.02 ± 0.02 | 0.00 ± 0.00 | 1.02 | 0.461 |
| Acanthuridae | *Acanthurus olivaceus* | Planktivore | 0.24 ± 0.14 | 0.18 ± 0.07 | 0.04 ± 0.04 | 0.36 | 0.730 |
| Acanthuridae | *Acanthurus pyroferus* | Herbivore | 0.19 ± 0.10 | 0.24 ± 0.08 | 0.00 ± 0.00 | 1.50 | 0.221 |
| Acanthuridae | *Acanthurus thompsoni* | Planktivore | 0.04 ± 0.03 | 0.37 ± 0.16 | 0.00 ± 0.00 | 4.56 | **0.009** |
| Acanthuridae | *Acanthurus triostegus* | Omnivore | 0.15 ± 0.13 | 0.00 ± 0.00 | 0.00 ± 0.00 | 1.50 | 0.191 |
| Acanthuridae | *Acanthurus xanthopterus* | Planktivore | 0.13 ± 0.06 | 0.02 ± 0.02 | 0.00 ± 0.00 | 2.50 | 0.080 |
| Acanthuridae | *Ctenochaetus binotatus* | Planktivore | 0.00 ± 0.00 | 0.04 ± 0.03 | 0.00 ± 0.00 | 2.73 | **0.049** |
| Acanthuridae | *Ctenochaetus cyanocheilus* | Planktivore | 0.07 ± 0.04 | 0.12 ± 0.09 | 0.00 ± 0.00 | 0.53 | 0.623 |
| Acanthuridae | *Ctenochaetus striatus* | Planktivore | 1.17 ± 0.22 | 0.52 ± 0.17 | 0.02 ± 0.02 | 1.83 | 0.172 |
| Acanthuridae | *Ctenochaetus tominiensis* | Planktivore | 0.00 ± 0.00 | 0.08 ± 0.06 | 0.00 ± 0.00 | 3.50 | **0.017** |
| Acanthuridae | *Naso annulatus* | Herbivore | 0.28 ± 0.16 | 0.24 ± 0.16 | 0.07 ± 0.05 | 0.40 | 0.687 |
| Acanthuridae | *Naso brachycentron* | Herbivore & Planktivore | 0.31 ± 0.18 | 0.30 ± 0.26 | 0.00 ± 0.00 | 0.63 | 0.589 |
| Acanthuridae | *Naso brevirostris* | Herbivore | 0.56 ± 0.21 | 0.02 ± 0.02 | 0.00 ± 0.00 | 7.89 | **0.001** |
| Acanthuridae | *Naso caeruleacauda* | Herbivore & Planktivore | 0.31 ± 0.24 | 0.08 ± 0.08 | 0.11 ± 0.11 | 0.17 | 0.901 |
| Acanthuridae | *Naso caesius* | Herbivore & Planktivore | 0.46 ± 0.21 | 0.38 ± 0.22 | 0.02 ± 0.02 | 0.38 | 0.694 |
| Acanthuridae | *Naso hexacanthus* | Planktivore | 0.52 ± 0.26 | 0.44 ± 0.17 | 0.13 ± 0.11 | 1.09 | 0.345 |
| Acanthuridae | *Naso lituratus* | Herbivore & Planktivore | 0.43 ± 0.11 | 0.20 ± 0.07 | 0.00 ± 0.00 | 0.61 | 0.557 |
| Acanthuridae | *Naso lopezi* | Planktivore | 0.04 ± 0.03 | 0.00 ± 0.00 | 0.00 ± 0.00 | 0.28 | 0.819 |
| Acanthuridae | *Naso mcdadei* | Herbivore & Planktivore | 0.04 ± 0.04 | 0.00 ± 0.00 | 0.00 ± 0.00 | 0.46 | 0.755 |
| Acanthuridae | *Naso minor* | Herbivore & Planktivore | 0.57 ± 0.56 | 0.14 ± 0.09 | 0.20 ± 0.20 | 0.59 | 0.654 |
| Acanthuridae | *Naso thynnoides* | Omnivore | 0.06 ± 0.06 | 2.56 ± 1.13 | 0.02 ± 0.02 | 4.32 | **0.012** |
| Acanthuridae | *Naso tonganus* | Herbivore | 0.00 ± 0.00 | 0.02 ± 0.02 | 0.00 ± 0.00 | 1.47 | 0.155 |
| Acanthuridae | *Naso unicornis* | Herbivore | 0.19 ± 0.09 | 0.04 ± 0.03 | 0.00 ± 0.00 | 4.25 | **0.009** |
| Acanthuridae | *Naso vlamingii* | Herbivore | 0.24 ± 0.19 | 0.06 ± 0.03 | 0.07 ± 0.07 | 1.00 | 0.394 |
| Acanthuridae | *Paracanthurus hepatus* | Planktivore | 0.02 ± 0.02 | 0.00 ± 0.00 | 0.00 ± 0.00 | 1.11 | 0.412 |
| Acanthuridae | *Zebrasoma scopas* | Herbivore | 0.46 ± 0.10 | 0.12 ± 0.07 | 0.02 ± 0.02 | 0.96 | 0.393 |
| Acanthuridae | *Zebrasoma velifer* | Herbivore | 0.06 ± 0.04 | 0.08 ± 0.06 | 0.00 ± 0.00 | 0.92 | 0.439 |
| Antennariidae | *Antennatus rosaceus* | Piscivore | 0.00 ± 0.00 | 0.00 ± 0.00 | 0.02 ± 0.02 | 0.79 | 0.547 |
| Apogonidae | *Cheilodipterus nigrotaeniatus* | Piscivore & Mobile Invertebrate | 0.02 ± 0.02 | 0.02 ± 0.02 | 0.00 ± 0.00 | 0.16 | 0.909 |
| Apogonidae | *Cheilodipterus parazonatus* | Piscivore & Mobile Invertebrate | 0.07 ± 0.04 | 0.03 ± 0.02 | 0.09 ± 0.05 | 0.78 | 0.498 |
| Apogonidae | *Ostorhinchus cavitensis* | Mobile Invertebrate | 0.09 ± 0.09 | 0.00 ± 0.00 | 0.00 ± 0.00 | 0.32 | 0.868 |
| Apogonidae | *Ostorhinchus hoevenii* | Mobile Invertebrate | 0.00 ± 0.00 | 0.00 ± 0.00 | 0.02 ± 0.02 | 1.22 | 0.338 |
| Apogonidae | *Pristiapogon exostigma* | Mobile Invertebrate | 0.24 ± 0.24 | 0.00 ± 0.00 | 0.00 ± 0.00 | 0.32 | 0.865 |
| Apogonidae | *Rhabdamia spilota* | Planktivore | 0.00 ± 0.00 | 0.02 ± 0.02 | 0.00 ± 0.00 | 2.23 | 0.063 |
| Apogonidae | *Taeniamia fucata* | Mobile Invertebrate | 0.00 ± 0.00 | 0.08 ± 0.08 | 0.00 ± 0.00 | 1.64 | 0.137 |
| Atherinidae | *Atherinomorus lacunosus* | Planktivore | 0.17 ± 0.17 | 0.00 ± 0.00 | 0.00 ± 0.00 | 1.73 | 0.130 |
| Aulostomidae | *Aulostomus chinensis* | Piscivore & Mobile Invertebrate | 0.11 ± 0.05 | 0.04 ± 0.04 | 0.00 ± 0.00 | 0.02 | 0.982 |
| Balistidae | *Abalistes stellatus* | Piscivore & Mobile Invertebrate | 0.02 ± 0.02 | 0.05 ± 0.03 | 0.39 ± 0.10 | 7.36 | **0.001** |
| Balistidae | *Balistapus undulatus* | Mobile Invertebrate | 2.30 ± 0.32 | 0.92 ± 0.16 | 0.28 ± 0.08 | 3.76 | **0.025** |
| Balistidae | *Balistoides conspicillum* | Mobile Invertebrate | 0.04 ± 0.03 | 0.08 ± 0.05 | 0.09 ± 0.04 | 2.57 | 0.076 |
| Balistidae | *Balistoides viridescens* | Mobile Invertebrate | 0.04 ± 0.03 | 0.08 ± 0.04 | 0.04 ± 0.03 | 1.11 | 0.330 |
| Balistidae | *Canthidermis maculata* | Mobile Invertebrate & Planktivore | 0.00 ± 0.00 | 0.02 ± 0.02 | 0.00 ± 0.00 | 1.32 | 0.245 |
| Balistidae | *Melichthys niger* | Herbivore & Planktivore | 0.00 ± 0.00 | 1.46 ± 1.46 | 0.00 ± 0.00 | 0.96 | 0.479 |
| Balistidae | *Melichthys vidua* | Piscivore & Mobile Invertebrate | 0.30 ± 0.08 | 0.20 ± 0.06 | 0.04 ± 0.03 | 0.76 | 0.467 |
| Balistidae | *Odonus niger* | Planktivore | 3.39 ± 1.25 | 8.32 ± 2.12 | 1.04 ± 0.47 | 5.70 | **0.004** |
| Balistidae | *Pseudobalistes flavimarginatus* | Omnivore | 0.04 ± 0.03 | 0.06 ± 0.03 | 0.07 ± 0.05 | 0.20 | 0.842 |
| Balistidae | *Pseudobalistes fuscus* | Mobile Invertebrate | 0.19 ± 0.19 | 0.46 ± 0.44 | 0.09 ± 0.05 | 0.58 | 0.633 |
| Balistidae | *Rhinecanthus rectangulus* | Mobile Invertebrate | 0.02 ± 0.02 | 0.08 ± 0.04 | 0.00 ± 0.00 | 2.98 | **0.049** |
| Balistidae | *Rhinecanthus verrucosus* | Mobile Invertebrate | 0.07 ± 0.04 | 0.02 ± 0.02 | 0.02 ± 0.02 | 0.40 | 0.689 |
| Balistidae | *Sufflamen bursa* | Mobile Invertebrate | 0.04 ± 0.03 | 0.42 ± 0.09 | 0.28 ± 0.10 | 10.37 | **<0.001** |
| Balistidae | *Sufflamen chrysopterum* | Mobile Invertebrate & Planktivore | 0.26 ± 0.09 | 0.32 ± 0.10 | 0.13 ± 0.07 | 1.62 | 0.196 |
| Balistidae | *Sufflamen fraenatum* | Mobile Invertebrate | 0.00 ± 0.00 | 0.16 ± 0.07 | 0.24 ± 0.08 | 1.22 | 0.298 |
| Balistidae | *Xanthichthys auromarginatus* | Planktivore | 0.02 ± 0.02 | 0.00 ± 0.00 | 0.00 ± 0.00 | 0.32 | 0.858 |
| Belonidae | *Tylosurus crocodilus* | Piscivore | 0.04 ± 0.03 | 0.00 ± 0.00 | 0.00 ± 0.00 | 3.43 | **0.022** |
| Blenniidae | *Aspidontus taeniatus* | Piscivore & Mobile Invertebrate | 0.06 ± 0.04 | 0.02 ± 0.02 | 0.01 ± 0.01 | 0.24 | 0.798 |
| Blenniidae | *Ecsenius midas* | Herbivore & Planktivore | 0.00 ± 0.00 | 0.02 ± 0.02 | 0.00 ± 0.00 | 1.17 | 0.376 |
| Blenniidae | *Meiacanthus atrodorsalis* | Mobile Invertebrate & Planktivore | 0.00 ± 0.00 | 0.04 ± 0.03 | 0.00 ± 0.00 | 2.07 | 0.142 |
| Blenniidae | *Meiacanthus crinitus* | Planktivore | 0.00 ± 0.00 | 0.02 ± 0.02 | 0.09 ± 0.09 | 1.27 | 0.284 |
| Blenniidae | *Meiacanthus grammistes* | Planktivore | 0.02 ± 0.02 | 0.14 ± 0.05 | 0.04 ± 0.03 | 4.12 | **0.016** |
| Blenniidae | *Plagiotremus rhinorhynchos* | Piscivore | 0.00 ± 0.00 | 0.02 ± 0.02 | 0.00 ± 0.00 | 0.73 | 0.578 |
| Caesionidae | *Caesio caerulaurea* | Planktivore | 3.43 ± 2.80 | 0.08 ± 0.06 | 0.00 ± 0.00 | 0.41 | 0.784 |
| Caesionidae | *Caesio cuning* | Planktivore | 4.78 ± 2.15 | 5.06 ± 2.35 | 0.00 ± 0.00 | 0.86 | 0.435 |
| Caesionidae | *Caesio lunaris* | Planktivore | 4.69 ± 1.91 | 1.46 ± 0.98 | 0.24 ± 0.20 | 2.86 | 0.054 |
| Caesionidae | *Caesio teres* | Planktivore | 2.33 ± 1.38 | 1.86 ± 1.70 | 0.02 ± 0.02 | 0.19 | 0.841 |
| Caesionidae | *Gymnocaesio gymnoptera* | Planktivore | 0.13 ± 0.13 | 0.04 ± 0.04 | 0.00 ± 0.00 | 0.77 | 0.535 |
| Caesionidae | *Pterocaesio digramma* | Planktivore | 0.19 ± 0.19 | 0.84 ± 0.46 | 0.00 ± 0.00 | 1.97 | 0.133 |
| Caesionidae | *Pterocaesio marri* | Planktivore | 2.85 ± 2.15 | 3.44 ± 1.81 | 1.83 ± 1.56 | 0.50 | 0.634 |
| Caesionidae | *Pterocaesio pisang* | Planktivore | 0.02 ± 0.02 | 0.96 ± 0.67 | 0.00 ± 0.00 | 3.58 | **0.014** |
| Caesionidae | *Pterocaesio randalli* | Planktivore | 0.00 ± 0.00 | 0.00 ± 0.00 | 0.04 ± 0.04 | 0.56 | 0.668 |
| Caesionidae | *Pterocaesio tile* | Planktivore | 3.37 ± 1.53 | 0.30 ± 0.30 | 0.07 ± 0.07 | 7.08 | **0.001** |
| Carangidae | *Alectis ciliaris* | Piscivore & Mobile Invertebrate | 0.06 ± 0.04 | 0.00 ± 0.00 | 0.02 ± 0.02 | 1.66 | 0.162 |
| Carangidae | *Alepes kleinii* | Planktivore | 0.00 ± 0.00 | 0.02 ± 0.02 | 0.00 ± 0.00 | 1.08 | 0.425 |
| Carangidae | *Atule mate* | Piscivore & Mobile Invertebrate | 0.02 ± 0.02 | 0.12 ± 0.08 | 0.00 ± 0.00 | 1.12 | 0.340 |
| Carangidae | *Carangoides bajad* | Piscivore & Mobile Invertebrate | 0.39 ± 0.22 | 0.41 ± 0.28 | 0.60 ± 0.54 | 0.05 | 0.963 |
| Carangidae | *Carangoides ferdau* | Piscivore & Mobile Invertebrate | 0.17 ± 0.07 | 0.14 ± 0.05 | 0.79 ± 0.29 | 2.83 | 0.056 |
| Carangidae | *Carangoides fulvoguttatus* | Piscivore & Mobile Invertebrate | 0.09 ± 0.08 | 0.04 ± 0.04 | 0.05 ± 0.04 | 0.36 | 0.730 |
| Carangidae | *Carangoides gymnostethus* | Piscivore & Mobile Invertebrate | 0.06 ± 0.03 | 0.04 ± 0.03 | 0.00 ± 0.00 | 0.70 | 0.521 |
| Carangidae | *Carangoides oblongus* | Piscivore & Mobile Invertebrate | 0.06 ± 0.03 | 0.05 ± 0.04 | 0.17 ± 0.15 | 0.14 | 0.920 |
| Carangidae | *Carangoides plagiotaenia* | Piscivore & Mobile Invertebrate | 0.07 ± 0.06 | 0.20 ± 0.12 | 0.02 ± 0.02 | 1.41 | 0.254 |
| Carangidae | *Caranx ignobilis* | Piscivore | 0.33 ± 0.26 | 0.09 ± 0.06 | 0.07 ± 0.07 | 0.41 | 0.772 |
| Carangidae | *Caranx melampygus* | Piscivore | 1.17 ± 0.38 | 1.24 ± 0.46 | 0.32 ± 0.09 | 3.23 | **0.034** |
| Carangidae | *Caranx papuensis* | Piscivore & Mobile Invertebrate | 0.11 ± 0.08 | 0.02 ± 0.02 | 0.02 ± 0.02 | 1.88 | 0.127 |
| Carangidae | *Caranx sexfasciatus* | Mobile Invertebrate | 0.07 ± 0.06 | 0.00 ± 0.00 | 4.07 ± 4.04 | 0.99 | 0.462 |
| Carangidae | *Elagatis bipinnulata* | Mobile Invertebrate | 0.00 ± 0.00 | 0.00 ± 0.00 | 0.07 ± 0.05 | 1.06 | 0.356 |
| Carangidae | *Gnathanodon speciosus* | Piscivore & Mobile Invertebrate | 0.04 ± 0.03 | 0.24 ± 0.22 | 0.04 ± 0.04 | 1.24 | 0.295 |
| Carangidae | *Scomberoides commersonnianus* | Piscivore | 0.00 ± 0.00 | 0.12 ± 0.08 | 0.00 ± 0.00 | 2.46 | 0.076 |
| Carangidae | *Scomberoides lysan* | Piscivore & Mobile Invertebrate | 0.11 ± 0.06 | 0.00 ± 0.00 | 0.00 ± 0.00 | 3.14 | **0.037** |
| Carangidae | *Scomberoides tol* | Piscivore | 0.35 ± 0.35 | 0.00 ± 0.00 | 0.00 ± 0.00 | 0.55 | 0.687 |
| Carangidae | *Selar boops* | Mobile Invertebrate & Planktivore | 0.00 ± 0.00 | 0.00 ± 0.00 | 0.24 ± 0.24 | 0.38 | 0.832 |
| Carangidae | *Trachinotus blochii* | Mobile Invertebrate & Planktivore | 0.02 ± 0.02 | 0.00 ± 0.00 | 0.00 ± 0.00 | 1.15 | 0.409 |
| Carcharhinidae | *Carcharhinus albimarginatus* | Piscivore | 0.00 ± 0.00 | 0.00 ± 0.00 | 0.02 ± 0.02 | 0.45 | 0.758 |
| Carcharhinidae | *Carcharhinus amblyrhynchos* | Piscivore | 0.02 ± 0.02 | 0.02 ± 0.02 | 0.02 ± 0.02 | 0.00 | 0.994 |
| Carcharhinidae | *Carcharhinus melanopterus* | Piscivore & Mobile Invertebrate | 0.50 ± 0.09 | 0.29 ± 0.06 | 0.18 ± 0.06 | 4.48 | **0.013** |
| Carcharhinidae | *Carcharhinus obscurus* | Piscivore | 0.00 ± 0.00 | 0.04 ± 0.03 | 0.00 ± 0.00 | 1.97 | 0.159 |
| Carcharhinidae | *Triaenodon obesus* | Piscivore & Mobile Invertebrate | 0.00 ± 0.00 | 0.13 ± 0.05 | 0.04 ± 0.03 | 5.73 | **0.004** |
| Chaetodontidae | *Chaetodon auriga* | Mobile Invertebrate | 0.06 ± 0.04 | 0.10 ± 0.05 | 0.04 ± 0.04 | 1.30 | 0.279 |
| Chaetodontidae | *Chaetodon baronessa* | Corallivore | 0.35 ± 0.10 | 0.02 ± 0.02 | 0.00 ± 0.00 | 3.94 | **0.020** |
| Chaetodontidae | *Chaetodon bennetti* | Corallivore | 0.00 ± 0.00 | 0.02 ± 0.02 | 0.00 ± 0.00 | 2.23 | 0.070 |
| Chaetodontidae | *Chaetodon citrinellus* | Omnivore | 0.19 ± 0.08 | 0.00 ± 0.00 | 0.00 ± 0.00 | 5.52 | **0.004** |
| Chaetodontidae | *Chaetodon ephippium* | Omnivore | 0.24 ± 0.10 | 0.10 ± 0.06 | 0.00 ± 0.00 | 3.99 | **0.016** |
| Chaetodontidae | *Chaetodon kleinii* | Corallivore | 1.09 ± 0.28 | 0.84 ± 0.15 | 0.59 ± 0.18 | 0.05 | 0.958 |
| Chaetodontidae | *Chaetodon lineolatus* | Corallivore | 0.26 ± 0.08 | 0.08 ± 0.05 | 0.00 ± 0.00 | 2.30 | 0.100 |
| Chaetodontidae | *Chaetodon lunula* | Mobile Invertebrate | 0.19 ± 0.06 | 0.06 ± 0.04 | 0.00 ± 0.00 | 1.84 | 0.162 |
| Chaetodontidae | *Chaetodon lunulatus* | Corallivore | 0.48 ± 0.12 | 0.04 ± 0.04 | 0.00 ± 0.00 | 3.66 | **0.027** |
| Chaetodontidae | *Chaetodon melannotus* | Corallivore | 0.04 ± 0.04 | 0.04 ± 0.03 | 0.00 ± 0.00 | 0.29 | 0.781 |
| Chaetodontidae | *Chaetodon meyeri* | Corallivore | 0.04 ± 0.04 | 0.04 ± 0.04 | 0.00 ± 0.00 | 0.25 | 0.849 |
| Chaetodontidae | *Chaetodon ocellicaudus* | Corallivore | 0.02 ± 0.02 | 0.10 ± 0.08 | 0.00 ± 0.00 | 3.17 | **0.023** |
| Chaetodontidae | *Chaetodon octofasciatus* | Corallivore | 0.09 ± 0.05 | 0.04 ± 0.03 | 0.00 ± 0.00 | 0.21 | 0.822 |
| Chaetodontidae | *Chaetodon ornatissimus* | Corallivore | 0.04 ± 0.04 | 0.10 ± 0.05 | 0.00 ± 0.00 | 2.15 | 0.105 |
| Chaetodontidae | *Chaetodon oxycephalus* | Corallivore | 0.11 ± 0.06 | 0.04 ± 0.03 | 0.04 ± 0.04 | 0.11 | 0.906 |
| Chaetodontidae | *Chaetodon punctatofasciatus* | Corallivore | 0.04 ± 0.04 | 0.00 ± 0.00 | 0.00 ± 0.00 | 0.88 | 0.511 |
| Chaetodontidae | *Chaetodon rafflesii* | Corallivore | 0.50 ± 0.11 | 0.00 ± 0.00 | 0.00 ± 0.00 | 7.26 | **0.001** |
| Chaetodontidae | *Chaetodon selene* | Omnivore | 0.00 ± 0.00 | 0.00 ± 0.00 | 0.04 ± 0.04 | 0.67 | 0.623 |
| Chaetodontidae | *Chaetodon semeion* | Omnivore | 0.15 ± 0.06 | 0.00 ± 0.00 | 0.00 ± 0.00 | 8.24 | **0.001** |
| Chaetodontidae | *Chaetodon speculum* | Corallivore | 0.20 ± 0.08 | 0.04 ± 0.03 | 0.00 ± 0.00 | 4.00 | **0.018** |
| Chaetodontidae | *Chaetodon trifascialis* | Corallivore | 0.30 ± 0.08 | 0.00 ± 0.00 | 0.00 ± 0.00 | 5.11 | **0.006** |
| Chaetodontidae | *Chaetodon ulietensis* | Omnivore | 0.09 ± 0.07 | 0.10 ± 0.06 | 0.02 ± 0.02 | 0.53 | 0.610 |
| Chaetodontidae | *Chaetodon unimaculatus* | Mobile Invertebrate | 0.04 ± 0.04 | 0.02 ± 0.02 | 0.00 ± 0.00 | 0.51 | 0.692 |
| Chaetodontidae | *Chaetodon vagabundus* | Corallivore | 0.65 ± 0.12 | 0.22 ± 0.09 | 0.11 ± 0.08 | 3.30 | **0.039** |
| Chaetodontidae | *Chelmon muelleri* | Mobile Invertebrate | 0.00 ± 0.00 | 0.00 ± 0.00 | 0.02 ± 0.02 | 1.15 | 0.401 |
| Chaetodontidae | *Chelmon rostratus* | Mobile Invertebrate | 0.06 ± 0.06 | 0.00 ± 0.00 | 0.00 ± 0.00 | 0.65 | 0.621 |
| Chaetodontidae | *Coradion altivelis* | Omnivore | 0.02 ± 0.02 | 0.14 ± 0.07 | 0.09 ± 0.05 | 1.33 | 0.269 |
| Chaetodontidae | *Coradion chrysozonus* | Omnivore | 0.04 ± 0.03 | 0.28 ± 0.09 | 0.11 ± 0.06 | 5.80 | **0.003** |
| Chaetodontidae | *Coradion melanopus* | Omnivore | 0.02 ± 0.02 | 0.00 ± 0.00 | 0.02 ± 0.02 | 1.15 | 0.314 |
| Chaetodontidae | *Forcipiger flavissimus* | Mobile Invertebrate | 0.00 ± 0.00 | 0.08 ± 0.04 | 0.00 ± 0.00 | 6.23 | **0.002** |
| Chaetodontidae | *Forcipiger longirostris* | Mobile Invertebrate | 0.02 ± 0.02 | 0.20 ± 0.08 | 0.00 ± 0.00 | 5.40 | **0.005** |
| Chaetodontidae | *Hemitaurichthys polylepis* | Planktivore | 0.00 ± 0.00 | 0.16 ± 0.16 | 0.00 ± 0.00 | 3.22 | **0.020** |
| Chaetodontidae | *Heniochus acuminatus* | Mobile Invertebrate & Planktivore | 0.00 ± 0.00 | 0.06 ± 0.06 | 0.02 ± 0.02 | 1.28 | 0.277 |
| Chaetodontidae | *Heniochus chrysostomus* | Corallivore | 0.04 ± 0.03 | 0.04 ± 0.04 | 0.00 ± 0.00 | 0.11 | 0.921 |
| Chaetodontidae | *Heniochus monoceros* | Mobile Invertebrate | 0.00 ± 0.00 | 0.06 ± 0.04 | 0.00 ± 0.00 | 2.22 | 0.087 |
| Chaetodontidae | *Heniochus singularius* | Corallivore | 0.11 ± 0.06 | 0.04 ± 0.04 | 0.00 ± 0.00 | 0.98 | 0.400 |
| Chaetodontidae | *Heniochus varius* | Mobile Invertebrate | 0.22 ± 0.08 | 0.10 ± 0.06 | 0.00 ± 0.00 | 0.18 | 0.838 |
| Chaetodontidae | *Parachaetodon ocellatus* | Omnivore | 0.00 ± 0.00 | 0.06 ± 0.04 | 0.00 ± 0.00 | 1.98 | 0.130 |
| Chanidae | *Chanos chanos* | Planktivore | 0.09 ± 0.09 | 0.00 ± 0.00 | 0.00 ± 0.00 | 0.55 | 0.695 |
| Cirrhitidae | *Cirrhitichthys aprinus* | Mobile Invertebrate | 0.00 ± 0.00 | 0.02 ± 0.02 | 0.04 ± 0.04 | 1.06 | 0.346 |
| Cirrhitidae | *Cirrhitichthys falco* | Piscivore & Mobile Invertebrate | 0.00 ± 0.00 | 0.02 ± 0.02 | 0.00 ± 0.00 | 3.22 | **0.017** |
| Cirrhitidae | *Paracirrhites forsteri* | Piscivore | 0.09 ± 0.04 | 0.00 ± 0.00 | 0.00 ± 0.00 | 3.54 | **0.023** |
| Dasyatididae | *Pastinachus sephen* | Piscivore & Mobile Invertebrate | 0.00 ± 0.00 | 0.00 ± 0.00 | 0.02 ± 0.01 | 0.71 | 0.500 |
| Dasyatididae | *Taeniura lymma* | Mobile Invertebrate | 0.00 ± 0.00 | 0.04 ± 0.03 | 0.00 ± 0.00 | 1.90 | 0.167 |
| Diodontidae | *Diodon hystrix* | Mobile Invertebrate | 0.04 ± 0.03 | 0.02 ± 0.02 | 0.00 ± 0.00 | 0.07 | 0.889 |
| Echeneidae | *Echeneis naucrates* | Planktivore | 0.11 ± 0.06 | 0.10 ± 0.04 | 0.02 ± 0.02 | 1.48 | 0.226 |
| Ephippidae | *Platax batavianus* | Mobile Invertebrate | 0.00 ± 0.00 | 0.02 ± 0.02 | 0.02 ± 0.02 | 0.27 | 0.829 |
| Ephippidae | *Platax boersii* | Piscivore & Mobile Invertebrate | 0.04 ± 0.04 | 0.24 ± 0.09 | 0.04 ± 0.03 | 3.97 | **0.019** |
| Ephippidae | *Platax orbicularis* | Piscivore & Mobile Invertebrate | 0.15 ± 0.15 | 0.16 ± 0.14 | 0.46 ± 0.43 | 0.32 | 0.793 |
| Ephippidae | *Platax pinnatus* | Mobile Invertebrate & Planktivore | 0.02 ± 0.02 | 0.00 ± 0.00 | 0.00 ± 0.00 | 0.07 | 1.000 |
| Fistulariidae | *Fistularia commersonii* | Piscivore | 0.11 ± 0.05 | 0.08 ± 0.04 | 0.07 ± 0.04 | 0.81 | 0.447 |
| Gerreidae | *Gerres erythrourus* | Mobile Invertebrate | 0.00 ± 0.00 | 0.00 ± 0.00 | 0.02 ± 0.02 | 1.17 | 0.375 |
| Ginglymostomatidae | *Nebrius ferrugineus* | Piscivore & Mobile Invertebrate | 0.04 ± 0.03 | 0.02 ± 0.02 | 0.00 ± 0.00 | 0.03 | 0.936 |
| Gobiidae | *Amblyeleotris steinitzi* | Mobile Invertebrate | 0.00 ± 0.00 | 0.04 ± 0.04 | 0.00 ± 0.00 | 1.19 | 0.364 |
| Gobiidae | *Valenciennea strigata* | Piscivore & Mobile Invertebrate | 0.02 ± 0.02 | 0.02 ± 0.02 | 0.00 ± 0.00 | 0.06 | 0.982 |
| Gobiidae | *Valenciennea wardii* | Mobile Invertebrate | 0.00 ± 0.00 | 0.00 ± 0.00 | 0.02 ± 0.02 | 0.38 | 0.841 |
| Haemulidae | *Diagramma pictum* | Mobile Invertebrate | 0.02 ± 0.02 | 0.00 ± 0.00 | 0.00 ± 0.00 | 0.52 | 0.689 |
| Haemulidae | *Plectorhinchus albovittatus* | Piscivore & Mobile Invertebrate | 0.02 ± 0.02 | 0.00 ± 0.00 | 0.00 ± 0.00 | 0.28 | 0.906 |
| Haemulidae | *Plectorhinchus gibbosus* | Piscivore & Mobile Invertebrate | 0.00 ± 0.00 | 0.04 ± 0.03 | 0.00 ± 0.00 | 2.06 | 0.139 |
| Haemulidae | *Plectorhinchus lessonii* | Piscivore & Mobile Invertebrate | 0.04 ± 0.03 | 0.00 ± 0.00 | 0.00 ± 0.00 | 2.86 | **0.041** |
| Haemulidae | *Plectorhinchus lineatus* | Piscivore & Mobile Invertebrate | 0.04 ± 0.04 | 0.08 ± 0.08 | 0.02 ± 0.02 | 0.55 | 0.640 |
| Haemulidae | *Plectorhinchus polytaenia* | Piscivore & Mobile Invertebrate | 0.02 ± 0.02 | 0.10 ± 0.06 | 0.02 ± 0.02 | 1.74 | 0.176 |
| Haemulidae | *Plectorhinchus schotaf* | Piscivore & Mobile Invertebrate | 0.00 ± 0.00 | 0.02 ± 0.02 | 0.00 ± 0.00 | 1.03 | 0.452 |
| Hemiscylliidae | *Chiloscyllium punctatum* | Piscivore & Mobile Invertebrate | 0.00 ± 0.00 | 0.02 ± 0.02 | 0.00 ± 0.00 | 0.55 | 0.680 |
| Holocentridae | *Myripristis berndti* | Planktivore | 0.00 ± 0.00 | 0.02 ± 0.02 | 0.00 ± 0.00 | 2.23 | 0.069 |
| Holocentridae | *Myripristis botche* | Piscivore & Mobile Invertebrate | 0.07 ± 0.05 | 0.04 ± 0.04 | 0.00 ± 0.00 | 0.03 | 0.945 |
| Holocentridae | *Sargocentron caudimaculatum* | Piscivore & Mobile Invertebrate | 0.06 ± 0.04 | 0.02 ± 0.02 | 0.00 ± 0.00 | 0.42 | 0.700 |
| Holocentridae | *Sargocentron violaceum* | Piscivore & Mobile Invertebrate | 0.02 ± 0.02 | 0.00 ± 0.00 | 0.00 ± 0.00 | 1.11 | 0.407 |
| Kyphosidae | *Kyphosus bigibbus* | Omnivore | 0.41 ± 0.41 | 0.00 ± 0.00 | 0.00 ± 0.00 | 1.15 | 0.405 |
| Kyphosidae | *Kyphosus cinerascens* | Omnivore | 0.69 ± 0.65 | 0.00 ± 0.00 | 0.00 ± 0.00 | 1.26 | 0.286 |
| Kyphosidae | *Kyphosus vaigiensis* | Omnivore | 1.15 ± 1.06 | 0.00 ± 0.00 | 0.00 ± 0.00 | 1.17 | 0.351 |
| Labridae | *Anampses caeruleopunctatus* | Mobile Invertebrate | 0.02 ± 0.02 | 0.06 ± 0.03 | 0.00 ± 0.00 | 2.65 | 0.061 |
| Labridae | *Anampses geographicus* | Mobile Invertebrate | 0.02 ± 0.02 | 0.00 ± 0.00 | 0.00 ± 0.00 | 0.28 | 0.917 |
| Labridae | *Anampses meleagrides* | Mobile Invertebrate | 0.09 ± 0.07 | 0.08 ± 0.05 | 0.00 ± 0.00 | 0.64 | 0.558 |
| Labridae | *Anampses neoguinaicus* | Mobile Invertebrate | 0.06 ± 0.03 | 0.02 ± 0.02 | 0.07 ± 0.07 | 0.28 | 0.804 |
| Labridae | *Bodianus axillaris* | Mobile Invertebrate | 0.06 ± 0.03 | 0.02 ± 0.02 | 0.00 ± 0.00 | 0.07 | 0.959 |
| Labridae | *Bodianus bilunulatus* | Mobile Invertebrate | 0.02 ± 0.02 | 0.02 ± 0.02 | 0.00 ± 0.00 | 0.01 | 0.999 |
| Labridae | *Bodianus bimaculatus* | Mobile Invertebrate | 0.00 ± 0.00 | 0.02 ± 0.02 | 0.00 ± 0.00 | 0.73 | 0.573 |
| Labridae | *Bodianus diana* | Mobile Invertebrate | 0.02 ± 0.02 | 0.12 ± 0.05 | 0.00 ± 0.00 | 4.97 | **0.007** |
| Labridae | *Bodianus dictynna* | Mobile Invertebrate | 0.02 ± 0.02 | 0.18 ± 0.08 | 0.00 ± 0.00 | 3.82 | **0.017** |
| Labridae | *Bodianus mesothorax* | Mobile Invertebrate | 0.07 ± 0.04 | 0.10 ± 0.04 | 0.00 ± 0.00 | 1.33 | 0.269 |
| Labridae | *Cheilinus chlorourus* | Mobile Invertebrate | 0.02 ± 0.02 | 0.02 ± 0.02 | 0.00 ± 0.00 | 0.26 | 0.840 |
| Labridae | *Cheilinus fasciatus* | Mobile Invertebrate | 0.17 ± 0.05 | 0.24 ± 0.07 | 0.00 ± 0.00 | 3.77 | **0.024** |
| Labridae | *Cheilinus trilobatus* | Piscivore & Mobile Invertebrate | 0.06 ± 0.03 | 0.06 ± 0.03 | 0.00 ± 0.00 | 0.87 | 0.415 |
| Labridae | *Cheilinus undulatus* | Mobile Invertebrate | 0.09 ± 0.06 | 0.04 ± 0.03 | 0.07 ± 0.04 | 0.62 | 0.575 |
| Labridae | *Cheilio inermis* | Mobile Invertebrate | 0.06 ± 0.04 | 0.09 ± 0.04 | 0.02 ± 0.02 | 1.17 | 0.311 |
| Labridae | *Choerodon anchorago* | Mobile Invertebrate | 0.19 ± 0.05 | 0.02 ± 0.02 | 0.00 ± 0.00 | 3.36 | **0.038** |
| Labridae | *Choerodon jordani* | Mobile Invertebrate | 0.00 ± 0.00 | 0.00 ± 0.00 | 0.26 ± 0.11 | 7.08 | **0.001** |
| Labridae | *Choerodon monostigma* | Mobile Invertebrate | 0.00 ± 0.00 | 0.01 ± 0.01 | 0.00 ± 0.00 | 1.88 | 0.105 |
| Labridae | *Choerodon schoenleinii* | Mobile Invertebrate | 0.04 ± 0.03 | 0.00 ± 0.00 | 0.00 ± 0.00 | 0.80 | 0.438 |
| Labridae | *Choerodon zamboangae* | Mobile Invertebrate | 0.00 ± 0.00 | 0.06 ± 0.06 | 0.13 ± 0.07 | 1.47 | 0.235 |
| Labridae | *Choerodon zosterophorus* | Mobile Invertebrate | 0.00 ± 0.00 | 0.34 ± 0.12 | 0.04 ± 0.03 | 7.68 | **0.001** |
| Labridae | *Cirrhilabrus beauperryi* | Planktivore | 0.00 ± 0.00 | 0.32 ± 0.30 | 0.00 ± 0.00 | 1.36 | 0.215 |
| Labridae | *Cirrhilabrus condei* | Planktivore | 0.00 ± 0.00 | 0.00 ± 0.00 | 0.30 ± 0.30 | 1.77 | 0.116 |
| Labridae | *Cirrhilabrus cyanopleura* | Planktivore | 0.09 ± 0.09 | 0.28 ± 0.28 | 0.00 ± 0.00 | 1.04 | 0.359 |
| Labridae | *Cirrhilabrus exquisitus* | Planktivore | 0.00 ± 0.00 | 0.10 ± 0.10 | 0.00 ± 0.00 | 1.35 | 0.223 |
| Labridae | *Cirrhilabrus flavidorsalis* | Planktivore | 0.85 ± 0.75 | 0.00 ± 0.00 | 0.00 ± 0.00 | 1.99 | 0.084 |
| Labridae | *Cirrhilabrus lubbocki* | Planktivore | 0.37 ± 0.37 | 0.00 ± 0.00 | 0.00 ± 0.00 | 0.59 | 0.650 |
| Labridae | *Cirrhilabrus tonozukai* | Planktivore | 0.13 ± 0.13 | 0.00 ± 0.00 | 0.00 ± 0.00 | 1.11 | 0.409 |
| Labridae | *Coris batuensis* | Mobile Invertebrate | 0.02 ± 0.02 | 0.08 ± 0.04 | 0.00 ± 0.00 | 2.55 | 0.077 |
| Labridae | *Coris dorsomacula* | Mobile Invertebrate | 0.00 ± 0.00 | 0.00 ± 0.00 | 0.02 ± 0.02 | 2.85 | **0.024** |
| Labridae | *Coris gaimard* | Mobile Invertebrate | 0.02 ± 0.02 | 0.06 ± 0.04 | 0.02 ± 0.02 | 0.81 | 0.475 |
| Labridae | *Coris pictoides* | Mobile Invertebrate | 0.61 ± 0.33 | 0.82 ± 0.37 | 0.65 ± 0.25 | 1.35 | 0.264 |
| Labridae | *Diproctacanthus xanthurus* | Corallivore | 0.85 ± 0.30 | 0.18 ± 0.11 | 0.07 ± 0.05 | 3.62 | **0.025** |
| Labridae | *Epibulus brevis* | Piscivore & Mobile Invertebrate | 0.06 ± 0.04 | 0.02 ± 0.02 | 0.00 ± 0.00 | 0.14 | 0.896 |
| Labridae | *Epibulus insidiator* | Piscivore & Mobile Invertebrate | 0.02 ± 0.02 | 0.00 ± 0.00 | 0.00 ± 0.00 | 0.51 | 0.695 |
| Labridae | *Gomphosus varius* | Mobile Invertebrate | 0.19 ± 0.08 | 0.12 ± 0.08 | 0.00 ± 0.00 | 0.51 | 0.634 |
| Labridae | *Halichoeres bicolor* | Mobile Invertebrate | 0.46 ± 0.34 | 0.16 ± 0.12 | 0.03 ± 0.02 | 1.54 | 0.199 |
| Labridae | *Halichoeres biocellatus* | Mobile Invertebrate | 0.00 ± 0.00 | 0.02 ± 0.02 | 0.00 ± 0.00 | 1.03 | 0.465 |
| Labridae | *Halichoeres chloropterus* | Mobile Invertebrate | 0.11 ± 0.05 | 0.06 ± 0.06 | 0.02 ± 0.02 | 0.65 | 0.549 |
| Labridae | *Halichoeres chrysus* | Mobile Invertebrate | 0.02 ± 0.02 | 0.82 ± 0.26 | 0.04 ± 0.04 | 8.74 | **<0.001** |
| Labridae | *Halichoeres hortulanus* | Mobile Invertebrate | 0.24 ± 0.06 | 0.10 ± 0.05 | 0.00 ± 0.00 | 1.86 | 0.161 |
| Labridae | *Halichoeres margaritaceus* | Mobile Invertebrate | 0.06 ± 0.06 | 0.00 ± 0.00 | 0.00 ± 0.00 | 1.15 | 0.402 |
| Labridae | *Halichoeres marginatus* | Mobile Invertebrate | 0.04 ± 0.03 | 0.00 ± 0.00 | 0.00 ± 0.00 | 1.28 | 0.288 |
| Labridae | *Halichoeres melanochir* | Mobile Invertebrate | 0.02 ± 0.02 | 0.02 ± 0.02 | 0.02 ± 0.02 | 0.87 | 0.464 |
| Labridae | *Halichoeres melanurus* | Mobile Invertebrate | 0.13 ± 0.05 | 0.00 ± 0.00 | 0.00 ± 0.00 | 2.76 | 0.063 |
| Labridae | *Halichoeres melasmapomus* | Mobile Invertebrate | 0.02 ± 0.02 | 0.00 ± 0.00 | 0.00 ± 0.00 | 0.46 | 0.748 |
| Labridae | *Halichoeres miniatus* | Mobile Invertebrate | 0.04 ± 0.04 | 0.00 ± 0.00 | 0.00 ± 0.00 | 0.32 | 0.861 |
| Labridae | *Halichoeres nigrescens* | Mobile Invertebrate | 0.02 ± 0.02 | 0.02 ± 0.02 | 0.00 ± 0.00 | 0.43 | 0.704 |
| Labridae | *Halichoeres papilionaceus* | Mobile Invertebrate | 0.04 ± 0.03 | 0.00 ± 0.00 | 0.00 ± 0.00 | 1.46 | 0.243 |
| Labridae | *Halichoeres podostigma* | Mobile Invertebrate | 0.11 ± 0.06 | 0.00 ± 0.00 | 0.00 ± 0.00 | 2.61 | 0.054 |
| Labridae | *Halichoeres prosopeion* | Mobile Invertebrate | 0.07 ± 0.04 | 0.40 ± 0.11 | 0.04 ± 0.03 | 13.02 | **<0.001** |
| Labridae | *Halichoeres richmondi* | Mobile Invertebrate | 0.07 ± 0.06 | 0.00 ± 0.00 | 0.00 ± 0.00 | 1.01 | 0.373 |
| Labridae | *Halichoeres scapularis* | Mobile Invertebrate | 0.15 ± 0.10 | 0.00 ± 0.00 | 0.08 ± 0.07 | 0.77 | 0.495 |
| Labridae | *Halichoeres solorensis* | Mobile Invertebrate | 1.00 ± 0.91 | 0.04 ± 0.03 | 0.00 ± 0.00 | 0.20 | 0.937 |
| Labridae | *Halichoeres trimaculatus* | Mobile Invertebrate | 0.02 ± 0.02 | 0.00 ± 0.00 | 0.00 ± 0.00 | 2.02 | 0.088 |
| Labridae | *Hemigymnus fasciatus* | Mobile Invertebrate | 0.00 ± 0.00 | 0.02 ± 0.02 | 0.00 ± 0.00 | 1.35 | 0.228 |
| Labridae | *Hemigymnus melapterus* | Mobile Invertebrate | 0.44 ± 0.14 | 0.12 ± 0.05 | 0.00 ± 0.00 | 2.28 | 0.103 |
| Labridae | *Hologymnosus annulatus* | Piscivore | 0.31 ± 0.15 | 0.08 ± 0.05 | 0.00 ± 0.00 | 1.59 | 0.197 |
| Labridae | *Hologymnosus doliatus* | Piscivore & Mobile Invertebrate | 0.02 ± 0.02 | 0.06 ± 0.04 | 0.00 ± 0.00 | 1.66 | 0.174 |
| Labridae | *Iniistius aneitensis* | Mobile Invertebrate | 0.02 ± 0.02 | 0.00 ± 0.00 | 0.11 ± 0.05 | 1.75 | 0.174 |
| Labridae | *Iniistius twistii* | Mobile Invertebrate | 0.00 ± 0.00 | 0.00 ± 0.00 | 0.01 ± 0.01 | 0.70 | 0.592 |
| Labridae | *Labrichthys unilineatus* | Corallivore | 0.02 ± 0.02 | 0.00 ± 0.00 | 0.00 ± 0.00 | 1.32 | 0.244 |
| Labridae | *Labroides bicolor* | Piscivore & Mobile Invertebrate | 0.07 ± 0.04 | 0.04 ± 0.03 | 0.02 ± 0.02 | 0.04 | 0.968 |
| Labridae | *Labroides dimidiatus* | Piscivore & Mobile Invertebrate | 0.96 ± 0.21 | 0.74 ± 0.16 | 0.28 ± 0.11 | 1.66 | 0.188 |
| Labridae | *Labroides pectoralis* | Piscivore & Mobile Invertebrate | 0.06 ± 0.04 | 0.02 ± 0.02 | 0.00 ± 0.00 | 1.86 | 0.142 |
| Labridae | *Labropsis manabei* | Corallivore | 0.04 ± 0.04 | 0.00 ± 0.00 | 0.02 ± 0.02 | 0.54 | 0.662 |
| Labridae | *Leptojulis cyanopleura* | Planktivore | 0.17 ± 0.10 | 0.00 ± 0.00 | 0.00 ± 0.00 | 1.39 | 0.254 |
| Labridae | *Macropharyngodon negrosensis* | Mobile Invertebrate & Planktivore | 0.02 ± 0.02 | 0.14 ± 0.10 | 0.00 ± 0.00 | 1.73 | 0.144 |
| Labridae | *Novaculichthys taeniourus* | Mobile Invertebrate | 0.00 ± 0.00 | 0.06 ± 0.04 | 0.00 ± 0.00 | 1.67 | 0.196 |
| Labridae | *Oxycheilinus arenatus* | Piscivore & Mobile Invertebrate | 0.04 ± 0.03 | 0.00 ± 0.00 | 0.00 ± 0.00 | 0.91 | 0.398 |
| Labridae | *Oxycheilinus celebicus* | Piscivore & Mobile Invertebrate | 0.06 ± 0.06 | 0.14 ± 0.12 | 0.02 ± 0.02 | 1.49 | 0.222 |
| Labridae | *Oxycheilinus digramma* | Mobile Invertebrate | 0.02 ± 0.02 | 0.00 ± 0.00 | 0.00 ± 0.00 | 2.02 | 0.089 |
| Labridae | *Oxycheilinus orientalis* | Piscivore & Mobile Invertebrate | 0.00 ± 0.00 | 0.02 ± 0.02 | 0.00 ± 0.00 | 0.96 | 0.481 |
| Labridae | *Oxycheilinus unifasciatus* | Piscivore & Mobile Invertebrate | 0.06 ± 0.03 | 0.18 ± 0.09 | 0.02 ± 0.02 | 2.73 | 0.051 |
| Labridae | *Paracheilinus filamentosus* | Planktivore | 0.00 ± 0.00 | 0.02 ± 0.02 | 0.00 ± 0.00 | 1.08 | 0.437 |
| Labridae | *Paracheilinus walton* | Planktivore | 0.06 ± 0.06 | 0.00 ± 0.00 | 0.00 ± 0.00 | 0.51 | 0.693 |
| Labridae | *Pseudocoris heteroptera* | Planktivore | 0.93 ± 0.75 | 0.16 ± 0.08 | 0.07 ± 0.05 | 1.97 | 0.090 |
| Labridae | *Pseudocoris yamashiroi* | Planktivore | 0.87 ± 0.65 | 0.30 ± 0.26 | 0.00 ± 0.00 | 0.84 | 0.476 |
| Labridae | *Pseudodax moluccanus* | Omnivore | 0.02 ± 0.02 | 0.04 ± 0.03 | 0.00 ± 0.00 | 2.06 | 0.120 |
| Labridae | *Pseudojuloides splendens* | Mobile Invertebrate | 0.02 ± 0.02 | 0.00 ± 0.00 | 0.02 ± 0.02 | 0.54 | 0.601 |
| Labridae | *Stethojulis bandanensis* | Mobile Invertebrate & Planktivore | 0.00 ± 0.00 | 0.26 ± 0.18 | 0.03 ± 0.03 | 1.21 | 0.310 |
| Labridae | *Stethojulis interrupta* | Mobile Invertebrate | 0.07 ± 0.07 | 0.00 ± 0.00 | 0.00 ± 0.00 | 0.51 | 0.705 |
| Labridae | *Stethojulis strigiventer* | Mobile Invertebrate | 0.02 ± 0.02 | 0.26 ± 0.22 | 0.00 ± 0.00 | 1.62 | 0.162 |
| Labridae | *Thalassoma amblycephalum* | Planktivore | 0.76 ± 0.32 | 0.26 ± 0.19 | 0.37 ± 0.33 | 0.74 | 0.507 |
| Labridae | *Thalassoma hardwicke* | Mobile Invertebrate | 1.85 ± 0.33 | 0.02 ± 0.02 | 0.00 ± 0.00 | 12.33 | **<0.001** |
| Labridae | *Thalassoma jansenii* | Mobile Invertebrate | 0.07 ± 0.04 | 0.04 ± 0.04 | 0.00 ± 0.00 | 1.62 | 0.194 |
| Labridae | *Thalassoma lunare* | Mobile Invertebrate | 2.94 ± 0.52 | 0.38 ± 0.11 | 0.00 ± 0.00 | 12.73 | **<0.001** |
| Labridae (Scarinae) | *Bolbometopon muricatum* | Corallivore | 0.00 ± 0.00 | 0.04 ± 0.03 | 0.04 ± 0.03 | 1.24 | 0.323 |
| Labridae (Scarinae) | *Cetoscarus ocellatus* | Herbivore | 0.02 ± 0.02 | 0.02 ± 0.02 | 0.00 ± 0.00 | 0.28 | 0.817 |
| Labridae (Scarinae) | *Chlorurus bleekeri* | Herbivore | 0.07 ± 0.04 | 0.00 ± 0.00 | 0.00 ± 0.00 | 1.58 | 0.219 |
| Labridae (Scarinae) | *Chlorurus bowersi* | Omnivore | 0.02 ± 0.02 | 0.00 ± 0.00 | 0.00 ± 0.00 | 1.15 | 0.402 |
| Labridae (Scarinae) | *Chlorurus japanensis* | Herbivore | 0.02 ± 0.02 | 0.00 ± 0.00 | 0.00 ± 0.00 | 2.74 | **0.029** |
| Labridae (Scarinae) | *Chlorurus microrhinos* | Herbivore | 0.07 ± 0.04 | 0.00 ± 0.00 | 0.00 ± 0.00 | 2.51 | 0.075 |
| Labridae (Scarinae) | *Chlorurus sordidus* | Herbivore | 0.20 ± 0.07 | 0.02 ± 0.02 | 0.02 ± 0.02 | 2.82 | 0.060 |
| Labridae (Scarinae) | *Hipposcarus longiceps* | Omnivore | 0.02 ± 0.02 | 0.00 ± 0.00 | 0.00 ± 0.00 | 0.51 | 0.701 |
| Labridae (Scarinae) | *Scarus chameleon* | Herbivore | 0.07 ± 0.07 | 0.00 ± 0.00 | 0.00 ± 0.00 | 0.79 | 0.554 |
| Labridae (Scarinae) | *Scarus dimidiatus* | Herbivore | 0.11 ± 0.04 | 0.00 ± 0.00 | 0.00 ± 0.00 | 6.23 | **0.003** |
| Labridae (Scarinae) | *Scarus flavipectoralis* | Herbivore | 0.09 ± 0.04 | 0.20 ± 0.08 | 0.00 ± 0.00 | 3.23 | **0.036** |
| Labridae (Scarinae) | *Scarus forsteni* | Herbivore | 0.06 ± 0.04 | 0.00 ± 0.00 | 0.00 ± 0.00 | 1.60 | 0.216 |
| Labridae (Scarinae) | *Scarus ghobban* | Herbivore | 0.02 ± 0.02 | 0.02 ± 0.02 | 0.00 ± 0.00 | 0.19 | 0.892 |
| Labridae (Scarinae) | *Scarus niger* | Herbivore | 0.02 ± 0.02 | 0.00 ± 0.00 | 0.00 ± 0.00 | 1.73 | 0.136 |
| Labridae (Scarinae) | *Scarus oviceps* | Herbivore | 0.13 ± 0.07 | 0.10 ± 0.05 | 0.04 ± 0.04 | 0.47 | 0.639 |
| Labridae (Scarinae) | *Scarus prasiognathos* | Herbivore | 0.04 ± 0.03 | 0.02 ± 0.02 | 0.00 ± 0.00 | 0.12 | 0.841 |
| Labridae (Scarinae) | *Scarus psittacus* | Herbivore | 0.07 ± 0.04 | 0.04 ± 0.04 | 0.00 ± 0.00 | 0.97 | 0.402 |
| Labridae (Scarinae) | *Scarus quoyi* | Herbivore | 0.07 ± 0.04 | 0.00 ± 0.00 | 0.00 ± 0.00 | 1.49 | 0.205 |
| Labridae (Scarinae) | *Scarus rivulatus* | Herbivore | 0.19 ± 0.19 | 0.02 ± 0.02 | 0.00 ± 0.00 | 0.38 | 0.817 |
| Labridae (Scarinae) | *Scarus rubroviolaceus* | Herbivore | 0.00 ± 0.00 | 0.04 ± 0.03 | 0.00 ± 0.00 | 2.31 | 0.098 |
| Labridae (Scarinae) | *Scarus schlegeli* | Omnivore | 0.02 ± 0.02 | 0.00 ± 0.00 | 0.02 ± 0.02 | 0.75 | 0.473 |
| Labridae (Scarinae) | *Scarus spinus* | Herbivore | 0.04 ± 0.04 | 0.00 ± 0.00 | 0.00 ± 0.00 | 0.55 | 0.682 |
| Labridae (Scarinae) | *Scarus tricolor* | Herbivore | 0.02 ± 0.02 | 0.14 ± 0.08 | 0.02 ± 0.02 | 1.91 | 0.147 |
| Lethrinidae | *Gnathodentex aureolineatus* | Mobile Invertebrate | 0.20 ± 0.20 | 0.00 ± 0.00 | 0.09 ± 0.07 | 0.36 | 0.807 |
| Lethrinidae | *Gymnocranius elongatus* | Mobile Invertebrate | 0.00 ± 0.00 | 0.00 ± 0.00 | 0.02 ± 0.02 | 1.44 | 0.178 |
| Lethrinidae | *Gymnocranius grandoculis* | Piscivore & Mobile Invertebrate | 0.02 ± 0.02 | 0.13 ± 0.08 | 0.18 ± 0.09 | 0.75 | 0.503 |
| Lethrinidae | *Lethrinus amboinensis* | Piscivore & Mobile Invertebrate | 0.02 ± 0.02 | 0.16 ± 0.08 | 0.60 ± 0.21 | 2.40 | 0.086 |
| Lethrinidae | *Lethrinus atkinsoni* | Mobile Invertebrate | 0.02 ± 0.02 | 0.06 ± 0.04 | 0.15 ± 0.11 | 0.73 | 0.531 |
| Lethrinidae | *Lethrinus erythracanthus* | Mobile Invertebrate | 0.00 ± 0.00 | 0.02 ± 0.02 | 0.00 ± 0.00 | 2.23 | 0.069 |
| Lethrinidae | *Lethrinus erythropterus* | Piscivore & Mobile Invertebrate | 0.02 ± 0.02 | 0.00 ± 0.00 | 0.02 ± 0.02 | 1.04 | 0.351 |
| Lethrinidae | *Lethrinus harak* | Mobile Invertebrate | 0.43 ± 0.14 | 0.14 ± 0.10 | 0.65 ± 0.54 | 0.66 | 0.604 |
| Lethrinidae | *Lethrinus laticaudis* | Piscivore & Mobile Invertebrate | 0.00 ± 0.00 | 0.00 ± 0.00 | 0.09 ± 0.07 | 2.45 | 0.054 |
| Lethrinidae | *Lethrinus lentjan* | Piscivore & Mobile Invertebrate | 0.00 ± 0.00 | 0.00 ± 0.00 | 0.14 ± 0.08 | 1.69 | 0.179 |
| Lethrinidae | *Lethrinus microdon* | Piscivore & Mobile Invertebrate | 0.00 ± 0.00 | 0.17 ± 0.05 | 0.20 ± 0.09 | 1.05 | 0.348 |
| Lethrinidae | *Lethrinus nebulosus* | Piscivore & Mobile Invertebrate | 0.00 ± 0.00 | 0.16 ± 0.07 | 0.22 ± 0.14 | 0.23 | 0.844 |
| Lethrinidae | *Lethrinus obsoletus* | Mobile Invertebrate | 0.02 ± 0.02 | 0.24 ± 0.12 | 0.30 ± 0.13 | 1.41 | 0.252 |
| Lethrinidae | *Lethrinus olivaceus* | Piscivore & Mobile Invertebrate | 0.06 ± 0.03 | 0.18 ± 0.07 | 0.09 ± 0.04 | 1.61 | 0.199 |
| Lethrinidae | *Lethrinus ornatus* | Mobile Invertebrate | 0.00 ± 0.00 | 0.04 ± 0.03 | 0.17 ± 0.15 | 1.15 | 0.337 |
| Lethrinidae | *Lethrinus ravus* | Mobile Invertebrate | 0.00 ± 0.00 | 0.01 ± 0.01 | 0.00 ± 0.00 | 1.88 | 0.109 |
| Lethrinidae | *Lethrinus rubrioperculatus* | Piscivore & Mobile Invertebrate | 0.00 ± 0.00 | 0.00 ± 0.00 | 0.17 ± 0.17 | 0.99 | 0.467 |
| Lethrinidae | *Lethrinus semicinctus* | Piscivore & Mobile Invertebrate | 0.11 ± 0.05 | 0.78 ± 0.45 | 0.92 ± 0.32 | 1.00 | 0.392 |
| Lethrinidae | *Lethrinus variegatus* | Mobile Invertebrate | 0.00 ± 0.00 | 0.01 ± 0.01 | 0.00 ± 0.00 | 1.88 | 0.104 |
| Lethrinidae | *Monotaxis grandoculis* | Mobile Invertebrate | 0.04 ± 0.03 | 0.24 ± 0.09 | 0.05 ± 0.03 | 3.57 | **0.028** |
| Lethrinidae | *Monotaxis heterodon* | Mobile Invertebrate | 0.06 ± 0.04 | 0.10 ± 0.08 | 0.00 ± 0.00 | 1.20 | 0.311 |
| Lutjanidae | *Aphareus furca* | Piscivore & Mobile Invertebrate | 0.00 ± 0.00 | 0.02 ± 0.02 | 0.11 ± 0.11 | 2.01 | 0.087 |
| Lutjanidae | *Aprion virescens* | Piscivore & Mobile Invertebrate | 0.04 ± 0.03 | 0.16 ± 0.05 | 0.14 ± 0.05 | 1.31 | 0.278 |
| Lutjanidae | *Lutjanus argentimaculatus* | Piscivore & Mobile Invertebrate | 0.00 ± 0.00 | 0.00 ± 0.00 | 0.02 ± 0.02 | 0.48 | 0.742 |
| Lutjanidae | *Lutjanus biguttatus* | Piscivore & Mobile Invertebrate | 0.57 ± 0.57 | 0.06 ± 0.04 | 0.04 ± 0.03 | 0.24 | 0.912 |
| Lutjanidae | *Lutjanus bohar* | Mobile Invertebrate | 0.65 ± 0.15 | 0.32 ± 0.10 | 0.07 ± 0.04 | 3.02 | **0.046** |
| Lutjanidae | *Lutjanus carponotatus* | Piscivore & Mobile Invertebrate | 0.22 ± 0.10 | 0.02 ± 0.02 | 0.00 ± 0.00 | 0.40 | 0.726 |
| Lutjanidae | *Lutjanus decussatus* | Piscivore & Mobile Invertebrate | 0.39 ± 0.09 | 0.08 ± 0.04 | 0.00 ± 0.00 | 4.54 | **0.012** |
| Lutjanidae | *Lutjanus ehrenbergii* | Piscivore & Mobile Invertebrate | 0.09 ± 0.06 | 0.00 ± 0.00 | 0.00 ± 0.00 | 0.37 | 0.782 |
| Lutjanidae | *Lutjanus fulviflamma* | Piscivore & Mobile Invertebrate | 0.04 ± 0.03 | 0.00 ± 0.00 | 0.00 ± 0.00 | 0.88 | 0.407 |
| Lutjanidae | *Lutjanus fulvus* | Piscivore & Mobile Invertebrate | 0.22 ± 0.09 | 0.00 ± 0.00 | 0.00 ± 0.00 | 1.76 | 0.162 |
| Lutjanidae | *Lutjanus gibbus* | Mobile Invertebrate | 0.24 ± 0.24 | 0.42 ± 0.24 | 0.00 ± 0.00 | 1.31 | 0.272 |
| Lutjanidae | *Lutjanus lemniscatus* | Piscivore & Mobile Invertebrate | 0.04 ± 0.03 | 0.02 ± 0.02 | 0.02 ± 0.02 | 0.49 | 0.609 |
| Lutjanidae | *Lutjanus lunulatus* | Piscivore & Mobile Invertebrate | 0.06 ± 0.06 | 0.00 ± 0.00 | 0.00 ± 0.00 | 0.52 | 0.691 |
| Lutjanidae | *Lutjanus lutjanus* | Piscivore & Mobile Invertebrate | 0.04 ± 0.04 | 0.02 ± 0.02 | 0.00 ± 0.00 | 0.24 | 0.888 |
| Lutjanidae | *Lutjanus maxweberi* | Piscivore & Mobile Invertebrate | 0.02 ± 0.02 | 0.00 ± 0.00 | 0.00 ± 0.00 | 0.89 | 0.495 |
| Lutjanidae | *Lutjanus monostigma* | Piscivore | 0.07 ± 0.04 | 0.12 ± 0.05 | 0.00 ± 0.00 | 2.17 | 0.116 |
| Lutjanidae | *Lutjanus rivulatus* | Piscivore & Mobile Invertebrate | 0.15 ± 0.07 | 0.04 ± 0.03 | 0.00 ± 0.00 | 2.65 | 0.059 |
| Lutjanidae | *Lutjanus russellii* | Piscivore & Mobile Invertebrate | 0.15 ± 0.06 | 0.00 ± 0.00 | 0.04 ± 0.04 | 3.48 | **0.028** |
| Lutjanidae | *Lutjanus semicinctus* | Piscivore & Mobile Invertebrate | 0.81 ± 0.12 | 0.22 ± 0.06 | 0.00 ± 0.00 | 9.17 | **<0.001** |
| Lutjanidae | *Lutjanus vitta* | Piscivore & Mobile Invertebrate | 0.04 ± 0.03 | 0.42 ± 0.33 | 0.20 ± 0.08 | 0.60 | 0.643 |
| Lutjanidae | *Macolor macularis* | Piscivore & Mobile Invertebrate | 0.20 ± 0.11 | 0.54 ± 0.19 | 0.00 ± 0.00 | 4.42 | **0.009** |
| Lutjanidae | *Macolor niger* | Piscivore & Mobile Invertebrate | 0.11 ± 0.06 | 0.06 ± 0.03 | 0.00 ± 0.00 | 0.37 | 0.698 |
| Lutjanidae | *Pinjalo lewisi* | Mobile Invertebrate & Planktivore | 0.00 ± 0.00 | 0.96 ± 0.96 | 0.00 ± 0.00 | 1.02 | 0.469 |
| Malacanthidae | *Hoplolatilus chlupatyi* | Planktivore | 0.00 ± 0.00 | 0.02 ± 0.02 | 0.02 ± 0.02 | 0.32 | 0.794 |
| Malacanthidae | *Hoplolatilus cuniculus* | Planktivore | 0.00 ± 0.00 | 0.00 ± 0.00 | 0.07 ± 0.05 | 0.97 | 0.376 |
| Malacanthidae | *Hoplolatilus purpureus* | Planktivore | 0.00 ± 0.00 | 0.00 ± 0.00 | 0.39 ± 0.17 | 2.40 | **0.092** |
| Malacanthidae | *Malacanthus brevirostris* | Mobile Invertebrate & Planktivore | 0.00 ± 0.00 | 0.02 ± 0.02 | 0.07 ± 0.05 | 0.32 | 0.748 |
| Malacanthidae | *Malacanthus latovittatus* | Mobile Invertebrate | 0.04 ± 0.04 | 0.02 ± 0.02 | 0.04 ± 0.03 | 0.25 | 0.795 |
| Microdesmidae | *Gunnellichthys monostigma* | Mobile Invertebrate & Planktivore | 0.00 ± 0.00 | 0.02 ± 0.02 | 0.00 ± 0.00 | 0.68 | 0.602 |
| Microdesmidae | *Gunnellichthys pleurotaenia* | Mobile Invertebrate & Planktivore | 0.06 ± 0.03 | 0.00 ± 0.00 | 0.02 ± 0.02 | 1.30 | 0.302 |
| Monacanthidae | *Aluterus scriptus* | Omnivore | 0.04 ± 0.03 | 0.00 ± 0.00 | 0.09 ± 0.05 | 3.73 | **0.018** |
| Monacanthidae | *Amanses scopas* | Piscivore & Mobile Invertebrate | 0.06 ± 0.03 | 0.06 ± 0.03 | 0.02 ± 0.02 | 0.08 | 0.938 |
| Monacanthidae | *Cantherhines dumerilii* | Mobile Invertebrate | 0.15 ± 0.09 | 0.04 ± 0.03 | 0.05 ± 0.03 | 0.30 | 0.783 |
| Monacanthidae | *Cantherhines fronticinctus* | Mobile Invertebrate | 0.04 ± 0.03 | 0.00 ± 0.00 | 0.00 ± 0.00 | 2.69 | 0.051 |
| Monacanthidae | *Cantherhines pardalis* | Mobile Invertebrate | 0.00 ± 0.00 | 0.02 ± 0.02 | 0.00 ± 0.00 | 1.23 | 0.293 |
| Monacanthidae | *Paraluteres prionurus* | Omnivore | 0.00 ± 0.00 | 0.14 ± 0.09 | 0.00 ± 0.00 | 2.86 | **0.035** |
| Monacanthidae | *Paramonacanthus japonicus* | Omnivore | 0.00 ± 0.00 | 0.00 ± 0.00 | 0.03 ± 0.02 | 0.67 | 0.558 |
| Monacanthidae | *Pervagor janthinosoma* | Omnivore | 0.04 ± 0.04 | 0.00 ± 0.00 | 0.00 ± 0.00 | 0.19 | 0.940 |
| Monacanthidae | *Pseudomonacanthus macrurus* | Piscivore & Mobile Invertebrate | 0.00 ± 0.00 | 0.00 ± 0.00 | 0.02 ± 0.02 | 1.38 | 0.224 |
| Monodactylidae | *Monodactylus argenteus* | Planktivore | 0.00 ± 0.00 | 0.02 ± 0.02 | 0.00 ± 0.00 | 1.15 | 0.389 |
| Mugilidae | *Crenimugil crenilabis* | Planktivore | 0.81 ± 0.81 | 0.00 ± 0.00 | 0.00 ± 0.00 | 0.79 | 0.555 |
| Mullidae | *Mulloidichthys flavolineatus* | Mobile Invertebrate | 0.06 ± 0.06 | 0.02 ± 0.02 | 0.11 ± 0.08 | 0.48 | 0.655 |
| Mullidae | *Mulloidichthys vanicolensis* | Piscivore & Mobile Invertebrate | 0.00 ± 0.00 | 0.00 ± 0.00 | 0.02 ± 0.02 | 1.38 | 0.224 |
| Mullidae | *Parupeneus barberinoides* | Mobile Invertebrate | 0.00 ± 0.00 | 0.02 ± 0.02 | 0.00 ± 0.00 | 1.81 | 0.108 |
| Mullidae | *Parupeneus barberinus* | Mobile Invertebrate | 0.56 ± 0.09 | 0.76 ± 0.16 | 0.30 ± 0.09 | 3.34 | **0.035** |
| Mullidae | *Parupeneus chrysopleuron* | Mobile Invertebrate | 0.00 ± 0.00 | 0.02 ± 0.02 | 0.00 ± 0.00 | 1.43 | 0.177 |
| Mullidae | *Parupeneus crassilabris* | Mobile Invertebrate | 0.35 ± 0.09 | 0.06 ± 0.04 | 0.00 ± 0.00 | 3.47 | **0.030** |
| Mullidae | *Parupeneus cyclostomus* | Piscivore & Mobile Invertebrate | 0.13 ± 0.05 | 0.52 ± 0.16 | 0.35 ± 0.12 | 2.19 | 0.114 |
| Mullidae | *Parupeneus heptacanthus* | Piscivore & Mobile Invertebrate | 0.00 ± 0.00 | 0.10 ± 0.07 | 0.30 ± 0.13 | 2.54 | 0.073 |
| Mullidae | *Parupeneus indicus* | Mobile Invertebrate | 0.09 ± 0.05 | 0.08 ± 0.06 | 0.17 ± 0.11 | 0.21 | 0.821 |
| Mullidae | *Parupeneus multifasciatus* | Mobile Invertebrate | 0.94 ± 0.14 | 1.28 ± 0.19 | 0.65 ± 0.25 | 2.34 | 0.100 |
| Mullidae | *Parupeneus pleurostigma* | Mobile Invertebrate | 0.00 ± 0.00 | 0.08 ± 0.06 | 0.00 ± 0.00 | 1.49 | 0.220 |
| Mullidae | *Upeneus tragula* | Mobile Invertebrate | 0.04 ± 0.03 | 0.04 ± 0.03 | 0.20 ± 0.16 | 0.85 | 0.472 |
| Muraenidae | *Echidna nebulosa* | Piscivore & Mobile Invertebrate | 0.02 ± 0.02 | 0.00 ± 0.00 | 0.00 ± 0.00 | 2.74 | **0.030** |
| Muraenidae | *Gymnothorax chilospilus* | Piscivore & Mobile Invertebrate | 0.02 ± 0.02 | 0.00 ± 0.00 | 0.00 ± 0.00 | 2.07 | **0.080** |
| Muraenidae | *Gymnothorax enigmaticus* | Piscivore & Mobile Invertebrate | 0.02 ± 0.02 | 0.00 ± 0.00 | 0.00 ± 0.00 | 1.18 | 0.369 |
| Muraenidae | *Gymnothorax flavimarginatus* | Piscivore & Mobile Invertebrate | 0.02 ± 0.02 | 0.00 ± 0.00 | 0.00 ± 0.00 | 0.31 | 0.879 |
| Muraenidae | *Gymnothorax javanicus* | Piscivore & Mobile Invertebrate | 0.13 ± 0.05 | 0.06 ± 0.03 | 0.00 ± 0.00 | 0.04 | 0.971 |
| Muraenidae | *Gymnothorax undulatus* | Piscivore | 0.00 ± 0.00 | 0.02 ± 0.02 | 0.00 ± 0.00 | 2.27 | 0.055 |
| Muraenidae | *Gymnothorax zonipectis* | Piscivore & Mobile Invertebrate | 0.00 ± 0.00 | 0.02 ± 0.02 | 0.00 ± 0.00 | 1.39 | 0.200 |
| Nemipteridae | *Pentapodus aureofasciatus* | Piscivore & Mobile Invertebrate | 0.06 ± 0.03 | 1.72 ± 0.44 | 1.46 ± 0.71 | 3.22 | **0.040** |
| Nemipteridae | *Pentapodus caninus* | Piscivore & Mobile Invertebrate | 0.02 ± 0.02 | 1.48 ± 0.78 | 3.39 ± 1.66 | 2.36 | 0.095 |
| Nemipteridae | *Pentapodus emeryii* | Piscivore & Mobile Invertebrate | 0.17 ± 0.10 | 0.64 ± 0.13 | 0.82 ± 0.26 | 1.90 | 0.150 |
| Nemipteridae | *Pentapodus nagasakiensis* | Mobile Invertebrate | 0.00 ± 0.00 | 0.38 ± 0.20 | 2.65 ± 1.35 | 1.04 | 0.376 |
| Nemipteridae | *Pentapodus numberii* | Piscivore & Mobile Invertebrate | 0.00 ± 0.00 | 0.10 ± 0.05 | 0.70 ± 0.61 | 0.79 | 0.550 |
| Nemipteridae | *Pentapodus paradiseus* | Piscivore & Mobile Invertebrate | 0.04 ± 0.04 | 0.04 ± 0.04 | 1.66 ± 0.79 | 4.17 | **0.009** |
| Nemipteridae | *Pentapodus porosus* | Piscivore & Mobile Invertebrate | 0.04 ± 0.03 | 0.62 ± 0.27 | 0.48 ± 0.46 | 0.51 | 0.667 |
| Nemipteridae | *Pentapodus trivittatus* | Piscivore & Mobile Invertebrate | 0.85 ± 0.19 | 0.18 ± 0.08 | 0.07 ± 0.05 | 3.09 | **0.046** |
| Nemipteridae | *Scolopsis affinis* | Mobile Invertebrate | 0.52 ± 0.38 | 0.65 ± 0.29 | 1.05 ± 0.24 | 0.20 | 0.831 |
| Nemipteridae | *Scolopsis bilineata* | Mobile Invertebrate | 0.52 ± 0.10 | 0.34 ± 0.09 | 0.04 ± 0.04 | 0.84 | 0.447 |
| Nemipteridae | *Scolopsis ciliata* | Piscivore & Mobile Invertebrate | 0.19 ± 0.08 | 0.12 ± 0.07 | 0.14 ± 0.10 | 0.08 | 0.930 |
| Nemipteridae | *Scolopsis lineata* | Piscivore & Mobile Invertebrate | 0.30 ± 0.24 | 0.00 ± 0.00 | 0.00 ± 0.00 | 0.86 | 0.508 |
| Nemipteridae | *Scolopsis margaritifera* | Mobile Invertebrate | 0.20 ± 0.08 | 0.56 ± 0.35 | 0.15 ± 0.10 | 1.50 | 0.220 |
| Nemipteridae | *Scolopsis monogramma* | Piscivore & Mobile Invertebrate | 0.06 ± 0.06 | 0.37 ± 0.32 | 0.04 ± 0.03 | 1.36 | 0.254 |
| Nemipteridae | *Scolopsis temporalis* | Piscivore & Mobile Invertebrate | 0.20 ± 0.09 | 0.70 ± 0.40 | 0.11 ± 0.05 | 1.90 | 0.116 |
| Nemipteridae | *Scolopsis trilineata* | Piscivore & Mobile Invertebrate | 0.19 ± 0.10 | 0.02 ± 0.02 | 0.02 ± 0.02 | 0.85 | 0.452 |
| Nemipteridae | *Scolopsis vosmeri* | Mobile Invertebrate | 0.00 ± 0.00 | 0.00 ± 0.00 | 0.43 ± 0.17 | 8.12 | **<0.001** |
| Nemipteridae | *Scolopsis xenochroa* | Mobile Invertebrate | 0.20 ± 0.08 | 0.10 ± 0.07 | 0.25 ± 0.14 | 0.39 | 0.702 |
| Ophichthidae | *Myrichthys colubrinus* | Mobile Invertebrate | 0.02 ± 0.02 | 0.00 ± 0.00 | 0.00 ± 0.00 | 0.27 | 0.914 |
| Ostraciidae | *Lactoria cornuta* | Mobile Invertebrate | 0.00 ± 0.00 | 0.02 ± 0.02 | 0.00 ± 0.00 | 0.83 | 0.533 |
| Ostraciidae | *Ostracion cubicus* | Mobile Invertebrate | 0.00 ± 0.00 | 0.02 ± 0.02 | 0.00 ± 0.00 | 1.47 | 0.159 |
| Ostraciidae | *Ostracion meleagris* | Omnivore | 0.02 ± 0.02 | 0.00 ± 0.00 | 0.00 ± 0.00 | 1.11 | 0.411 |
| Pempheridae | *Parapriacanthus ransonneti* | Planktivore | 0.00 ± 0.00 | 1.32 ± 1.32 | 0.00 ± 0.00 | 1.88 | 0.108 |
| Pholidichthyidae | *Pholidichthys leucotaenia* | Mobile Invertebrate | 0.00 ± 0.00 | 0.00 ± 0.00 | 0.48 ± 0.48 | 2.85 | **0.026** |
| Pinguipedidae | *Parapercis clathrata* | Mobile Invertebrate | 0.04 ± 0.03 | 0.00 ± 0.00 | 0.04 ± 0.04 | 0.66 | 0.570 |
| Pinguipedidae | *Parapercis cylindrica* | Mobile Invertebrate | 0.00 ± 0.00 | 0.04 ± 0.03 | 0.00 ± 0.00 | 1.39 | 0.257 |
| Pinguipedidae | *Parapercis hexophtalma* | Mobile Invertebrate | 0.07 ± 0.04 | 0.02 ± 0.02 | 0.01 ± 0.01 | 0.11 | 0.887 |
| Pinguipedidae | *Parapercis lineopunctata* | Mobile Invertebrate | 0.02 ± 0.02 | 0.06 ± 0.04 | 0.02 ± 0.02 | 0.50 | 0.641 |
| Pinguipedidae | *Parapercis snyderi* | Mobile Invertebrate | 0.00 ± 0.00 | 0.00 ± 0.00 | 0.13 ± 0.11 | 1.30 | 0.273 |
| Pinguipedidae | *Parapercis tetracantha* | Mobile Invertebrate | 0.04 ± 0.04 | 0.10 ± 0.06 | 0.11 ± 0.06 | 0.35 | 0.712 |
| Pinguipedidae | *Parapercis xanthozona* | Mobile Invertebrate | 0.00 ± 0.00 | 0.08 ± 0.04 | 0.04 ± 0.04 | 1.40 | 0.242 |
| Plotosidae | *Plotosus lineatus* | Mobile Invertebrate | 0.00 ± 0.00 | 9.12 ± 9.12 | 0.00 ± 0.00 | 1.15 | 0.385 |
| Pomacanthidae | *Apolemichthys trimaculatus* | Omnivore | 0.07 ± 0.04 | 0.10 ± 0.04 | 0.15 ± 0.07 | 0.94 | 0.395 |
| Pomacanthidae | *Centropyge bicolor* | Mobile Invertebrate | 0.07 ± 0.04 | 0.38 ± 0.12 | 0.02 ± 0.02 | 6.27 | **0.003** |
| Pomacanthidae | *Centropyge bispinosa* | Herbivore | 0.00 ± 0.00 | 0.02 ± 0.02 | 0.00 ± 0.00 | 3.22 | **0.019** |
| Pomacanthidae | *Centropyge fisheri* | Herbivore | 0.00 ± 0.00 | 0.02 ± 0.02 | 0.00 ± 0.00 | 1.47 | 0.155 |
| Pomacanthidae | *Centropyge nox* | Herbivore | 0.07 ± 0.04 | 0.00 ± 0.00 | 0.00 ± 0.00 | 1.87 | 0.137 |
| Pomacanthidae | *Centropyge tibicen* | Herbivore | 0.09 ± 0.06 | 0.08 ± 0.05 | 0.00 ± 0.00 | 0.31 | 0.767 |
| Pomacanthidae | *Centropyge vrolikii* | Herbivore | 0.04 ± 0.03 | 0.00 ± 0.00 | 0.00 ± 0.00 | 2.06 | 0.141 |
| Pomacanthidae | *Chaetodontoplus dimidiatus* | Omnivore | 0.00 ± 0.00 | 0.10 ± 0.05 | 0.29 ± 0.10 | 6.37 | **0.002** |
| Pomacanthidae | *Chaetodontoplus mesoleucus* | Omnivore | 0.22 ± 0.08 | 0.28 ± 0.09 | 0.02 ± 0.02 | 1.90 | 0.155 |
| Pomacanthidae | *Chaetodontoplus poliourus* | Omnivore | 0.06 ± 0.04 | 0.00 ± 0.00 | 0.00 ± 0.00 | 0.30 | 0.862 |
| Pomacanthidae | *Genicanthus lamarck* | Planktivore | 0.00 ± 0.00 | 0.50 ± 0.18 | 0.07 ± 0.07 | 6.00 | **0.002** |
| Pomacanthidae | *Genicanthus melanospilos* | Planktivore | 0.00 ± 0.00 | 0.02 ± 0.02 | 0.00 ± 0.00 | 1.35 | 0.229 |
| Pomacanthidae | *Paracentropyge multifasciata* | Corallivore | 0.02 ± 0.02 | 0.02 ± 0.02 | 0.02 ± 0.02 | 0.50 | 0.684 |
| Pomacanthidae | *Pomacanthus imperator* | Omnivore | 0.11 ± 0.05 | 0.20 ± 0.06 | 0.09 ± 0.04 | 1.23 | 0.299 |
| Pomacanthidae | *Pomacanthus navarchus* | Omnivore | 0.04 ± 0.03 | 0.02 ± 0.02 | 0.00 ± 0.00 | 0.22 | 0.789 |
| Pomacanthidae | *Pomacanthus semicirculatus* | Omnivore | 0.02 ± 0.02 | 0.04 ± 0.04 | 0.00 ± 0.00 | 0.67 | 0.547 |
| Pomacanthidae | *Pomacanthus sexstriatus* | Omnivore | 0.07 ± 0.04 | 0.22 ± 0.09 | 0.09 ± 0.06 | 1.80 | 0.166 |
| Pomacanthidae | *Pomacanthus xanthometopon* | Omnivore | 0.02 ± 0.02 | 0.02 ± 0.02 | 0.00 ± 0.00 | 0.94 | 0.390 |
| Pomacanthidae | *Pygoplites diacanthus* | Omnivore | 0.19 ± 0.07 | 0.24 ± 0.08 | 0.00 ± 0.00 | 2.06 | 0.128 |
| Pomacentridae | *Abudefduf bengalensis* | Omnivore | 0.02 ± 0.02 | 0.00 ± 0.00 | 0.00 ± 0.00 | 1.18 | 0.373 |
| Pomacentridae | *Abudefduf sexfasciatus* | Herbivore & Planktivore | 0.15 ± 0.15 | 0.00 ± 0.00 | 0.00 ± 0.00 | 1.15 | 0.395 |
| Pomacentridae | *Abudefduf vaigiensis* | Mobile Invertebrate | 0.65 ± 0.63 | 0.00 ± 0.00 | 0.00 ± 0.00 | 0.53 | 0.683 |
| Pomacentridae | *Acanthochromis polyacanthus* | Herbivore & Planktivore | 0.43 ± 0.16 | 0.74 ± 0.72 | 0.00 ± 0.00 | 0.71 | 0.574 |
| Pomacentridae | *Amblyglyphidodon aureus* | Herbivore & Planktivore | 0.02 ± 0.02 | 0.14 ± 0.11 | 0.00 ± 0.00 | 2.60 | **0.048** |
| Pomacentridae | *Amblyglyphidodon batunai* | Herbivore & Planktivore | 0.13 ± 0.13 | 0.04 ± 0.03 | 0.00 ± 0.00 | 0.55 | 0.695 |
| Pomacentridae | *Amblyglyphidodon curacao* | Herbivore & Planktivore | 0.35 ± 0.17 | 0.02 ± 0.02 | 0.00 ± 0.00 | 2.64 | 0.063 |
| Pomacentridae | *Amblyglyphidodon leucogaster* | Mobile Invertebrate & Planktivore | 0.35 ± 0.17 | 0.20 ± 0.15 | 0.00 ± 0.00 | 0.44 | 0.656 |
| Pomacentridae | *Amblyglyphidodon ternatensis* | Herbivore & Planktivore | 0.41 ± 0.21 | 0.00 ± 0.00 | 0.00 ± 0.00 | 1.67 | 0.174 |
| Pomacentridae | *Amblypomacentrus breviceps* | Planktivore | 0.04 ± 0.03 | 0.00 ± 0.00 | 0.00 ± 0.00 | 0.32 | 0.788 |
| Pomacentridae | *Amblypomacentrus clarus* | Planktivore | 1.04 ± 1.02 | 0.02 ± 0.02 | 0.02 ± 0.02 | 0.42 | 0.784 |
| Pomacentridae | *Cheiloprion labiatus* | Corallivore | 0.00 ± 0.00 | 0.04 ± 0.04 | 0.00 ± 0.00 | 1.17 | 0.387 |
| Pomacentridae | *Chromis alpha* | Planktivore | 0.00 ± 0.00 | 0.02 ± 0.02 | 0.00 ± 0.00 | 1.21 | 0.343 |
| Pomacentridae | *Chromis amboinensis* | Herbivore & Planktivore | 1.63 ± 1.23 | 0.08 ± 0.05 | 0.00 ± 0.00 | 1.24 | 0.301 |
| Pomacentridae | *Chromis analis* | Planktivore | 0.39 ± 0.24 | 0.00 ± 0.00 | 0.00 ± 0.00 | 1.04 | 0.373 |
| Pomacentridae | *Chromis atripectoralis* | Planktivore | 0.02 ± 0.02 | 0.00 ± 0.00 | 0.00 ± 0.00 | 0.48 | 0.734 |
| Pomacentridae | *Chromis caudalis* | Planktivore | 0.13 ± 0.10 | 0.02 ± 0.02 | 0.28 ± 0.17 | 1.18 | 0.320 |
| Pomacentridae | *Chromis cinerascens* | Herbivore & Planktivore | 0.06 ± 0.06 | 0.00 ± 0.00 | 0.00 ± 0.00 | 0.32 | 0.859 |
| Pomacentridae | *Chromis fumea* | Planktivore | 0.41 ± 0.32 | 0.16 ± 0.08 | 0.00 ± 0.00 | 0.95 | 0.437 |
| Pomacentridae | *Chromis margaritifer* | Herbivore & Planktivore | 0.78 ± 0.23 | 0.08 ± 0.05 | 0.02 ± 0.02 | 7.38 | **0.001** |
| Pomacentridae | *Chromis retrofasciata* | Herbivore & Planktivore | 0.00 ± 0.00 | 0.06 ± 0.06 | 0.00 ± 0.00 | 1.32 | 0.249 |
| Pomacentridae | *Chromis scotochiloptera* | Planktivore | 0.43 ± 0.24 | 0.14 ± 0.10 | 0.00 ± 0.00 | 0.75 | 0.508 |
| Pomacentridae | *Chromis ternatensis* | Planktivore | 3.07 ± 1.53 | 0.06 ± 0.03 | 0.00 ± 0.00 | 1.92 | 0.146 |
| Pomacentridae | *Chromis viridis* | Herbivore & Planktivore | 0.76 ± 0.54 | 0.00 ± 0.00 | 0.00 ± 0.00 | 0.44 | 0.732 |
| Pomacentridae | *Chromis weberi* | Planktivore | 1.09 ± 0.55 | 0.22 ± 0.16 | 0.00 ± 0.00 | 0.48 | 0.659 |
| Pomacentridae | *Chromis xanthochira* | Herbivore & Planktivore | 0.11 ± 0.06 | 0.04 ± 0.04 | 0.00 ± 0.00 | 0.35 | 0.725 |
| Pomacentridae | *Chromis xanthura* | Planktivore | 0.39 ± 0.19 | 0.06 ± 0.03 | 0.00 ± 0.00 | 0.65 | 0.548 |
| Pomacentridae | *Chrysiptera bleekeri* | Herbivore & Planktivore | 0.00 ± 0.00 | 0.32 ± 0.26 | 0.15 ± 0.12 | 1.63 | 0.181 |
| Pomacentridae | *Chrysiptera cyanea* | Herbivore & Planktivore | 1.28 ± 0.70 | 0.02 ± 0.02 | 0.00 ± 0.00 | 3.63 | **0.018** |
| Pomacentridae | *Chrysiptera flavipinnis* | Herbivore & Planktivore | 0.00 ± 0.00 | 0.10 ± 0.10 | 0.07 ± 0.07 | 0.83 | 0.439 |
| Pomacentridae | *Chrysiptera glauca* | Mobile Invertebrate | 0.06 ± 0.04 | 0.00 ± 0.00 | 0.00 ± 0.00 | 0.69 | 0.540 |
| Pomacentridae | *Chrysiptera hemicyanea* | Herbivore & Planktivore | 0.04 ± 0.04 | 0.12 ± 0.10 | 0.00 ± 0.00 | 1.15 | 0.328 |
| Pomacentridae | *Chrysiptera parasema* | Planktivore | 0.09 ± 0.08 | 0.08 ± 0.06 | 0.02 ± 0.02 | 0.08 | 0.940 |
| Pomacentridae | *Chrysiptera pricei* | Herbivore & Planktivore | 0.02 ± 0.02 | 0.00 ± 0.00 | 0.00 ± 0.00 | 2.07 | 0.082 |
| Pomacentridae | *Chrysiptera rex* | Herbivore | 0.15 ± 0.09 | 0.08 ± 0.08 | 0.00 ± 0.00 | 0.43 | 0.671 |
| Pomacentridae | *Chrysiptera rollandi* | Herbivore & Planktivore | 0.11 ± 0.07 | 0.20 ± 0.08 | 0.00 ± 0.00 | 2.92 | 0.055 |
| Pomacentridae | *Chrysiptera springeri* | Herbivore & Planktivore | 0.06 ± 0.06 | 0.02 ± 0.02 | 0.00 ± 0.00 | 0.05 | 0.975 |
| Pomacentridae | *Chrysiptera talboti* | Herbivore & Planktivore | 0.06 ± 0.03 | 0.00 ± 0.00 | 0.00 ± 0.00 | 1.33 | 0.254 |
| Pomacentridae | *Dascyllus aruanus* | Planktivore | 0.15 ± 0.13 | 0.04 ± 0.04 | 0.00 ± 0.00 | 1.15 | 0.332 |
| Pomacentridae | *Dascyllus melanurus* | Omnivore | 0.11 ± 0.09 | 0.00 ± 0.00 | 0.00 ± 0.00 | 0.63 | 0.641 |
| Pomacentridae | *Dascyllus reticulatus* | Planktivore | 0.33 ± 0.14 | 0.04 ± 0.04 | 0.00 ± 0.00 | 2.75 | 0.059 |
| Pomacentridae | *Dascyllus trimaculatus* | Planktivore | 0.31 ± 0.17 | 0.04 ± 0.04 | 0.04 ± 0.04 | 0.58 | 0.615 |
| Pomacentridae | *Dischistodus chrysopoecilus* | Herbivore | 0.06 ± 0.06 | 0.00 ± 0.00 | 0.00 ± 0.00 | 0.16 | 0.956 |
| Pomacentridae | *Dischistodus fasciatus* | Herbivore | 0.02 ± 0.02 | 0.06 ± 0.04 | 0.00 ± 0.00 | 1.16 | 0.327 |
| Pomacentridae | *Dischistodus melanotus* | Herbivore | 0.28 ± 0.16 | 0.02 ± 0.02 | 0.00 ± 0.00 | 1.83 | 0.133 |
| Pomacentridae | *Dischistodus perspicillatus* | Herbivore | 0.22 ± 0.10 | 0.00 ± 0.00 | 0.00 ± 0.00 | 1.77 | 0.163 |
| Pomacentridae | *Dischistodus prosopotaenia* | Mobile Invertebrate & Planktivore | 0.09 ± 0.06 | 0.00 ± 0.00 | 0.00 ± 0.00 | 0.75 | 0.530 |
| Pomacentridae | *Dischistodus pseudochrysopoecilus* | Herbivore | 0.26 ± 0.20 | 0.08 ± 0.06 | 0.00 ± 0.00 | 0.64 | 0.590 |
| Pomacentridae | *Hemiglyphidodon plagiometopon* | Herbivore | 0.11 ± 0.06 | 0.12 ± 0.12 | 0.00 ± 0.00 | 0.37 | 0.760 |
| Pomacentridae | *Lepidozygus tapeinosoma* | Planktivore | 0.00 ± 0.00 | 0.06 ± 0.06 | 0.00 ± 0.00 | 1.53 | 0.156 |
| Pomacentridae | *Neoglyphidodon crossi* | Omnivore | 0.04 ± 0.03 | 0.02 ± 0.02 | 0.00 ± 0.00 | 0.06 | 0.897 |
| Pomacentridae | *Neoglyphidodon melas* | Omnivore | 0.24 ± 0.10 | 0.04 ± 0.04 | 0.00 ± 0.00 | 1.21 | 0.316 |
| Pomacentridae | *Neoglyphidodon nigroris* | Mobile Invertebrate & Planktivore | 0.19 ± 0.09 | 0.28 ± 0.16 | 0.00 ± 0.00 | 1.25 | 0.299 |
| Pomacentridae | *Neoglyphidodon oxyodon* | Herbivore & Planktivore | 0.06 ± 0.03 | 0.10 ± 0.07 | 0.00 ± 0.00 | 1.10 | 0.346 |
| Pomacentridae | *Neoglyphidodon thoracotaeniatus* | Herbivore & Planktivore | 0.00 ± 0.00 | 0.08 ± 0.06 | 0.00 ± 0.00 | 4.04 | **0.012** |
| Pomacentridae | *Neopomacentrus azysron* | Planktivore | 0.07 ± 0.06 | 0.00 ± 0.00 | 0.00 ± 0.00 | 1.29 | 0.272 |
| Pomacentridae | *Neopomacentrus cyanomos* | Planktivore | 0.00 ± 0.00 | 0.02 ± 0.02 | 0.00 ± 0.00 | 1.03 | 0.462 |
| Pomacentridae | *Neopomacentrus filamentosus* | Planktivore | 0.02 ± 0.02 | 0.10 ± 0.07 | 0.00 ± 0.00 | 1.90 | 0.143 |
| Pomacentridae | *Plectroglyphidodon dickii* | Piscivore | 0.13 ± 0.07 | 0.00 ± 0.00 | 0.00 ± 0.00 | 1.88 | 0.135 |
| Pomacentridae | *Plectroglyphidodon lacrymatus* | Omnivore | 0.02 ± 0.02 | 0.02 ± 0.02 | 0.00 ± 0.00 | 0.33 | 0.779 |
| Pomacentridae | *Plectroglyphidodon leucozonus* | Herbivore | 0.04 ± 0.04 | 0.00 ± 0.00 | 0.00 ± 0.00 | 1.44 | 0.175 |
| Pomacentridae | *Pomacentrus adelus* | Herbivore & Planktivore | 0.00 ± 0.00 | 0.08 ± 0.08 | 0.00 ± 0.00 | 1.23 | 0.314 |
| Pomacentridae | *Pomacentrus amboinensis* | Herbivore & Planktivore | 0.02 ± 0.02 | 0.92 ± 0.46 | 0.04 ± 0.04 | 4.33 | **0.007** |
| Pomacentridae | *Pomacentrus auriventris* | Herbivore & Planktivore | 1.83 ± 0.93 | 1.64 ± 0.77 | 0.39 ± 0.39 | 1.24 | 0.292 |
| Pomacentridae | *Pomacentrus bankanensis* | Herbivore & Planktivore | 0.44 ± 0.16 | 0.06 ± 0.04 | 0.00 ± 0.00 | 3.03 | **0.044** |
| Pomacentridae | *Pomacentrus brachialis* | Herbivore & Planktivore | 0.24 ± 0.12 | 0.08 ± 0.05 | 0.00 ± 0.00 | 0.82 | 0.472 |
| Pomacentridae | *Pomacentrus chrysurus* | Omnivore | 0.50 ± 0.24 | 0.06 ± 0.06 | 0.00 ± 0.00 | 0.94 | 0.415 |
| Pomacentridae | *Pomacentrus coelestis* | Mobile Invertebrate & Planktivore | 0.46 ± 0.26 | 0.00 ± 0.00 | 0.00 ± 0.00 | 1.96 | 0.129 |
| Pomacentridae | *Pomacentrus cuneatus* | Herbivore & Planktivore | 0.02 ± 0.02 | 0.00 ± 0.00 | 0.00 ± 0.00 | 0.88 | 0.511 |
| Pomacentridae | *Pomacentrus lepidogenys* | Planktivore | 0.07 ± 0.04 | 0.08 ± 0.08 | 0.00 ± 0.00 | 0.28 | 0.818 |
| Pomacentridae | *Pomacentrus littoralis* | Herbivore & Planktivore | 0.02 ± 0.02 | 0.04 ± 0.03 | 0.00 ± 0.00 | 1.12 | 0.329 |
| Pomacentridae | *Pomacentrus moluccensis* | Omnivore | 1.87 ± 0.57 | 0.00 ± 0.00 | 0.00 ± 0.00 | 4.72 | **0.008** |
| Pomacentridae | *Pomacentrus nagasakiensis* | Herbivore & Planktivore | 0.00 ± 0.00 | 0.00 ± 0.00 | 0.17 ± 0.09 | 3.39 | **0.027** |
| Pomacentridae | *Pomacentrus nigromanus* | Planktivore | 0.15 ± 0.09 | 0.44 ± 0.15 | 0.02 ± 0.02 | 3.86 | **0.021** |
| Pomacentridae | *Pomacentrus nigromarginatus* | Herbivore & Planktivore | 0.00 ± 0.00 | 0.06 ± 0.04 | 0.00 ± 0.00 | 2.18 | 0.093 |
| Pomacentridae | *Pomacentrus opisthostigma* | Herbivore & Planktivore | 0.09 ± 0.06 | 0.00 ± 0.00 | 0.00 ± 0.00 | 1.76 | 0.150 |
| Pomacentridae | *Pomacentrus pavo* | Planktivore | 0.04 ± 0.03 | 0.00 ± 0.00 | 0.00 ± 0.00 | 1.57 | 0.219 |
| Pomacentridae | *Pomacentrus philippinus* | Herbivore & Planktivore | 0.07 ± 0.06 | 0.00 ± 0.00 | 0.00 ± 0.00 | 0.26 | 0.883 |
| Pomacentridae | *Pomacentrus reidi* | Herbivore & Planktivore | 0.20 ± 0.15 | 0.04 ± 0.03 | 0.00 ± 0.00 | 0.23 | 0.869 |
| Pomacentridae | *Pomacentrus simsiang* | Herbivore & Planktivore | 0.13 ± 0.07 | 0.04 ± 0.03 | 0.00 ± 0.00 | 1.07 | 0.368 |
| Pomacentridae | *Pomacentrus taeniometopon* | Herbivore | 0.00 ± 0.00 | 0.00 ± 0.00 | 0.04 ± 0.04 | 0.49 | 0.719 |
| Pomacentridae | *Pomacentrus tripunctatus* | Herbivore | 0.00 ± 0.00 | 0.02 ± 0.02 | 0.00 ± 0.00 | 1.17 | 0.374 |
| Pomacentridae | *Pomacentrus vaiuli* | Mobile Invertebrate & Planktivore | 0.02 ± 0.02 | 0.02 ± 0.02 | 0.00 ± 0.00 | 1.20 | 0.309 |
| Pomacentridae | *Stegastes albifasciatus* | Herbivore | 0.00 ± 0.00 | 0.02 ± 0.02 | 0.00 ± 0.00 | 2.23 | 0.068 |
| Pomacentridae | *Stegastes fasciolatus* | Herbivore | 0.02 ± 0.02 | 0.00 ± 0.00 | 0.00 ± 0.00 | 0.16 | 0.955 |
| Priacanthidae | *Priacanthus hamrur* | Piscivore & Mobile Invertebrate | 0.00 ± 0.00 | 0.02 ± 0.02 | 0.00 ± 0.00 | 1.35 | 0.220 |
| Pseudochromidae | *Pictichromis caitlinae* | Mobile Invertebrate | 0.00 ± 0.00 | 0.04 ± 0.04 | 0.00 ± 0.00 | 1.19 | 0.370 |
| Pseudochromidae | *Pseudochromis fuscus* | Mobile Invertebrate | 0.00 ± 0.00 | 0.02 ± 0.02 | 0.00 ± 0.00 | 1.24 | 0.285 |
| Pseudochromidae | *Pseudochromis jace* | Mobile Invertebrate | 0.00 ± 0.00 | 0.00 ± 0.00 | 0.02 ± 0.02 | 1.99 | 0.098 |
| Pseudochromidae | *Pseudochromis perspicillatus* | Mobile Invertebrate | 0.00 ± 0.00 | 0.00 ± 0.00 | 0.09 ± 0.05 | 3.21 | **0.030** |
| Pseudochromidae | *Pseudochromis pylei* | Mobile Invertebrate | 0.00 ± 0.00 | 0.02 ± 0.02 | 0.00 ± 0.00 | 1.24 | 0.291 |
| Ptereleotridae | *Ptereleotris evides* | Planktivore | 0.00 ± 0.00 | 0.02 ± 0.02 | 0.00 ± 0.00 | 1.24 | 0.290 |
| Ptereleotridae | *Ptereleotris hanae* | Planktivore | 0.04 ± 0.04 | 0.00 ± 0.00 | 0.00 ± 0.00 | 0.49 | 0.726 |
| Ptereleotridae | *Ptereleotris heteroptera* | Planktivore | 0.00 ± 0.00 | 0.02 ± 0.02 | 0.00 ± 0.00 | 1.19 | 0.361 |
| Rachycentridae | *Rachycentron canadum* | Piscivore & Mobile Invertebrate | 0.00 ± 0.00 | 0.02 ± 0.02 | 0.00 ± 0.00 | 0.83 | 0.538 |
| Scombridae | *Grammatorcynus bilineatus* | Piscivore & Mobile Invertebrate | 0.00 ± 0.00 | 0.06 ± 0.03 | 0.09 ± 0.07 | 0.35 | 0.757 |
| Scombridae | *Gymnosarda unicolor* | Piscivore | 0.00 ± 0.00 | 0.10 ± 0.10 | 0.07 ± 0.04 | 0.72 | 0.565 |
| Scombridae | *Scomberomorus commerson* | Piscivore & Mobile Invertebrate | 0.00 ± 0.00 | 0.08 ± 0.06 | 0.01 ± 0.01 | 1.71 | 0.171 |
| Serranidae | *Aethaloperca rogaa* | Piscivore & Mobile Invertebrate | 0.19 ± 0.06 | 0.08 ± 0.04 | 0.00 ± 0.00 | 2.92 | 0.054 |
| Serranidae | *Cephalopholis argus* | Piscivore | 0.31 ± 0.08 | 0.18 ± 0.07 | 0.00 ± 0.00 | 0.14 | 0.871 |
| Serranidae | *Cephalopholis boenak* | Piscivore & Mobile Invertebrate | 0.22 ± 0.06 | 0.04 ± 0.03 | 0.02 ± 0.02 | 3.90 | **0.023** |
| Serranidae | *Cephalopholis cyanostigma* | Piscivore & Mobile Invertebrate | 0.02 ± 0.02 | 0.08 ± 0.05 | 0.00 ± 0.00 | 2.51 | 0.068 |
| Serranidae | *Cephalopholis microprion* | Piscivore & Mobile Invertebrate | 0.04 ± 0.03 | 0.06 ± 0.04 | 0.02 ± 0.02 | 0.30 | 0.771 |
| Serranidae | *Cephalopholis miniata* | Piscivore & Mobile Invertebrate | 0.00 ± 0.00 | 0.02 ± 0.02 | 0.00 ± 0.00 | 1.02 | 0.468 |
| Serranidae | *Cephalopholis polleni* | Piscivore & Mobile Invertebrate | 0.00 ± 0.00 | 0.04 ± 0.04 | 0.00 ± 0.00 | 1.47 | 0.154 |
| Serranidae | *Cephalopholis sexmaculata* | Piscivore & Mobile Invertebrate | 0.00 ± 0.00 | 0.04 ± 0.03 | 0.04 ± 0.03 | 5.19 | **0.004** |
| Serranidae | *Cephalopholis sonnerati* | Piscivore & Mobile Invertebrate | 0.06 ± 0.04 | 0.00 ± 0.00 | 0.00 ± 0.00 | 1.26 | 0.296 |
| Serranidae | *Cephalopholis spiloparaea* | Piscivore & Mobile Invertebrate | 0.00 ± 0.00 | 0.02 ± 0.02 | 0.00 ± 0.00 | 2.23 | 0.073 |
| Serranidae | *Cephalopholis urodeta* | Piscivore & Mobile Invertebrate | 0.11 ± 0.05 | 0.12 ± 0.06 | 0.00 ± 0.00 | 0.73 | 0.506 |
| Serranidae | *Diploprion bifasciatum* | Piscivore & Mobile Invertebrate | 0.15 ± 0.07 | 0.22 ± 0.07 | 0.17 ± 0.09 | 3.74 | **0.025** |
| Serranidae | *Epinephelus areolatus* | Piscivore & Mobile Invertebrate | 0.04 ± 0.04 | 0.00 ± 0.00 | 0.08 ± 0.07 | 0.99 | 0.376 |
| Serranidae | *Epinephelus bontoides* | Piscivore & Mobile Invertebrate | 0.02 ± 0.02 | 0.02 ± 0.02 | 0.00 ± 0.00 | 0.14 | 0.927 |
| Serranidae | *Epinephelus coioides* | Piscivore & Mobile Invertebrate | 0.04 ± 0.03 | 0.08 ± 0.06 | 0.00 ± 0.00 | 0.77 | 0.511 |
| Serranidae | *Epinephelus fasciatus* | Mobile Invertebrate | 0.00 ± 0.00 | 0.04 ± 0.03 | 0.00 ± 0.00 | 2.31 | 0.097 |
| Serranidae | *Epinephelus fuscoguttatus* | Piscivore & Mobile Invertebrate | 0.02 ± 0.02 | 0.04 ± 0.03 | 0.00 ± 0.00 | 1.65 | 0.176 |
| Serranidae | *Epinephelus macrospilos* | Piscivore & Mobile Invertebrate | 0.02 ± 0.02 | 0.00 ± 0.00 | 0.00 ± 0.00 | 0.41 | 0.795 |
| Serranidae | *Epinephelus malabaricus* | Piscivore & Mobile Invertebrate | 0.02 ± 0.02 | 0.02 ± 0.02 | 0.00 ± 0.00 | 0.24 | 0.846 |
| Serranidae | *Epinephelus merra* | Piscivore | 0.06 ± 0.04 | 0.00 ± 0.00 | 0.00 ± 0.00 | 0.38 | 0.798 |
| Serranidae | *Epinephelus polyphekadion* | Piscivore & Mobile Invertebrate | 0.00 ± 0.00 | 0.04 ± 0.03 | 0.00 ± 0.00 | 2.08 | 0.139 |
| Serranidae | *Epinephelus spilotoceps* | Piscivore & Mobile Invertebrate | 0.00 ± 0.00 | 0.02 ± 0.02 | 0.00 ± 0.00 | 2.23 | 0.071 |
| Serranidae | *Gracila albomarginata* | Piscivore | 0.00 ± 0.00 | 0.02 ± 0.02 | 0.00 ± 0.00 | 1.81 | 0.114 |
| Serranidae | *Plectropomus areolatus* | Piscivore | 0.02 ± 0.02 | 0.08 ± 0.05 | 0.00 ± 0.00 | 2.58 | 0.065 |
| Serranidae | *Plectropomus laevis* | Piscivore & Mobile Invertebrate | 0.02 ± 0.02 | 0.00 ± 0.00 | 0.00 ± 0.00 | 0.88 | 0.503 |
| Serranidae | *Plectropomus leopardus* | Piscivore | 0.02 ± 0.02 | 0.02 ± 0.02 | 0.00 ± 0.00 | 0.27 | 0.829 |
| Serranidae | *Plectropomus maculatus* | Piscivore & Mobile Invertebrate | 0.02 ± 0.02 | 0.04 ± 0.04 | 0.00 ± 0.00 | 0.84 | 0.444 |
| Serranidae | *Plectropomus oligacanthus* | Piscivore & Mobile Invertebrate | 0.02 ± 0.02 | 0.18 ± 0.08 | 0.00 ± 0.00 | 3.51 | **0.029** |
| Serranidae | *Pseudanthias dispar* | Planktivore | 0.63 ± 0.63 | 0.00 ± 0.00 | 0.00 ± 0.00 | 0.17 | 0.945 |
| Serranidae | *Pseudanthias huchtii* | Planktivore | 0.02 ± 0.02 | 0.08 ± 0.08 | 0.00 ± 0.00 | 0.51 | 0.718 |
| Serranidae | *Pseudanthias pleurotaenia* | Planktivore | 0.00 ± 0.00 | 0.16 ± 0.16 | 0.00 ± 0.00 | 0.83 | 0.536 |
| Serranidae | *Variola albimarginata* | Piscivore | 0.02 ± 0.02 | 0.20 ± 0.07 | 0.04 ± 0.03 | 5.54 | **0.004** |
| Serranidae | *Variola louti* | Piscivore & Mobile Invertebrate | 0.15 ± 0.06 | 0.20 ± 0.08 | 0.11 ± 0.06 | 0.16 | 0.860 |
| Siganidae | *Siganus argenteus* | Omnivore | 0.41 ± 0.30 | 0.04 ± 0.04 | 0.00 ± 0.00 | 1.64 | 0.157 |
| Siganidae | *Siganus canaliculatus* | Omnivore | 0.02 ± 0.02 | 0.00 ± 0.00 | 0.00 ± 0.00 | 0.78 | 0.565 |
| Siganidae | *Siganus corallinus* | Herbivore & Planktivore | 0.15 ± 0.07 | 0.12 ± 0.06 | 0.00 ± 0.00 | 0.44 | 0.646 |
| Siganidae | *Siganus doliatus* | Herbivore | 0.15 ± 0.08 | 0.00 ± 0.00 | 0.00 ± 0.00 | 2.67 | 0.067 |
| Siganidae | *Siganus guttatus* | Herbivore & Planktivore | 0.00 ± 0.00 | 0.02 ± 0.02 | 0.00 ± 0.00 | 0.60 | 0.643 |
| Siganidae | *Siganus puellus* | Omnivore | 0.24 ± 0.10 | 0.08 ± 0.06 | 0.00 ± 0.00 | 1.56 | 0.224 |
| Siganidae | *Siganus punctatissimus* | Herbivore | 0.07 ± 0.04 | 0.12 ± 0.06 | 0.00 ± 0.00 | 1.99 | 0.133 |
| Siganidae | *Siganus punctatus* | Herbivore | 0.15 ± 0.07 | 0.12 ± 0.09 | 0.00 ± 0.00 | 0.30 | 0.758 |
| Siganidae | *Siganus vulpinus* | Herbivore & Planktivore | 0.24 ± 0.08 | 0.06 ± 0.04 | 0.00 ± 0.00 | 1.88 | 0.152 |
| Sphyraenidae | *Sphyraena barracuda* | Piscivore | 0.00 ± 0.00 | 0.00 ± 0.00 | 0.04 ± 0.04 | 0.99 | 0.468 |
| Syngnathidae | *Choeroichthys brachysoma* | Mobile Invertebrate | 0.00 ± 0.00 | 0.02 ± 0.02 | 0.00 ± 0.00 | 0.73 | 0.579 |
| Syngnathidae | *Dunckerocampus pessuliferus* | Mobile Invertebrate & Planktivore | 0.00 ± 0.00 | 0.02 ± 0.02 | 0.00 ± 0.00 | 2.23 | 0.073 |
| Synodontidae | *Synodus variegatus* | Piscivore | 0.00 ± 0.00 | 0.02 ± 0.02 | 0.00 ± 0.00 | 1.15 | 0.383 |
| Tetraodontidae | *Arothron caeruleopunctatus* | Mobile Invertebrate | 0.02 ± 0.02 | 0.06 ± 0.03 | 0.02 ± 0.02 | 0.45 | 0.634 |
| Tetraodontidae | *Arothron hispidus* | Mobile Invertebrate | 0.04 ± 0.03 | 0.00 ± 0.00 | 0.00 ± 0.00 | 1.43 | 0.253 |
| Tetraodontidae | *Arothron immaculatus* | Piscivore & Mobile Invertebrate | 0.00 ± 0.00 | 0.00 ± 0.00 | 0.02 ± 0.02 | 0.69 | 0.595 |
| Tetraodontidae | *Arothron mappa* | Omnivore | 0.02 ± 0.02 | 0.00 ± 0.00 | 0.04 ± 0.03 | 1.23 | 0.274 |
| Tetraodontidae | *Arothron nigropunctatus* | Mobile Invertebrate | 0.02 ± 0.02 | 0.08 ± 0.05 | 0.00 ± 0.00 | 1.79 | 0.167 |
| Tetraodontidae | *Arothron stellatus* | Mobile Invertebrate | 0.00 ± 0.00 | 0.00 ± 0.00 | 0.02 ± 0.02 | 0.93 | 0.484 |
| Tetraodontidae | *Canthigaster amboinensis* | Omnivore | 0.00 ± 0.00 | 0.04 ± 0.04 | 0.00 ± 0.00 | 0.73 | 0.584 |
| Tetraodontidae | *Canthigaster valentini* | Omnivore | 0.00 ± 0.00 | 0.12 ± 0.07 | 0.00 ± 0.00 | 3.00 | **0.032** |
| Tetraodontidae | *Lagocephalus sceleratus* | Piscivore & Mobile Invertebrate | 0.00 ± 0.00 | 0.01 ± 0.01 | 0.00 ± 0.00 | 1.88 | 0.110 |
| Toxotidae | *Toxotes jaculatrix* | Herbivore | 0.06 ± 0.04 | 0.00 ± 0.00 | 0.00 ± 0.00 | 0.28 | 0.869 |
| Zanclidae | *Zanclus cornutus* | Omnivore | 0.80 ± 0.14 | 0.82 ± 0.15 | 0.28 ± 0.11 | 2.71 | 0.066 |

ESM Table 6. Dufrêne and Legendre indicator species analysis showing fish species indicative of shallow, intermediate, and MCE depth zones. Analysis is based on fish abundance with results showing the Dufrêne and Legendre indicator value (measuring the association between the fish species and the depth group) and the associated permutational p-value (based on 99999 permutations). Only fish species with a p<0.05 association for the depth band are shown.

| **Family / Depth group** | **Species** | **Indicator value** | **P(perm)** | **Trophic Group** |
| --- | --- | --- | --- | --- |
| **Shallow** |  |  |  |  |
| Acanthuridae | *Acanthurus lineatus* | 0.38 | <0.001 | Herbivore |
| Acanthuridae | *Acanthurus maculiceps* | 0.53 | <0.001 | Herbivore |
| Acanthuridae | *Acanthurus nigricauda* | 0.16 | 0.044 | Omnivore |
| Acanthuridae | *Acanthurus xanthopterus* | 0.19 | 0.009 | Planktivore |
| Acanthuridae | *Ctenochaetus striatus* | 0.54 | <0.001 | Planktivore |
| Acanthuridae | *Naso brevirostris* | 0.32 | 0.001 | Herbivore |
| Acanthuridae | *Naso lituratus* | 0.33 | 0.002 | Herbivore_Planktivore |
| Acanthuridae | *Naso unicornis* | 0.18 | 0.044 | Herbivore |
| Acanthuridae | *Zebrasoma scopas* | 0.49 | <0.001 | Herbivore |
| Balistidae | *Balistapus undulatus* | 0.59 | <0.001 | Mobile.Invert |
| Caesionidae | *Caesio lunaris* | 0.28 | 0.017 | Planktivore |
| Caesionidae | *Pterocaesio tile* | 0.26 | 0.004 | Planktivore |
| Carangidae | *Scomberoides lysan* | 0.17 | 0.031 | Piscivore_Mobile.Invert |
| Carcharhinidae | *Carcharhinus melanopterus* | 0.37 | 0.011 | Piscivore_Mobile.Invert |
| Chaetodontidae | *Chaetodon baronessa* | 0.44 | <0.001 | Corallivore |
| Chaetodontidae | *Chaetodon citrinellus* | 0.21 | 0.009 | Omnivore |
| Chaetodontidae | *Chaetodon ephippium* | 0.21 | 0.021 | Omnivore |
| Chaetodontidae | *Chaetodon lineolatus* | 0.26 | 0.004 | Corallivore |
| Chaetodontidae | *Chaetodon lunula* | 0.23 | 0.011 | Mobile.Invert |
| Chaetodontidae | *Chaetodon lunulatus* | 0.48 | <0.001 | Corallivore |
| Chaetodontidae | *Chaetodon rafflesii* | 0.46 | <0.001 | Corallivore |
| Chaetodontidae | *Chaetodon semeion* | 0.25 | 0.003 | Omnivore |
| Chaetodontidae | *Chaetodon speculum* | 0.25 | 0.005 | Corallivore |
| Chaetodontidae | *Chaetodon trifascialis* | 0.46 | <0.001 | Corallivore |
| Chaetodontidae | *Chaetodon vagabundus* | 0.4 | <0.001 | Corallivore |
| Chaetodontidae | *Heniochus varius* | 0.21 | 0.022 | Mobile.Invert |
| Cirrhitidae | *Paracirrhites forsteri* | 0.21 | 0.009 | Piscivore |
| Labridae | *Choerodon anchorago* | 0.31 | 0.001 | Mobile.Invert |
| Labridae | *Diproctacanthus xanthurus* | 0.44 | <0.001 | Corallivore |
| Labridae | *Halichoeres hortulanus* | 0.3 | 0.002 | Mobile.Invert |
| Labridae | *Halichoeres melanurus* | 0.25 | 0.003 | Mobile.Invert |
| Labridae | *Halichoeres podostigma* | 0.17 | 0.031 | Mobile.Invert |
| Labridae | *Hemigymnus melapterus* | 0.37 | <0.001 | Mobile.Invert |
| Labridae | *Hologymnosus annulatus* | 0.24 | 0.014 | Piscivore |
| Labridae | *Labroides dimidiatus* | 0.38 | 0.013 | Piscivore_Mobile.Invert |
| Labridae | *Thalassoma hardwicke* | 0.87 | <0.001 | Mobile.Invert |
| Labridae | *Thalassoma lunare* | 0.75 | <0.001 | Mobile.Invert |
| Labridae (Scarinae) | *Chlorurus bleekeri* | 0.17 | 0.030 | Herbivore |
| Labridae (Scarinae) | *Chlorurus sordidus* | 0.29 | 0.002 | Herbivore |
| Labridae (Scarinae) | *Scarus dimidiatus* | 0.17 | 0.031 | Herbivore |
| Lutjanidae | *Lutjanus bohar* | 0.4 | 0.002 | Mobile.Invert |
| Lutjanidae | *Lutjanus carponotatus* | 0.23 | 0.006 | Piscivore_Mobile.Invert |
| Lutjanidae | *Lutjanus decussatus* | 0.33 | 0.001 | Piscivore_Mobile.Invert |
| Lutjanidae | *Lutjanus fulvus* | 0.25 | 0.003 | Piscivore_Mobile.Invert |
| Lutjanidae | *Lutjanus rivulatus* | 0.2 | 0.033 | Piscivore_Mobile.Invert |
| Lutjanidae | *Lutjanus russellii* | 0.19 | 0.014 | Piscivore_Mobile.Invert |
| Lutjanidae | *Lutjanus semicinctus* | 0.61 | <0.001 | Piscivore_Mobile.Invert |
| Mullidae | *Parupeneus crassilabris* | 0.42 | <0.001 | Mobile.Invert |
| Muraenidae | *Gymnothorax javanicus* | 0.18 | 0.048 | Piscivore_Mobile.Invert |
| Nemipteridae | *Pentapodus trivittatus* | 0.36 | 0.001 | Piscivore_Mobile.Invert |
| Nemipteridae | *Scolopsis bilineata* | 0.45 | <0.001 | Mobile.Invert |
| Nemipteridae | *Scolopsis trilineata* | 0.19 | 0.033 | Piscivore_Mobile.Invert |
| Pomacentridae | *Amblyglyphidodon curacao* | 0.2 | 0.015 | Herbivore_Planktivore |
| Pomacentridae | *Amblyglyphidodon ternatensis* | 0.17 | 0.031 | Herbivore_Planktivore |
| Pomacentridae | *Chromis margaritifer* | 0.45 | <0.001 | Herbivore_Planktivore |
| Pomacentridae | *Chromis ternatensis* | 0.29 | 0.003 | Planktivore |
| Pomacentridae | *Chromis weberi* | 0.27 | 0.006 | Planktivore |
| Pomacentridae | *Chrysiptera cyanea* | 0.25 | 0.004 | Herbivore_Planktivore |
| Pomacentridae | *Dascyllus reticulatus* | 0.18 | 0.022 | Planktivore |
| Pomacentridae | *Dischistodus melanotus* | 0.2 | 0.023 | Herbivore |
| Pomacentridae | *Neoglyphidodon melas* | 0.25 | 0.003 | Omnivore |
| Pomacentridae | *Plectroglyphidodon dickii* | 0.21 | 0.009 | Piscivore |
| Pomacentridae | *Pomacentrus bankanensis* | 0.34 | <0.001 | Herbivore_Planktivore |
| Pomacentridae | *Pomacentrus chrysurus* | 0.22 | 0.008 | Omnivore |
| Pomacentridae | *Pomacentrus coelestis* | 0.21 | 0.009 | Mobile.Invert_Planktivore |
| Pomacentridae | *Pomacentrus moluccensis* | 0.58 | <0.001 | Omnivore |
| Serranidae | *Aethaloperca rogaa* | 0.24 | 0.016 | Piscivore_Mobile.Invert |
| Serranidae | *Cephalopholis argus* | 0.31 | 0.004 | Piscivore |
| Serranidae | *Cephalopholis boenak* | 0.31 | 0.001 | Piscivore_Mobile.Invert |
| Siganidae | *Siganus argenteus* | 0.19 | 0.035 | Omnivore |
| Siganidae | *Siganus doliatus* | 0.17 | 0.032 | Herbivore |
| Siganidae | *Siganus puellus* | 0.21 | 0.008 | Omnivore |
| Siganidae | *Siganus vulpinus* | 0.31 | 0.001 | Herbivore_Planktivore |
| Zanclidae | *Zanclus cornutus* | 0.37 | 0.023 | Omnivore |
| **Intermediate** |  |  |  |  |
| Acanthuridae | *Acanthurus thompsoni* | 0.22 | 0.008 | Planktivore |
| Acanthuridae | *Naso thynnoides* | 0.32 | <0.001 | Omnivore |
| Balistidae | *Odonus niger* | 0.33 | 0.032 | Planktivore |
| Balistidae | *Sufflamen bursa* | 0.39 | 0.001 | Mobile.Invert |
| Caesionidae | *Pterocaesio digramma* | 0.17 | 0.028 | Planktivore |
| Carcharhinidae | *Triaenodon obesus* | 0.18 | 0.042 | Piscivore_Mobile.Invert |
| Chaetodontidae | *Coradion chrysozonus* | 0.22 | 0.033 | Omnivore |
| Chaetodontidae | *Forcipiger flavissimus* | 0.17 | 0.030 | Mobile.Invert |
| Chaetodontidae | *Forcipiger longirostris* | 0.19 | 0.015 | Mobile.Invert |
| Labridae | *Bodianus diana* | 0.19 | 0.009 | Mobile.Invert |
| Labridae | *Bodianus dictynna* | 0.22 | 0.010 | Mobile.Invert |
| Labridae | *Choerodon zosterophorus* | 0.22 | 0.012 | Mobile.Invert |
| Labridae | *Halichoeres chrysus* | 0.39 | <0.001 | Mobile.Invert |
| Labridae | *Halichoeres prosopeion* | 0.33 | 0.002 | Mobile.Invert |
| Lethrinidae | *Monotaxis grandoculis* | 0.22 | 0.020 | Mobile.Invert |
| Lutjanidae | *Macolor macularis* | 0.29 | 0.004 | Piscivore_Mobile.Invert |
| Monacanthidae | *Paraluteres prionurus* | 0.17 | 0.031 | Omnivore |
| Pomacanthidae | *Centropyge bicolor* | 0.3 | 0.002 | Mobile.Invert |
| Pomacanthidae | *Genicanthus lamarck* | 0.29 | 0.001 | Planktivore |
| Pomacentridae | *Pomacentrus amboinensis* | 0.36 | <0.001 | Herbivore_Planktivore |
| Pomacentridae | *Pomacentrus nigromanus* | 0.27 | 0.007 | Planktivore |
| Serranidae | *Plectropomus oligacanthus* | 0.19 | 0.016 | Piscivore_Mobile.Invert |
| Serranidae | *Variola albimarginata* | 0.19 | 0.034 | Piscivore |
| Tetraodontidae | *Canthigaster valentini* | 0.17 | 0.031 | Omnivore |
| **MCE** |  |  |  |  |
| Balistidae | *Abalistes stellatus* | 0.41 | <0.001 | Piscivore_Mobile.Invert |
| Balistidae | *Sufflamen fraenatum* | 0.23 | 0.023 | Mobile.Invert |
| Carangidae | *Carangoides ferdau* | 0.35 | 0.011 | Piscivore_Mobile.Invert |
| Labridae | *Choerodon jordani* | 0.25 | 0.003 | Mobile.Invert |
| Lethrinidae | *Lethrinus amboinensis* | 0.25 | 0.017 | Piscivore_Mobile.Invert |
| Lethrinidae | *Lethrinus lentjan* | 0.17 | 0.031 | Piscivore_Mobile.Invert |
| Malacanthidae | *Hoplolatilus purpureus* | 0.25 | 0.002 | Planktivore |
| Mullidae | *Parupeneus heptacanthus* | 0.2 | 0.027 | Piscivore_Mobile.Invert |
| Nemipteridae | *Pentapodus nagasakiensis* | 0.36 | 0.004 | Mobile.Invert |
| Nemipteridae | *Pentapodus paradiseus* | 0.39 | <0.001 | Piscivore_Mobile.Invert |
| Nemipteridae | *Scolopsis vosmeri* | 0.25 | 0.003 | Mobile.Invert |
| Pomacentridae | *Pomacentrus nagasakiensis* | 0.17 | 0.031 | Herbivore_Planktivore |

ESM Table 7. Linear mixed-effect model selection. All MaxN values were fourth root transformed before model fitting, and all continuous variables (population, benthic, wave, and distance) were scaled and centered prior to model fitting. Population represents 2015 human population within 10 km of the reef site, depth represents the depth grouping (factor with three levels: Shallow, Mid, Deep), benthic represents the first principal component from the principal components analysis of benthic habitat assessment score, wave represents the mean annual wave exposure, distance represents the linear distance to Sorong (the major regional market).

| **Model Number** | **Model update** | **Model structure** | **AICc** | **Conclusion / Additional notes** |
| --- | --- | --- | --- | --- |
| **Regional fisheries species** |  |  |  |  |
| 1 |  | MaxN ~  population * depth + benthic * depth +  wave * depth +  distance * depth +  MPA * depth + (1\|Site) | 261.88 | Initial fully parametrized model |
| 2 |  | MaxN ~ population * depth + benthic * depth +  wave + depth +  distance * depth +  MPA * depth + (1\|Site) | 258.17 | Dropping the wave:depth interaction and comparing to model 1. |
| 3 |  | MaxN ~ population +  benthic * depth +  wave + depth +  distance * depth +  MPA * depth + (1\|Site) | 255.86 | Dropping the population:depth interaction and comparing to model 2. |
| 4 |  | MaxN ~ population +  benthic +  wave + depth +  distance * depth +  MPA * depth + (1\|Site) | 256.03 | Dropping the benthic:depth interaction and comparing to model 3. Retain benthic:depth interaction in final model. |
| 5 |  | MaxN ~ population +  benthic * depth +  wave + depth +  distance +  MPA * depth + (1\|Site) | 251.87 | Dropping the distance:depth interaction and comparing to model 3. |
| 6 |  | MaxN ~ population +  benthic * depth +  wave + depth +  distance +  MPA + (1\|Site) | 248.94 | Dropping the MPA:depth interaction and comparing to model 5. |
| 7 |  | MaxN ~ population +  benthic * depth +  depth +  distance +  MPA + (1\|Site) | 246.74 | Dropping the wave and comparing to model 6. |
| 8 |  | MaxN ~  benthic * depth +  depth +  distance +  MPA + (1\|Site) | 247.95 | Dropping population and comparing to model 7. Retain population in final model. |
| 9 |  | MaxN ~ population +  benthic * depth +  depth +  MPA + (1\|Site) | 249.67 | Dropping distance and comparing to model 7. Retain distance in final model. |
| 10 |  | MaxN ~ population +  benthic * depth +  depth +  (1\|Site) | 247.79 | Dropping MPA and comparing to model 7. Retain MPA in final model. |
| Final model |  | MaxN ~ population +  benthic:depth + benthic +  depth +  distance +  MPA + (1\|Site) | 246.74 | Final model is model 7. |
| **Parrotfish and Surgeonfish** | | | | |
| 1 |  | MaxN ~ depth +  population + population:depth +  benthic + benthic:depth +  wave + wave:depth +  distance + distance:depth +  MPA + MPA:depth + (1\|Site) | 333.86 | Initial fully parametrized model |
| 2 |  | MaxN ~ depth +  population + population:depth +  benthic + benthic:depth +  wave +  distance + distance:depth +  MPA + MPA:depth + (1\|Site) | 329.63 | Dropping the wave:depth interaction and comparing to model 1. |
| 3 |  | MaxN ~ depth +  population +  benthic + benthic:depth +  wave +  distance + distance:depth +  MPA + MPA:depth + (1\|Site) | 328.01 | Dropping the population:depth interaction and comparing to model 2. |
| 4 |  | MaxN ~ depth +  population +  benthic + benthic:depth +  wave +  distance +  MPA + MPA:depth + (1\|Site) | 327.17 | Dropping the distance:depth interaction and comparing to model 3. |
| 5 |  | MaxN ~ depth +  population +  benthic +  wave +  distance +  MPA + MPA:depth + (1\|Site) | 332.58 | Dropping the benthic:depth interaction and comparing to model 4. Retain benthic:depth interaction in final model. |
| 6 |  | MaxN ~ depth +  population +  benthic + benthic:depth +  wave +  distance  MPA + (1\|Site) | 323.36 | Dropping the MPA:depth interaction and comparing to model 4. |
| 7 |  | MaxN ~ depth +  population +  benthic + benthic:depth +  distance  MPA + (1\|Site) | 321.04 | Dropping wave and comparing to model 6. |
| 8 |  | MaxN ~ depth +  benthic + benthic:depth +  distance +  MPA + (1\|Site) | 320.58 | Dropping population and comparing to model 7. |
| 9 |  | MaxN ~ depth +  benthic + benthic:depth +  MPA + (1\|Site) | 320.20 | Dropping distance and comparing to model 8. |
| 10 |  | MaxN ~ depth +  benthic + benthic:depth +  (1\|Site) | 318.00 | Dropping MPA and comparing to model 9. |
| Final model |  | MaxN ~ depth +  benthic + benthic:depth +  (1\|Site) | 318.00 |  |
| **Grouper and Snapper** | | | | |
| 1 |  | MaxN ~ depth +  population + population:depth + benthic + benthic:depth +  wave + wave:depth +  distance + distance:depth +  MPA + MPA:depth + (1\|Site) | 273.03 | Initial fully parametrized model |
| 2 |  | MaxN ~ depth +  population + population:depth + benthic + benthic:depth +  wave +  distance + distance:depth +  MPA + MPA:depth + (1\|Site) | 268.32 | Dropping the wave:depth interaction and comparing to model 1. |
| 3 |  | MaxN ~ depth +  population +  benthic + benthic:depth +  wave +  distance + distance:depth +  MPA + MPA:depth + (1\|Site) | 268.58 | Dropping the population:depth interaction and comparing to model 2. Retain the population:depth interaction. |
| 4 |  | MaxN ~ depth +  population + population:depth +  benthic + benthic:depth +  wave +  distance +  MPA + MPA:depth + (1\|Site) | 269.18 | Dropping the distance:depth interaction and comparing to model 2. Retain distance:depth interaction. |
| 5 |  | MaxN ~ depth +  population + population:depth +  benthic +  wave +  distance + distance:depth +  MPA + MPA:depth + (1\|Site) | 267.95 | Dropping the benthic:depth interaction and comparing to model 2. |
| 6 |  | MaxN ~ depth +  population + population:depth +  benthic +  wave +  distance + distance:depth +  MPA + (1\|Site) | 267.24 | Dropping the MPA:depth interaction and comparing to model 5. |
| 7 |  | MaxN ~ depth +  population + population:depth +  benthic +  distance + distance:depth +  MPA + (1\|Site) | 264.99 | Dropping wave and comparing to model 6. |
| 8 |  | MaxN ~ depth +  population + population:depth +  distance + distance:depth +  MPA + (1\|Site) | 309.06 | Dropping benthic and comparing to model 7. Retain benthic. |
| 9 |  | MaxN ~ depth +  population + population:depth +  benthic +  distance + distance:depth +  (1\|Site) | 266.35 | Dropping MPA and comparing to model 7. Retain MPA. |
| Final model |  | MaxN ~ depth +  population + population:depth +  benthic +  distance + distance:depth +  MPA + (1\|Site) | 264.99 |  |
